# Characterization of Cysteine-rich protein families of *Giardia lamblia* and their role during antigenic variation

**DOI:** 10.1101/2022.03.07.483299

**Authors:** Macarena Rodríguez-Walker, Cecilia R. Molina, Lucas A. Luján, Alicia Saura, Verónica M. Baronetto, Jon Jerlström-Hultqvist, Staffan G. Svärd, Elmer A. Fernández, Hugo D. Luján

## Abstract

*Giardia lamblia* encode several families of cysteine-rich proteins. Among these families are the Variant-specific Surface Proteins (VSPs), which are involved in the process of antigenic variation. In addition to VSPs, other Cys-rich proteins have been described, such as High Cysteine Membrane Proteins (HCMPs), High Cysteine Proteins (HCPs) and Tenascin-like Proteins (TLPs). However, these proteins are less characterized and there is no consensus on their subcellular localization, function, expression, and relationship with the VSPs. Although numerous efforts have been made to determine the distinctive characteristics of VSPs and the number of VSP genes present in the genome of *Giardia*, a clear profile of the VSP repertoire is still lacking. Here, we performed an exhaustive analysis of the Cys-rich families in the recently updated version of the *Giardia* genome, including their organization, characteristic features, evolution and levels of expression, by combining simple pattern searches and predictions with massive sequencing techniques, integrating and reanalyzing as much omics data as possible. We propose a new classification for the Cys-rich protein-encoding genes and pseudogenes that better describes their involvement in the parasite biology and define unique characteristics of the VSPs that include, besides their known features, an Initiator element/Kozak-like sequence, an extended polyadenylation signal and a unique pattern of mutually exclusive transcript accumulation. Our findings also imply that the HCMPs (now named Cys-Rich Membrane Proteins, CRMPs) are upregulated under stress conditions and might protect the parasite during VSP switching. These results contribute to a better understanding of the process of antigenic variation in this pathogen.

**Author Summary:** The most common cause of parasite-induced diarrhea is *Giardia lamblia*. This intestinal parasite causes 180 million cases of symptomatic disease (giardiasis) yearly but also many asymptomatic infections, resulting in that more than 0.5 billion people are currently colonized by the parasite. One key virulence mechanism of *G. lamblia*, resulting in long-term infections and frequent re-infections, is antigenic variation of Variant-specific Surface Proteins (VSPs). The cysteine-rich VSP proteins are encoded by a multi-gene family but until now the number of genes and how they are regulated has been unclear. Here a recently assembled *Giardia* reference genome was analyzed using different bioinformatic analyses and this revealed that there are 136 VSP genes in the *Giardia* genome. The analyses revealed that there are additional cysteine rich proteins in gene families with fewer members that are related to VSPs but with other roles than antigenic variation. A large number of incomplete VSP genes were also identified and they can function as a sequence reservoir for generation of VSP variability. The VSP genes have unique regulatory elements in the upstream and downstream regions, suggesting a role in regulation. Several gene expression data sets were re-analyzed and it showed that one major VSP is expressed per cell. This study is the first to reveal the organization of VSPs in *Giardia* and it will be the basis for further studies of the mechanism of antigenic variation in this important intestinal parasite.

## Introduction

*Giardia lamblia* (syn., *G. intestinalis*, *G. duodenalis*) is a flagellated parasitic protozoan that colonizes the upper small intestine of many vertebrate hosts and is one of the most common causes of diarrhea and intestinal upset in humans worldwide. During its life cycle, *G. lamblia* takes two morphologically distinct forms: the replicative trophozoite and the environmentally resistant cyst. Infection is initiated by ingestion of cysts, followed by excystation and colonization of the upper small intestine by the trophozoites, which are responsible for the clinical manifestations of the disease. When trophozoites descend through the intestine, encystation takes place and cysts are shed in feces to the environment; thus, cysts are ready to initiate new infections when ingested by susceptible hosts. Currently, *G. lamblia* can be divided into eight different genetic groups or assemblages, named from A to H, which mainly segregate according to their host specificity. Parasites within assemblages A and B (and potentially E) infect humans and the most studied *G. lamblia* isolate is WB from assemblage A1 [1].

*Giardia* undergoes antigenic variation, a mechanism that allows the parasite to switch the expression of its variant surface antigens, producing chronic and/or recurrent infections. Antigenic variation in unicellular microorganisms involves three critical requirements: (1) the possession of a large family of homologous genes encoding immunodominant surface antigens; (2) a mechanism that allows the exclusive expression of only one member of such family in individual cells; and (3) a molecular process that changes the expressed antigen for another [2]. In *G. lamblia*, trophozoites are completely covered with a single member of a family of Variant-specific Surface Proteins (VSPs), which are the main antigens recognized by the hosts’ immune systems [3]. The mechanisms proposed to explain antigenic variation in *G. lamblia* are based on the premise that this process must involve the mutually exclusive expression of only one VSP on the surface of individual cells, selected from hundreds of VSP genes encoded in the parasite genome [1,4,5]. Both post-transcriptional and epigenetic mechanisms have been shown to regulate the expression of a unique variant antigen [6,7].

VSPs are type 1a integral membrane proteins. They have a signal peptide (SP), which allows VSPs to enter the secretory pathway, and a conserved C-terminal region comprising a single transmembrane domain (TMD) and a short cytoplasmic tail (CT) of only five amino acids (CRGKA) [8]. The extracellular portion of VSPs is of variable length and rich in cysteine (Cys; C), mainly as CXXC motifs. VSPs are resistant to proteolytic digestion, extreme pH and temperature; besides their role in antigenic variation, VSPs are thought to protect the parasite under the digestive conditions of the upper small intestine [9]. Furthermore, VSPs have been reported to coordinate metals such as iron and zinc [10].

In addition to VSPs, other Cys-rich proteins have been described in *G. lamblia*: High Cysteine Membrane Proteins (HCMPs), High Cysteine Proteins (HCPs), High Cysteine Non-variant Cyst protein (HCNCp) and Tenascin-like Proteins (TLPs) [11–15]. HCMPs are Cys-rich proteins with a predicted TMD near the C-terminal end and a CT different from that of the VSPs [11]. In contrast, HCPs lack the hydrophobic TMD. TLPs have been reported to be secreted by the parasite and been proposed to function as virulence factors [14], and the single HCNCp was described as a novel invariant HCMP specifically overexpressed during cyst formation [11]. However, there is no consensus for all these Cys-rich proteins regarding their subcellular localization, function and relationship among them and, in particular, with the VSPs. Moreover, none of these proteins has been localized with specific antibodies and, in many cases, it is not known whether all are actually transcribed and translated.

VSPs have been the focus of intensive research because they were discovered earlier than the other Cys-rich proteins and are known to be involved in antigenic variation. Although numerous efforts have been made to determine how many VSP genes exist in the genome of *G. lamblia*, at present there is no clear agreement. This characterization was limited for a long time because the first reference genome of this parasite (obtained from the WB isolate) was published more than 10 years ago and was of relatively low quality and coverage [15]. It was not completely sequenced (presence of numerous gaps) and chromosomes were not fully assembled due to limitations in sequencing technologies. Furthermore, the presence of several homologous gene families with highly repetitive sequence elements hindered the generation of a complete and accurate genome assembly for this pathogen. Another disadvantage was that the genome was over-annotated and genes had incorrectly assigned start codons.

From the previous genome version GL2.0, several attempts have been made to characterize and quantify the VSP repertoire, with dissimilar and inconclusive results accentuated by the fact of lacking an established criterion to delineate the features necessary to define a VSP. For example, in an original report of Adam’s group, between 235 and 275 genes were estimated to encode VSPs [16]. In a subsequent analysis, 228 complete VSPs and 75 partial VSPs were identified, estimating a repertoire of approximately 303 VSPs [16]; more recently, Li *et al*. suggested the presence of only 73 putative VSPs [17]. In addition, using monoclonal antibodies (mAb) directed to their ectodomains only a few VSPs were experimentally confirmed on the trophozoite plasma membrane [6]. Consequently, the VSP repertoire in *G. lamblia* has not been clearly characterized.

Recently, a new genome of the WB isolate at 200x coverage and sequenced using a combination of PacBio long-reads and Illumina short-reads (GenBank accession: GCA_000002435.2) combined with structural mapping was published [18]; later, this genome was defined by the NCBI as the new *Giardia intestinalis* Reference Genome (RefSeq accession: GCF_000002435.2). This new genome was assembled into five almost complete chromosomes and, in principle; it was better annotated both structurally and functionally. The haploid genome is 12.6 Mpb in size and contains approximately 5,000 protein-coding genes (CDS) and 320 pseudogenes. In particular, Xu *et al*. found 133 complete VSPs but did not provide specific details about the additional Cys-rich protein families or their relationship with the VSPs and, therefore, with the process of antigenic variation [18]. Since this new Reference Genome (GL2.1) differs substantially from the previous release, most of the transcriptomic and proteomic data produced in the past need to be re-evaluated.

The new version of the *Giardia* genome allows a more accurate analysis of the Cys-rich gene families, their organization, characteristic features, chromosomal localization and potential evolution. Here, different aspects of Cys-rich proteins of *G. lamblia* were exhaustively characterized, combining massive sequencing techniques and integrating as much omics data as possible. This work proposes a new classification and nomenclature for Cys-rich genes and pseudogenes to contribute to a better understanding of the function of these families of proteins.

## Results

### Cysteine-rich protein-coding genes and pseudogenes in the *G. lamblia* genome

Initially, the nucleotide and amino acid sequences of the 4,966 CDS deposited in *G. lamblia* RefSeq database were collected and analyzed. A limit of ≥4% of cysteine content and at least two of any of the CXC, CXXC and/or CXXXC motifs was established for a protein to be Cys-rich [3]. Since the major interest was in the VSPs and related families, a protein BLAST search for all CDS with similarity to previously annotated VSPs (n=133), HCMPs (n=104), HCPs (n=11) or TLPs (n=11) was performed with the aim of finding any other proteins that belonged to these families but that might have been misannotated. Results returned 273 CDS that met these three requirements: similarity (E value<0.0001), high percentage of Cys (≥ 4%) and presence of the characteristic motifs (CXC, CXXC and CXXXC).

Likewise, the nucleotide sequence of the 320 pseudogenes deposited in the RefSeq database was collected and a BLASTx search for all pseudogenes with similarity to previously annotated VSPs, HCMPs, HCPs or TLPs was performed (E value<0.0001). Results returned 217 pseudogenes; all except one have more than 4% of cysteine content and at least two motifs in one of the 3 translated frames. This yielded in total 490 sequences of putative Cys-rich genes and pseudogenes. As described later, some of the previously annotated genes should be considered to be pseudogenes and vice versa; thus, this first selection is mentioned just to describe how the original set of sequences was obtained for subsequent classification and characterization.

A simple decision tree was used for classification of the 490 Cys-rich genes and pseudogenes (**Fig 1**). First, a search for open reading frames (ORFs) from all the collected sequences was performed. If no ORF was found, the sequences were grouped as “pseudogenes type I”. If an ORF was indeed found, the presence of a predicted signal peptide (SP) was determined using Phobius, SignalP and TOPCONS [19–21]. At least two programs had to predict an SP to confirm the presence of this feature. It is worth mentioning that the search for an SP was performed not only in the annotated first methionine (Met; M), but also in up- and downstream Mets. Since the SP is key for protein vesicular trafficking and its deletion may result in loss of function, partial Cys-rich proteins without a proper SP were grouped as “pseudogenes type II” (**Fig 1**). If an SP was predicted, Phobius, TMHMM and TOPCONS were used to infer transmembrane domains (TMDs) [19,21,22], and TMDs were considered present if predicted by at least two of these programs (**Fig 1**). The transmembrane region was extracted using the sequence ranges predicted by Phobius. When the predicted N-terminal SP and the transmembrane regions overlapped, then the prediction returned by Phobius was used to discriminate between the two possibilities, as implemented by the Uniprot Automatic Annotation pipeline. If no TMD was predicted, the sequences were grouped as “Secretory Cys-Rich Proteins (SCRPs)” (**Fig 1**), expecting to find proteomic data demonstrating their translation and not being just other type of pseudogenes. Interestingly, all the genes encoding Cys-rich proteins with TMDs were found to have a single TMD located at the C-terminal end of the proteins (integral membrane proteins type 1a). Then, in the sequences that have a TMD, the presence of the CRGKA cytoplasmic tail (CT) typical of the characterized VSPs was determined. If found, the sequences were grouped as “VSPs”, otherwise, the sequences were grouped as “Cys-Rich Membrane Proteins (CRMPs)” (**Fig 1**). This analysis was done without taking into account previous annotations to avoid any bias in the classification.

**Fig 1.**
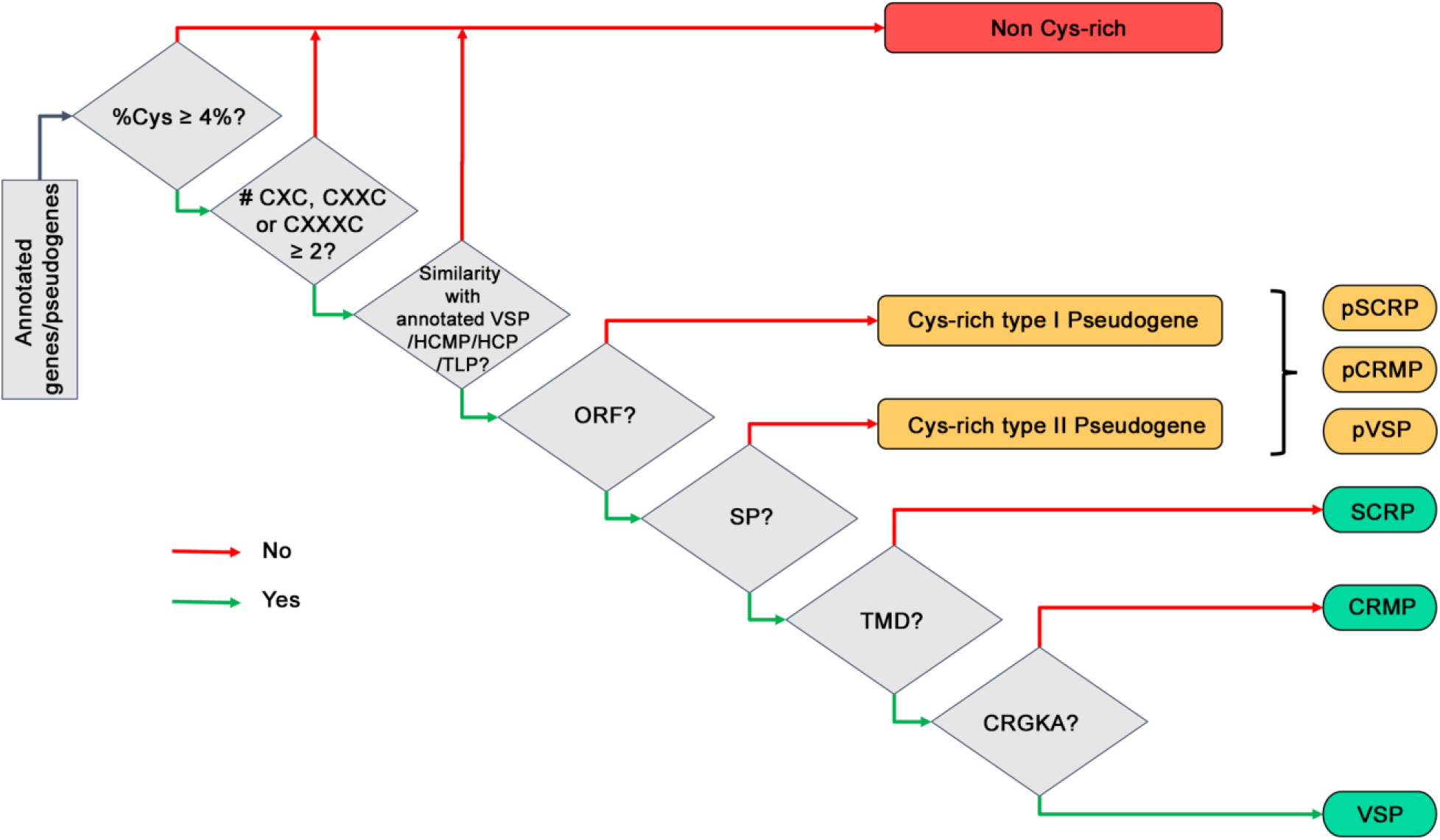
Decision tree used for classification of Cys-rich genes and pseudogenes. Initially, the nucleotide and amino acid sequences of genes and pseudogenes deposited in *G. lamblia* RefSeq database were collected. Three requirements were considered for classifying a gene/pseudogene as Cys-rich: (1) ≥ 4% of cysteine content, (2) at least 2 of any of the CXC, CXXC and/or CXXXC motifs in one of the 3 translated frames, and (3) sequence similarity to previously annotated VSPs, HCMPs, HCPs or TLPs. Next, a search for open reading frames (ORFs) was performed and sequences were grouped as described in the text.

At the same time, all pseudogenes (type I and type II) were divided into pseudo VSP (pVSP), pseudo CRMP (pCRMP) or pseudo SCRP (pSCRP), depending on their similarity to each Cys-rich gene group, using BLASTx against a local database constructed with the VSP, CRMP and SCRP sequences. The subject sequence with the highest score and the lowest E value was taken into account to define whether a pseudogene was a pVSP, a pCRMP or a pSCRP.

**Table 1** shows the number of sequences included in each group after applying the decision tree depicted in **Fig 1** and the assigned names to which they were previously annotated. It was found that not only VSPs, HCMPs, HCPs and TLPs appeared in this table, but other Cys-rich sequences are also included. These ORFs were formerly annotated as neurogenic locus notch proteins, neurogenic locus notch-like proteins, CXC-rich protein, EGF-like domain-containing protein, uncharacterized proteins and hypothetical proteins.

**Table 1.**
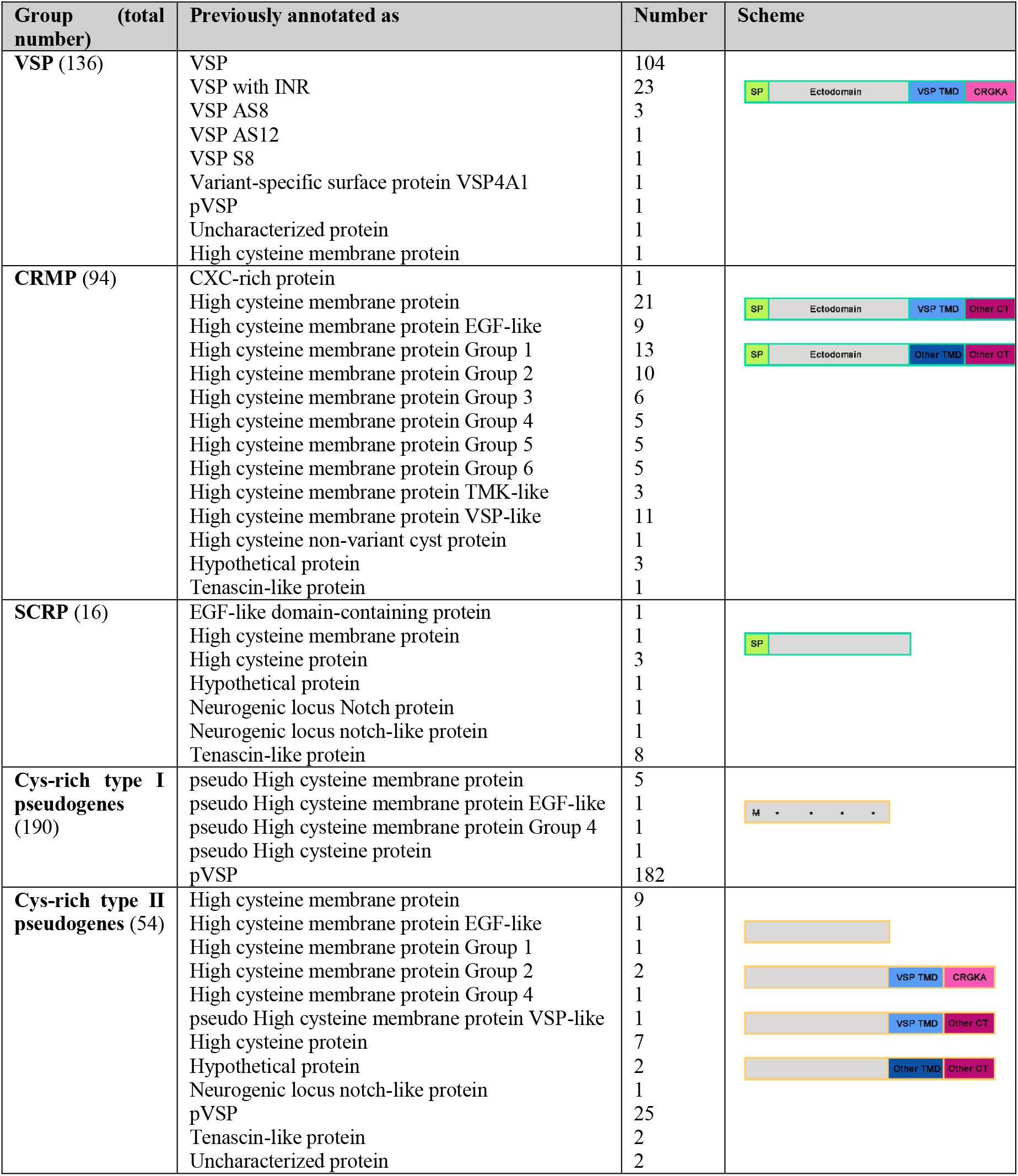
Groups of Cys-rich genes and pseudogenes defined in this work. Each group has its own new name, the total number of sequences and their previous annotation. Diagrams correspond to the detected features. Diagrams with green and orange border represent genes and putative pseudogenes, respectively.

| Group (total number) | Previously annotated as | Number | Scheme |
| --- | --- | --- | --- |
| **VSP (136)** | VSP | 104 |  |
|  | VSP with INR | 23 |  |
|  | VSP AS8 | 3 |  |
|  | VSP AS12 | 1 |  |
|  | VSP S8 | 1 |  |
|  | Variant-specific surface protein VSP4A1 | 1 |  |
|  | pVSP | 1 |  |
|  | Uncharacterized protein | 1 |  |
|  | High cysteine membrane protein | 1 |  |
| **CRMP (94)** | CXC-rich protein | 1 |  |
|  | High cysteine membrane protein | 21 |  |
|  | High cysteine membrane protein EGF-like | 9 |  |
|  | High cysteine membrane protein Group 1 | 13 |  |
|  | High cysteine membrane protein Group 2 | 10 |  |
|  | High cysteine membrane protein Group 3 | 6 |  |
|  | High cysteine membrane protein Group 4 | 5 |  |
|  | High cysteine membrane protein Group 5 | 5 |  |
|  | High cysteine membrane protein Group 6 | 5 |  |
|  | High cysteine membrane protein TMK-like | 3 |  |
|  | High cysteine membrane protein VSP-like | 11 |  |
|  | High cysteine non-variant cyst protein | 1 |  |
|  | Hypothetical protein | 3 |  |
|  | Tenascin-like protein | 1 |  |
| **SCRIP (16)** | EGF-like domain-containing protein | 1 |  |
|  | High cysteine membrane protein | 1 |  |
|  | High cysteine protein | 3 |  |
|  | Hypothetical protein | 1 |  |
|  | Neurogenic locus Notch protein | 1 |  |
|  | Neurogenic locus notch-like protein | 1 |  |
|  | Tenascin-like protein | 8 |  |
| **Cys-rich type I pseudogenes (190)** | pseudo High cysteine membrane protein | 5 |  |
|  | pseudo High cysteine membrane protein EGF-like | 1 |  |
|  | pseudo High cysteine membrane protein Group 4 | 1 |  |
|  | pseudo High cysteine protein | 1 |  |
|  | pVSP | 182 |  |
| **Cys-rich type II pseudogenes (54)** | High cysteine membrane protein | 9 |  |
|  | High cysteine membrane protein EGF-like | 1 |  |
|  | High cysteine membrane protein Group 1 | 1 |  |
|  | High cysteine membrane protein Group 2 | 2 |  |
|  | High cysteine membrane protein Group 4 | 1 |  |
|  | pseudo High cysteine membrane protein VSP-like | 1 |  |
|  | High cysteine protein | 7 |  |
|  | Hypothetical protein | 2 |  |
|  | Neurogenic locus notch-like protein | 1 |  |
|  | pVSP | 25 |  |
|  | Tenascin-like protein | 2 |  |
|  | Uncharacterized protein | 2 |  |

Of all the identified Cys-rich proteins, 55% corresponded to VSPs, 38% to CRMPs and 7% to SCRPs, making up 4.9% of the total protein coding genes and 4.6% of the entire genome. Moreover, Cys-rich protein-coding genes plus pseudogenes contributes to almost 10% of the *Giardia* WB genome.

## 1. General features

### 1.1. Transmembrane domain, cytoplasmic tail and signal peptide

VSPs differ from CRMPs in having a short CRGKA cytoplasmic tail; this is the only distinctive feature of the VSPs [16]. A total of 136 complete VSPs were found, all of them with SP, multiple CXXC motifs, a C-terminal TMD and a CRGKA CT. Of them, 133 were already named VSP in the GL2.1 genome, but the remaining 3 genes were misannotated as High Cysteine Membrane Protein (GL50803_0050332), pVSP (GL50803_0050390) and uncharacterized protein (GL50803_00114674), respectively. In the case of pVSP GL50803_0050390, the SP was found in the second Met in frame.

All 136 proteins have a single, highly conserved hydrophobic TMD of ∼25 amino acids (aa) located near the C-terminal end, according to Phobius prediction [19]. Of the 25 residues, 11 are totally conserved in all VSPs and are positioned at the beginning and at the end of the predicted TMD (**Fig 2**). In addition to the TMD, there is a highly conserved 13 amino acids stretch upstream the predicted TMD that includes a G-XXX-G motif (consensus GGSTNKSS<u>G</u>LST<u>G</u>; see below). On the other hand, SP sequences of the VSPs are not as conserved as the TMD, but still show a degree of similarity (**Fig 2**).

**Fig 2.**
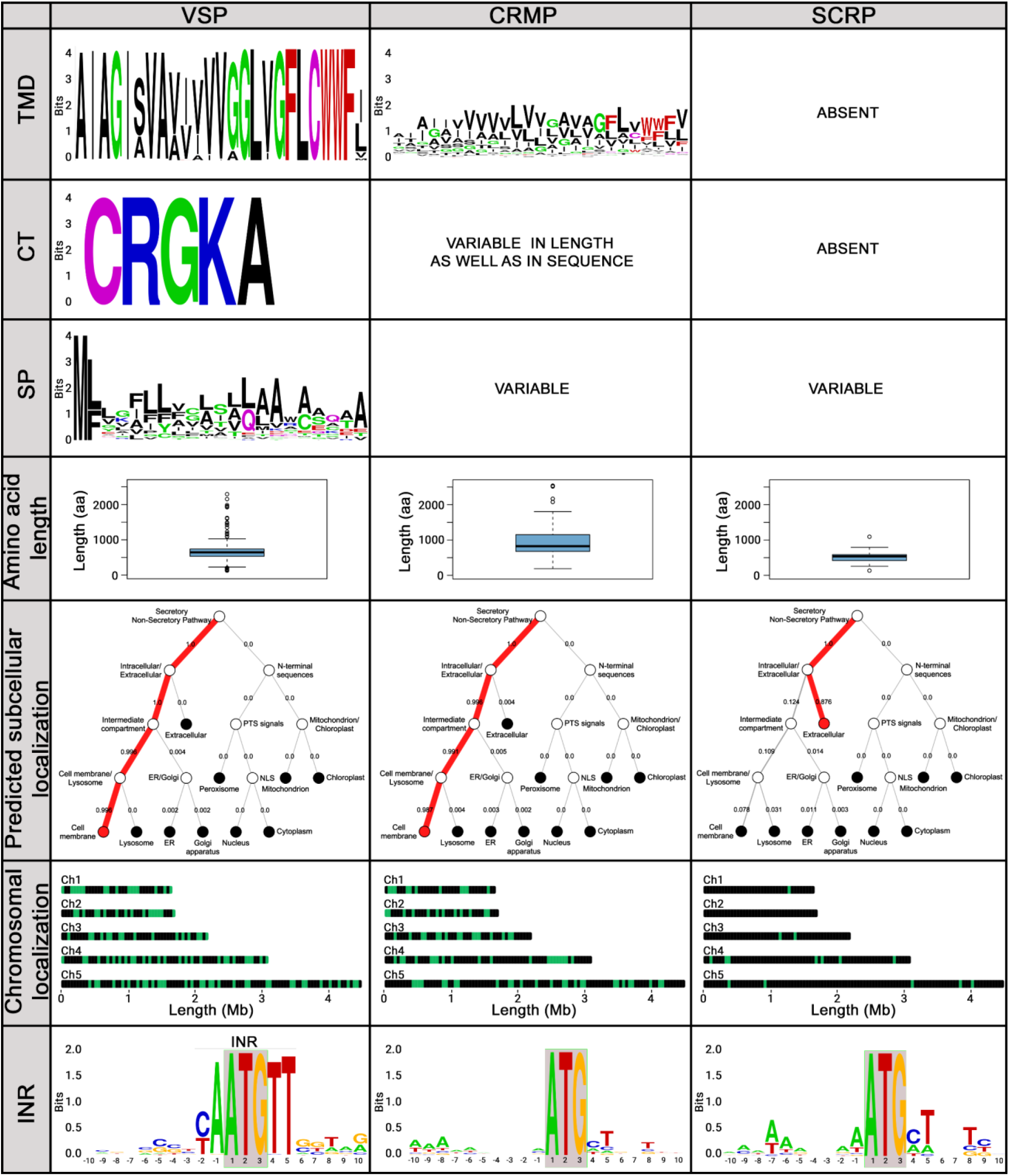
Comparison among VSP, CRMP and SCRP features. Rows (from top to bottom) indicate: Logo of the predicted transmembrane domain; cytoplasmic tail; signal peptide; boxplot of amino acid length; representative diagram outputted by DeepLoc-1.0 subcellular localization predictor; chromosomal distribution of Cys-rich genes (green bars indicate presence of one or more genes in a certain locus); and logo of −10 to +10 nt with respect to the first base of the translational start codon (ATG, boxed).

The CRMP group comprises all Cys-rich proteins with SP, TMD and a CT different from CRGKA. Of the 94 sequences that remained in this group, 88 were previously annotated as HCMPs, 1 as High Cysteine non-variant Cyst Protein (GL50803_0040376), 1 as CXC-rich protein (GL50803_0014225), 1 as Tenascin-like protein (GL50803_0094510), and 3 as hypothetical proteins (GL50803_008733, GL50803_0015450 and GL50803_00113565). All 94 sequences have a predicted single C-terminal TMD. However, while VSPs have a highly conserved TMD, the TMDs of CRMPs is much more variable, both in length (ranging from 20 to 28 aa) and sequence (**Fig 2**), according to Phobius prediction [19]. Notably, when TMHMM [22] was used, a TMD of 23 amino acids was predicted for all 136 VSPs and for all 94 CRMPs. Interestingly, only 2 CRMPs (GL50803_00101589 and GL50803_0010659) have a TMD identical to that of the VSP. Conversely, there are several CRMPs with a TMD completely different from that of the VSPs. Examples of TMDs of CRMPs identical, similar and dissimilar to TMDs of VSPs could be seen in **Table 2**.

**Table 2.**
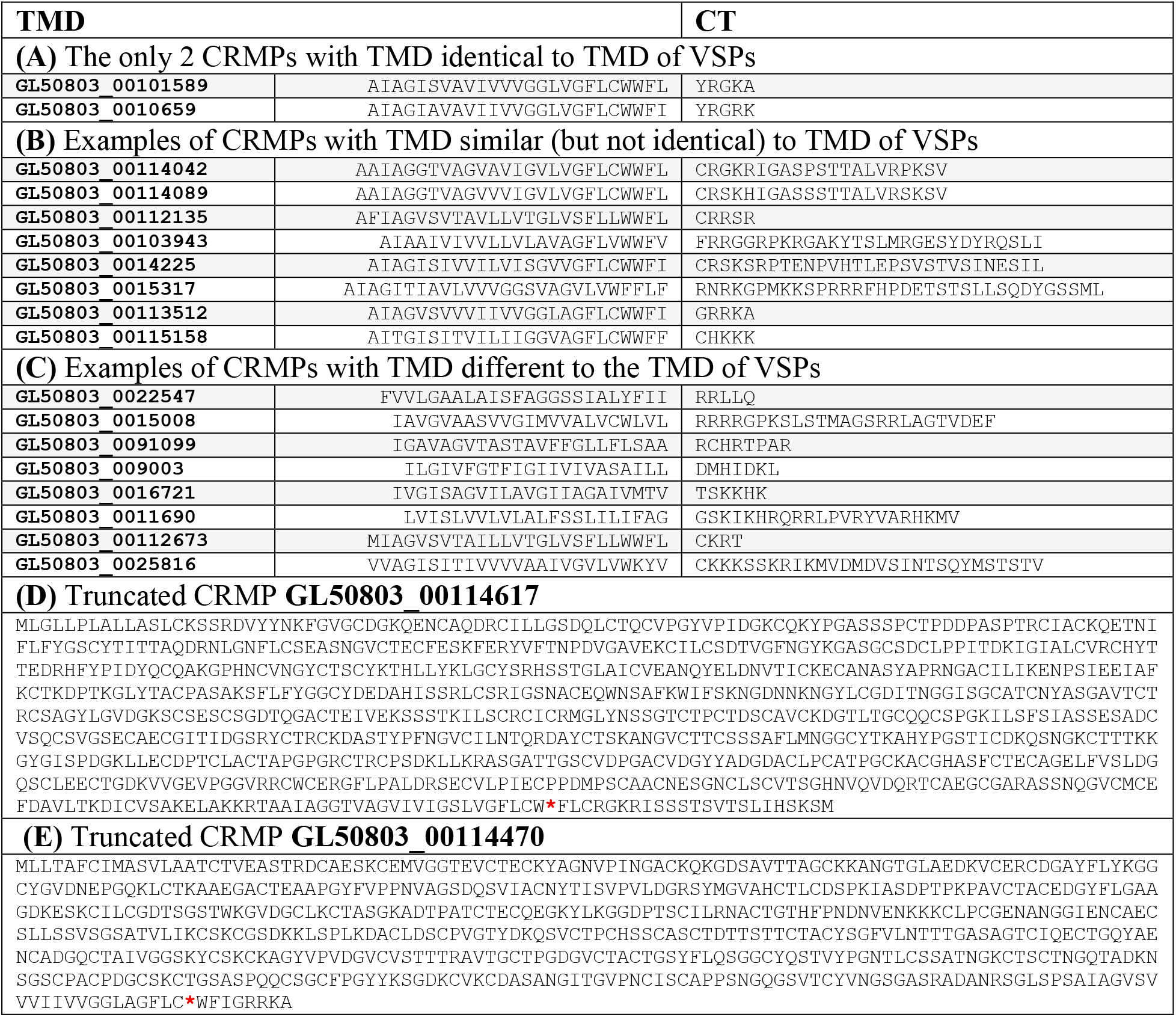
Examples of TMD and CT of CRMP. Examples of CRMP with identical **(A)**, similar **(B)** and different **(C)** TMD compared to VSPs TMD. In all cases, the CT is also shown. **(D)** and **(E)** are examples of CRMPs that have a truncated C-terminal tail after the TMD. These truncated genes, although classified as CRMP, might represent pseudogenes.

In contrast to the characteristic 5-amino acid C-terminal tail of VSPs, those of CRMPs are highly variable, with its length ranging from 1 to 60 aa. Notably, only one CRMP (GL50803_00101589) has an identical TMD to that of the VSPs, but an YRGKA in its CT, instead of the traditional CRGKA. A single nucleotide mutation can change a cysteine (codons UGU and UGC) to a tyrosine (codons UAU and UAC). Due to its characteristics, GL50803_00101589 appears as a true VSP that has suffered a point mutation in its tail (see below). No other CRMP has a C-terminal tail of 5 aa with a mismatch of only one amino acid relative to the CRGKA. Only 3 additional CRMPs have a CT of 5 aa that differs from the CRGKA at two or three positions: CRRSR (GL50803_00112135), GRRKA (GL50803_00113512) and CHKKK (GL50803_00115158). Like GL50803_00101589, GL50803_00113512 looks like a VSP having a mutated CT, which may suggest these two genes could be pseudogenized VSPs. Other CRMPs have completely different tails, both in length and sequence, although all comply with the positive-inside rule, which postulates the preferential occurrence of positively charged residues (K and R) at the cytoplasmic edge of TMDs [23]. The analysis of K and R content both in the ectodomain (ED) and in the CT of Cys-rich membrane proteins showed that, indeed, the CT of these proteins has a higher percentage of positively charged amino acids than the extra cytoplasmic counterpart. VSPs have a CT of 5 aa, of which 1 is R and 1 is K, R and K generate 40% (2 of 5) of positively charged amino acids. In contrast, the ED has between 3 and 11% of K and R. Similar results were observed with the CRMPs; although their CTs are much more variable, they all have R and K content greater than those of the EDs. This is further evidence that the CT of VSPs as well as of CRMPs localized at the cytoplasmic side of cellular membranes.

In addition, some CRMPs had a truncated C-terminal at the end of the TMD (GL50803_00114617 and GL50803_00114470) (**Table 2**). By looking at the respective chromosome, the sequences seem to continue after the stop codon (*FLCRGKRISSSTSVTSLIHSKSM and *WFIGRRKA, respectively), suggesting that a mutation might have occurred, leaving these truncated sequences. Notably, a single nucleotide mutation in the W codon (UGG) can produce two of the three stop codons (UGA and UAG), as observed on those sequences. These truncated genes, although classified as CRMP, might also represent pseudogenes.

Interestingly, of the 104 previously annotated HCMPs, only 88 remained in the CRMP group. Former HCMPs are now distributed across the groups defined here, according to the features. Fourteen of them fell within the pseudogenes Type I group (with ORF without SP), 1 within the SCRP group (GL50803_0029147), 1 is a complete VSP (GL50803_0050332) with a CRGKA tail, and 88 are complete CRMPs.

Among CRMPs, one gene (GL50803_0016454) was predicted by all the used programs to have 2 TMDs, 1 in the N-terminal portion and 1 in the C-terminal end. However, an inherent problem in transmembrane protein topology prediction and signal peptide prediction is the high similarity between the hydrophobic regions of a transmembrane helix and that of a signal peptide, leading to a lack of confidence between the two types of predictions. Therefore, although no SP was predicted for this sequence, it was considered that the N-terminal TMD might be a putative SP instead of a TMD. This was the only confusing case; in all remaining sequences, a clear separation between SP and TMD predictions was observed.

Additionally, all Cys-rich proteins with a predicted SP but lacking a TMD and a CT were named SCRPs. Only 16 proteins fell into this group, 8 of them were formerly named Tenascin-like proteins (GL50803_0010330, GL50803_00113038, GL50803_00114815, GL50803_0014573, GL50803_0016477, GL50803_0016833, GL50803_008687 and GL50803_0095162); 3 High cysteine proteins (GL50803_00115202, GL50803_0012063 and GL50803_009276); 1 High cysteine membrane protein (GL50803_0029147); 1 EGF-like domain-containing protein (GL50803_0037421); 1 hypothetical protein (GL50803_00113268); 1 neurogenic locus notch protein (GL50803_0016322); and 1 neurogenic locus notch-like protein (GL50803_0092495).

The SCRP group is composed mainly of the formerly named Tenascin-like proteins and HCPs. Of all previously annotated HCPs (11 in RefSeq genome), only 3 fell within this group. The remaining ones do not possess a predicted SP, and they were grouped together with pseudogenes type II. Moreover, although the 3 HCPs that remain in this group have a predicted SP, they might also be pseudogenes. This assumption is supported by the fact that for some of them it is possible to find a TMD in a different frame, for example in GL50803_00115202. Therefore, it is likely that most of previously called HCPs actually are pseudogenes (**Table 3**).

**Table 3.**
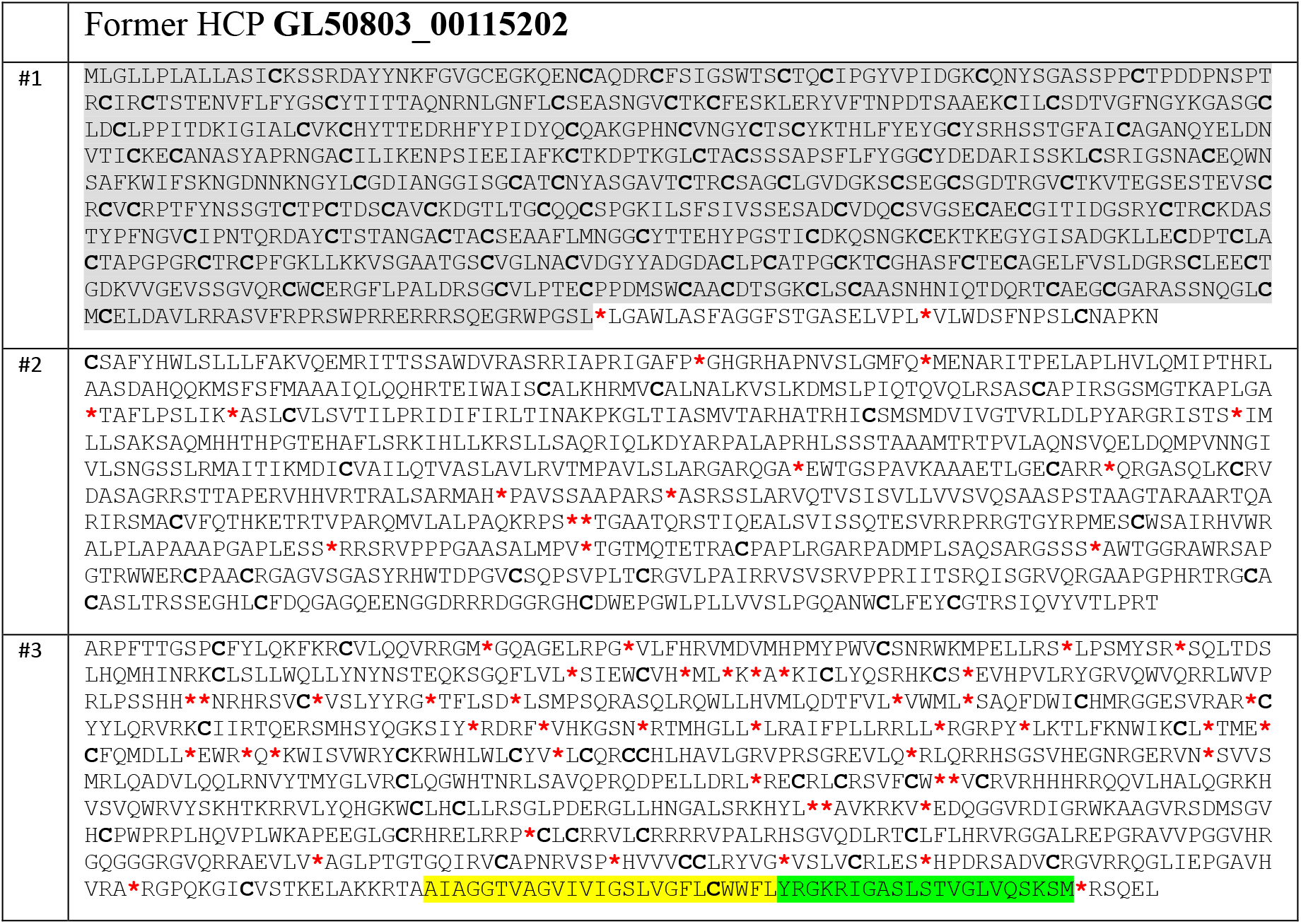
Evidence that former HCP might represent pseudogenes. Example of a previously annotated HCP (GL50803_00115202) in which a TMD and a CT was found in another frame. Highlighted in grey is the annotated ORF in frame 1 (#1). The TMD and CT found in frame 3 (#3) are highlighted in yellow and green, respectively.

On the other hand, of all previously annotated Tenascin-like proteins (11 in the RefSeq genome), 2 fell within the pseudogenes type II group, 8 in this group (SCRP), and only 1 (GL50803_0094510) has a complete ORF with SP and TMD, gene that was already included in the CRMP group. It is possible that true SCRPs have a function in stabilizing VSPs once secreted to the environment or that they may be acting as virulent factors on the host epithelium, as suggested [14].

A detailed information of all the gene and protein IDs, the annotated name, the nucleotide and protein sequence, as well as the predicted TMD and SP for each VSP, CRMP and SCRP are presented in **S1 table**.

### 1.2. Length and predicted molecular weight

The VSPs length ranges from 127 to 2293 aa, with a median of 645 aa (**Fig 2** and **S1 table**). The predicted VSPs mass is between 15 and 260 kDa, with a median equivalent to ∼74 kDa, which is consistent with previously reported estimations indicating that the molecular mass of *G. lamblia* VSPs varies from 20 to 200 kDa [3]. The CRMPs amino acid length ranges from 190 to 2539 aa, with a median of 824.5, equivalent to ∼95 kDa (**Fig 2** and **S1 table**). Lastly, the SCRPs amino acid length ranges from 133 to 1093 aa, with a median of 538 aa, which is expected, since these proteins lack the TMD and CT and, therefore, they should be shorter if derived from VSP or CRMP genes (**Fig 2** and **S1 table**).

### 1.3. Predicted subcellular localization

For this estimation, DeepLoc-1.0 was used [24], which is a eukaryotic protein subcellular localization predictor that applies neural networks trained on Uniprot proteins with experimental evidence of subcellular localization. All 136 VSPs were predicted to localize to the plasma membrane with high confidence (score ranges from 0.9538 to 0.9999, with 1.0 being the maximum score). All 94 CRMPs were also predicted to localize to the plasma membrane, with 96% of them having scores above 0.9. In light of functional and predictive bioinformatics, it is highly likely that SCRPs with SP but without TMD are secreted and, in fact, DeepLoc predicted all these proteins to be extracellular (score ranges between 0.43 and 1.00). A representative example of the output of DeepLoc-1.0 for each group is shown in **Fig 2**.

TMD lengths were found to serve as signatures for subcellular locations in different eukaryotic microorganisms, with plasma membrane TMD lengths being usually longer than those of intracellular organelle membranes [25]. Moreover, the length of the entire TMD determines retention in intracellular membranes or transport to the plasma membrane in yeast [26]. Notably, the TMD of all VSPs, the single CXC-rich membrane protein (GL50803_000014225), and a few CRMPs possess a TMD containing extended GAS_right_ (G-XXX-G-XXX-G) and small(V/L/I)-XXX-small(V/L/I) motifs, which were proposed to facilitate TMD di- and oligomerization [27,28]. The highly conserved TMD of VSPs and the TMD of either a VSP-like CRMP or a CRMP with a highly divergent TMD were analyzed by PREDDIMER [29] and TMHOP [30] to determine their oligomerization capability. Only the Cys-rich membrane proteins having the consensus TMD sequence GAIAGISVAVIVVVGGLVGFLCWWFJ were predicted to form oligomers, as compared to those possessing a different TMD (**Fig 3**). It is known that TMD length, sequences and palmitoylation of the CT are key features of proteins prone to localize into liquid-ordered microdomains of the plasma membrane [32–34]. Therefore, at least the VSPs, and a few CRMPs fulfill those criteria, suggesting that these molecules may be present in lipid raft-like structures.

**Fig 3.**
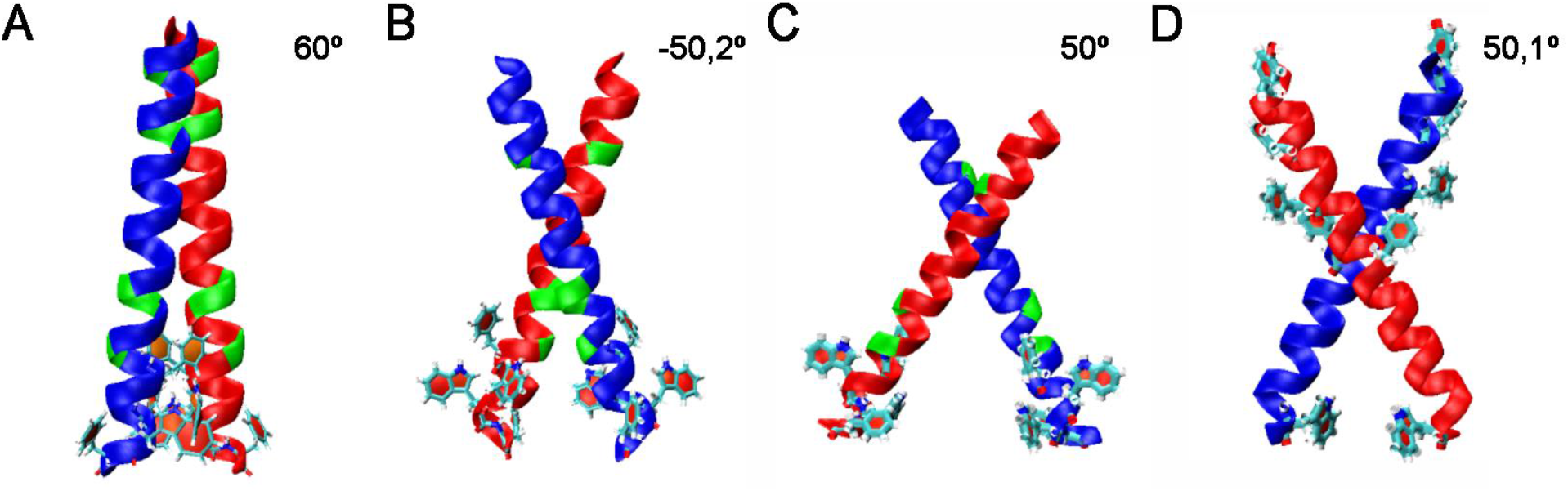
Oligomerization capability of VSP and CRMP TMDs. Predicted oligomers by PREDDIMER. Glycine residues are shown in green. Aromatic amino acid residues are shown as ball and sticks. **(A)** VSP GL50803_00113797 **(B)** CRMP (VSP-like) GL50803_0010659 **(C)** CRMP GL50803_0014225 **(D)** CRMP GL50803_0094510. The characteristic feature of close angle crossing typical of oligomerized TMD structures is only found on the VSP TMD (**A**).

### 1.4. Cysteine codon usage, cysteine content and characteristic motifs

The percentage of total Cys ranges from 7.06% to 15.46% among all VSPs, with a median of 11.89%, which is 5-fold higher than the median cysteine content of all CDS. The percentage of total Cys in CRMPs ranges from 4.93% to 15.63%, with a median of 11.54%, which is similar to the values seen in the VSPs. Interestingly, most VSPs (124 of 136) and most CRMPs (83 of 94) use TGC as the major Cys codon.

VSPs are proteins known to have multiple CXXC motifs distributed all over their extracellular domain, in which two cysteines are separated by two other residues [3,16]. The analysis showed that the minimum number of CXXC motifs found in VSPs is 2 and the maximum is 70, with a median of 26 motifs, and with more than half of the VSPs having between 20 and 30 CXXC motifs (**Fig 4**). Moreover, the number of CXXC motifs always depends, almost linearly, on the length of the VSP extracellular region. Longer VSPs tend to have more CXXC motifs. However, a fraction of the VSPs contain tandem repeats in addition to their common features, which allow them to be even longer than the average length of the VSPs (∼74 kDa). Of the 136 VSPs, only 36 have tandem repeats. Interestingly, similar repeats are found in several VSPs but no duplicated VSPs have repeats. Repeats were analyzed with Tandem Repeats Finder [35], MEME/MAST [36] and RADAR [37], and they were found in the ectodomain of the largest VSPs, with an average of 1174 aa, unlike sequences without repeats, which present an average of 556 aa. Examples of VSPs with tandem repeats are shown in **Table 4** and all VSPs having repeats are listed in **S2 table** (at the nt level) and **S1 fig** (at the aa level). Using this approach, no tandem motifs were found in any CRMPs.

**Fig 4.**
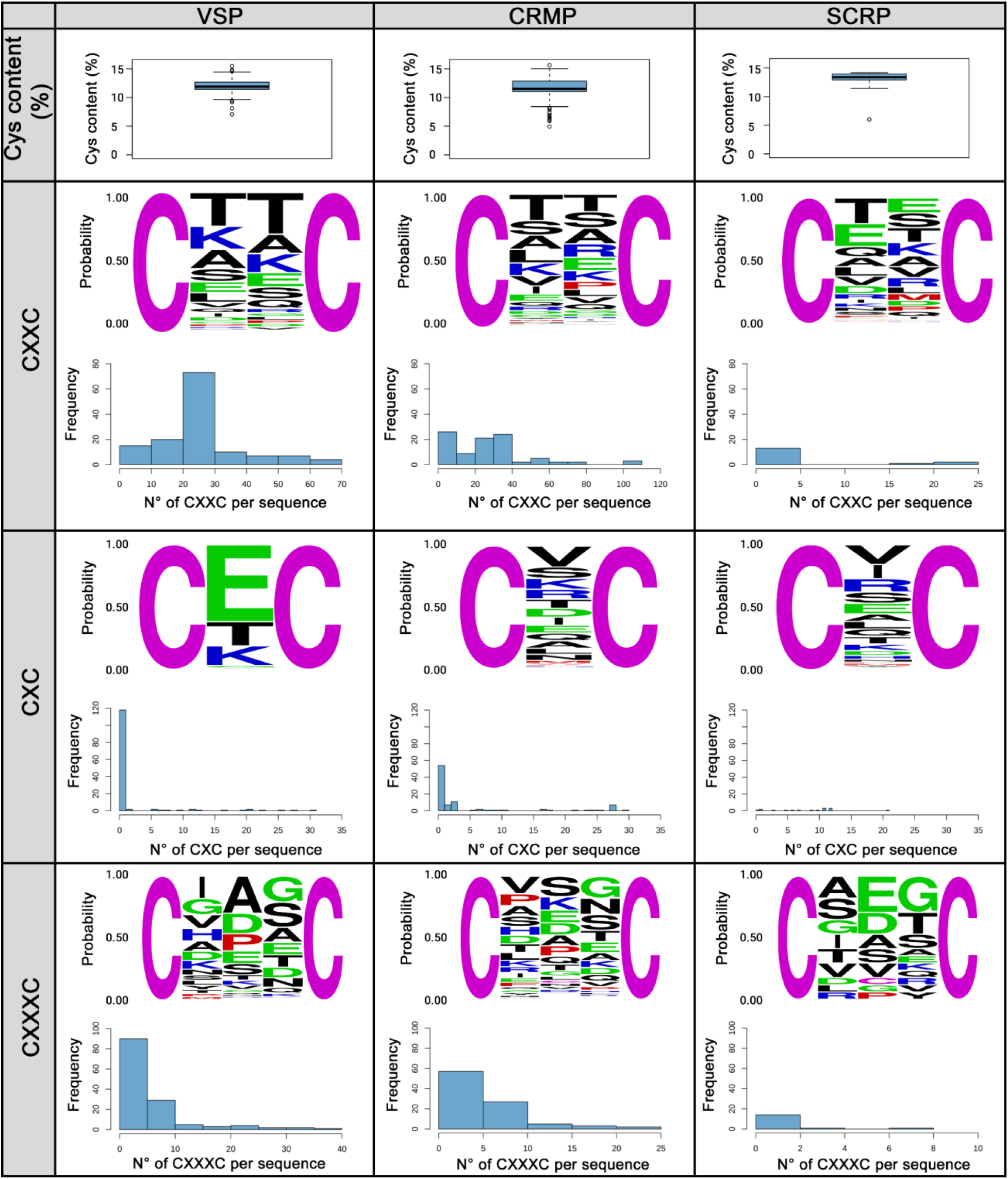
Cysteine content, CXC, CXXC and CXXXC motifs found in Cys-rich proteins. The first row shows cysteine content as the percentage of total amino acids in each group of Cys-rich proteins. The second row shows a logo of CXXC motifs indicating the identity of the second and third amino acids. It also shows a histogram of the number of CXXC motifs per sequence. The third and fourth rows show the same as row two, but regarding CXC and CXXXC motifs, respectively.

**Table 4.**
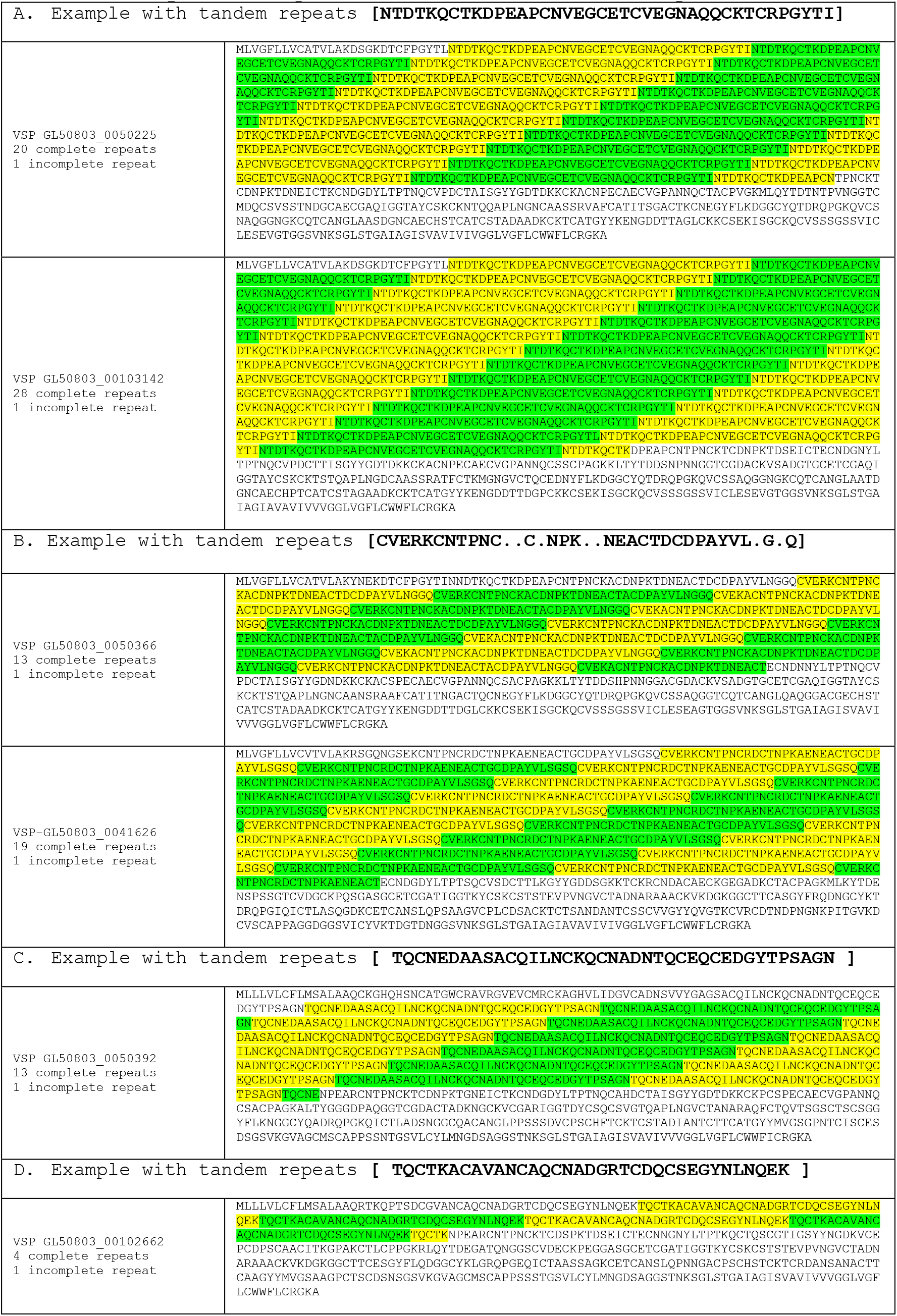
Tandem repeats. Examples of different VSPs with tandem repeats.

Regarding the identity of the X amino acids in the CXXC motifs, it was previously established that X could be any amino acid [8,16]. Here, it was found that in VSPs, X could be any amino acid except C and W. However, there seems to be a preference for T, K, A, S and E in both positions (**Fig 4**). When the combinations of both XX were investigated, of the 174 combinations found, only 6 explained more than 25% of the total frequency (CKTC, CATC, CTAC, CLTC, CETC and CTEC). The CXXC motif is used by many enzymes to catalyze the formation, isomerization, and reduction of disulfide bonds and other redox functions. The sequence of the XX dipeptide located between the cysteines in the active site is important for controlling the redox properties of the protein in which it is found [38,39].

Similar properties have been described for the CXC and CXXXC motifs [38,39]. Consequently, CXC and CXXC motifs in the putative VSPs were identified. Of all VSPs, 90 VSPs do not have any CXC motifs and 28 VSPs only have one. Therefore, only 18 VSPs have more than one CXC motif (**Fig 4**). Amino acid X in CXC is much less variable than in CXXC, being E, T, K, G or V the few amino acids found in the CXC motifs of VSPs (**Fig 4**). As a conclusion, the CXC motif is rare in VSPs, occurring only in one third of the proteins (46 out of 136). Regarding the CXXXC motifs, most VSPs have fewer than 10. There is also a slight tendency of an increase in the number of CXXXC motifs as the length of the protein increases. Furthermore, there are 7 VSPs that do not have any CXXXC motif. Collectively, X in CXXXC can be any amino acid except C and W, just like X in the CXXC motif (**Fig 4**), suggesting that both CXC and CXXXC motifs have evolved from the most ubiquitous CXXC motifs of the VSPs.

All 94 CRMPs have CXXC motifs, but not all of them have CXC (59 of 94) or CXXXC (84 of 94) motifs. This was also seen with the VSPs, where CXXC motifs need to be present, but the other two are not ubiquitous. Regarding the identity of the X amino acid in CXXC and CXXXC motifs, X in CRMP could be any amino acid, in contrast to what was seen in VSPs, in which X could be any amino acid except C and W (**Fig 4**).

Conversely, most SCRPs (12 of 16), especially those already annotated as Tenascin-like proteins, have a marked majority of CXC motifs over CXXC or CXXXC motifs, being a real difference from the VSP group and from most CRMPs. All 16 SCRPs have at least 1 CXXC motif, but not all of them have CXXXC motifs (8 of 16). With respect to the identity of the X amino acid in the major CXC motif, it was found that in SCRP X could be any amino acid except C and P (**Fig 4**).

The number of CXC, CXXC and CXXXC motifs, as well as the total number of cysteines of each Cys-rich protein can be found in **S1 table**.

Additionally, since CXXC motifs have been reported to coordinate Fe^2+^ or Zn^2+^ in a purified VSP [10], the DiANNA software [40] was used to determine not only the capability of all these motifs to bind these metals but also the possibility to form disulfide bonds. Results showed that all VSPs and most CRMPs were predicted to bind those metals and to form intra- and inter disulfide bonds (**S3 table**). It is likely that these characteristics favor the formation of the dense coat observed on the trophozoite [41], which is capable to resist the action of proteases present in the upper small intestine [42].

Besides the aforementioned highly conserved TMD and CT, the SP sequence and multiple CXXC motifs, MEME/MAST [36] analyses detected additional features in VSPs such as the tetrapeptide GGCY (**S1 table**). Previous works described that a GGCY motif is characteristic of VSPs and that mutations in GGCY result in loss of VSP surface localization [3,16]. However, only 117 out of the 136 complete VSPs have at least a single GGCY motif: 49 VSPs have one motif, 64 VSPs have two motifs, and 4 VSPs have more than two motifs (4, 6, 8 and 13 motifs each). VSPs that have a single GGCY motif (e.g., GL50803_00113797) have it located towards the C-terminus of the ectodomain, upstream the TMD. VSPs that have two GGCY motifs (e.g. GL50803_00112208) have an extra motif located closer to the N-terminus. Moreover, the GGCY motif is also found in 47 CRMPs: 1 CRMP has one motif, 9 CRMPs have two motifs and, interestingly, 26 CRMPs have more than two motifs. For example, CRMP GL50803_00114180 has 12 GGCY motifs. Regarding SCRPs, only 3 of 16 have at least one GGCY motif (**S1 table**). Remarkably, previously annotated TLPs that remained in the SCRP group do not possess this motif. Therefore, no clear function for this particular motif can be assumed.

Since no structure of the Cys-rich proteins from *Giardia* has been reported, all Cys-rich protein sequences were then analyzed with SMART (Simple Modular Architecture Research Tool) [43] with the aim of searching for conserved protein sequence motifs/domains. Regarding VSPs, no characteristic motif was found to be present in all 136 sequences. In fact, most sequences (74/136) had no matches in SMART database. The remaining sequences presented at least one of the following motifs: Furin-like repeats, EGF domain, and EGF-like domain or Pfam VSP domain, all of which which contain CXXC motifs. Interestingly, the Pfam VSP domain (PF03302) was found only in few VSPs (19/136), indicating that this domain cannot be used as a characteristic of this family of variable surface antigens. All the motifs found were located in different regions of the sequences with no particular order, sometimes near the N-terminus, sometimes near the C-terminus, with their multiplicity also varying, from one to multiple domains per sequence and with some of them overlapping. Similar results were found in the analysis of the 94 CRMPs. Twenty-seven of them had no matches in SMART database. In the remaining sequences, it was frequent to find Furin-like repeats, EGF domain, EGF-like domain or Pfam VSP domain, as already seen in VSP sequences. Exactly the same results were found for SCRPs, i.e. no particular motif was found present in any of the 16 sequences, 3 of them had no matches in SMART database, and the remaining 13 sequences presented at least one of the Furin-like repeats, EGF domain, EGF-like domain or Pfam VSP domain. The fact that all Cys-rich families present the same motifs, without a strong differentiation between them, is a clear indication that all the sequences do have a common origin (**S2 fig**).

In addition, the lack of similar protein structures with high cysteine content in public repositories makes their *in silico* modeling highly difficult. Numerous efforts were made to predict the structure of the VSPs using various approaches, with dissimilar results. VSP and CRMP structures were predicted by Phyre^2^ [44] and AlphaFold [45] **(Fig 5)**. Predictions made by Phyre^2^ show that both VSPs and CRMP share characteristics with solved domains of the EGF receptor, highlighted in green in **Fig 5A** and **B**. On the other hand, AlphaFold shows many short beta sheets in the ectodomain of both families of Cys-rich proteins (**Fig 5C** and **D**). Notably, similar structures are found in the Amyloid-β precursor protein (APP). APP is a lipophilic metal chelator with metal-reducing activity that binds copper, zinc and iron [46,47]. In general, the predicted structures show that these proteins possess a highly disordered organization, making almost impossible to predict any structural characteristics for the ectodomains of VSPs and CRMP. Nevertheless, both groups of proteins share similar features and properties when expressed on the plasma membrane of the trophozoites.

**Fig 5.**
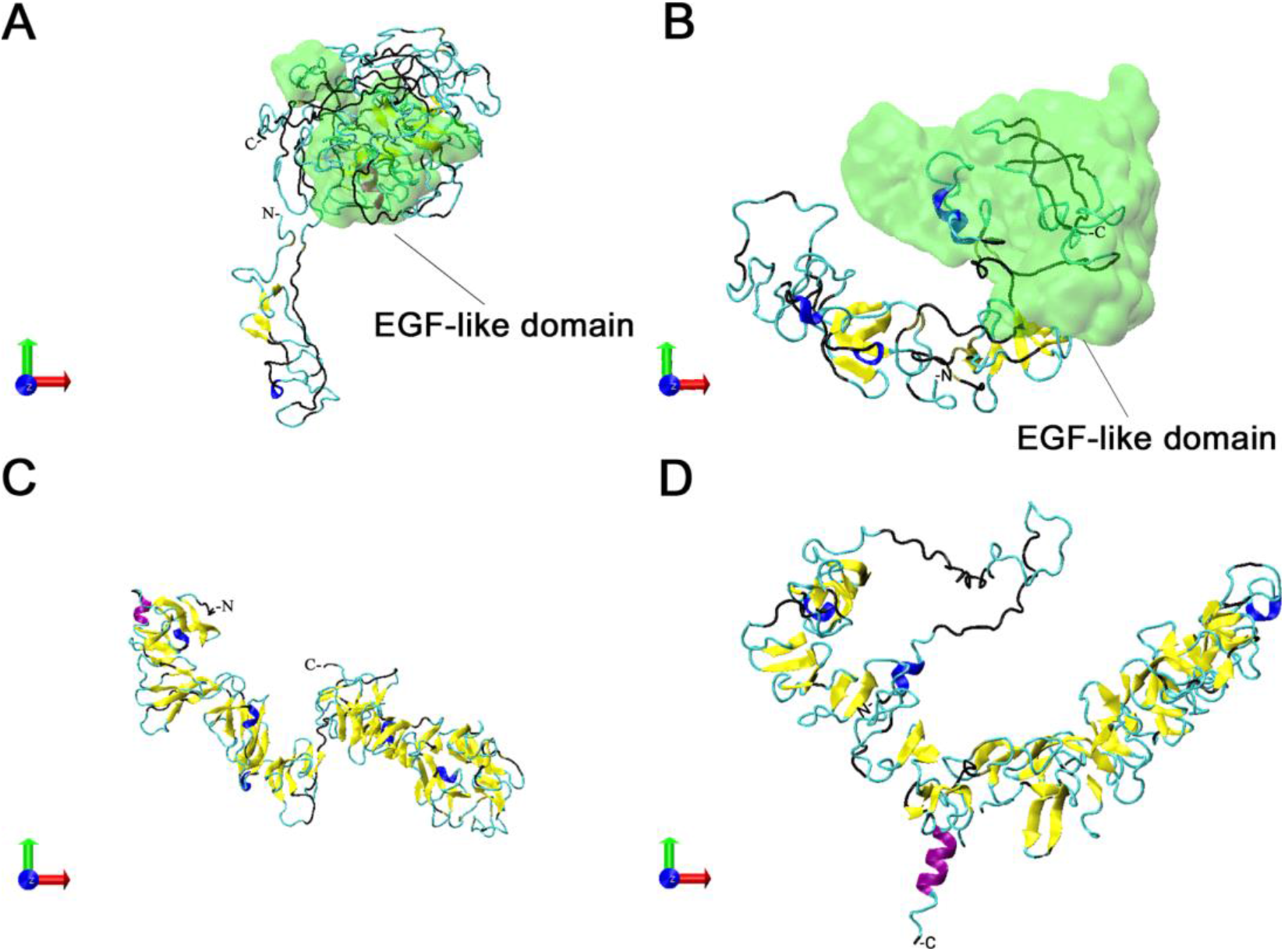
Predicted ectodomain structure of VSP and CRMP. (**A**) VSP GL50803_00113797 predicted by Phyre^2^. EGF-like domains are found in both VSPs and CRMP (highlighted in green). (**B**) CRMP GL50803_0016454 predicted by Phyre^2^. (**C**) VSP GL50803_00113797 predicted by AlphaFold. (**D**) CRMP GL50803_0016454 predicted by AlphaFold.

To determine the relationships between the amino acid sequences of VSPs and CRMP, a phylogenetic analysis was performed using MAFFT [48] and RAxML [49] and the output tree assembled using iTOL [50]. Notably, VSPs grouped together, whereas CRMP segregated into two well-separated groups, one including more divergent CRMP and the other including CRMPs that share some similarity with the VSPs (**S3 fig**). Few CRMP similar to VSPs grouped together with the VSPs, as expected (**S3 fig**).

### 1.5. Chromosomal location and duplicated pairs

All VSP and CRMP genes are distributed in noncontiguous locations on all five chromosomes (**Fig 2**) and the longer the chromosome, the more Cys-rich genes it encodes. There are 20, 20, 21, 32 and 43 VSP genes in chromosomes 1 (NC_051856.1), 2 (NC_051857.1), 3 (NC_051858.1), 4 (NC_051859.1) and 5 (NC_051860.1), respectively, and 11, 15, 15, 20 and 33 CRMP genes in chromosomes 1, 2, 3, 4 and 5, respectively. No complete VSP or CRMP coding genes were found in the 30 still unplaced scaffolds.

As mentioned by Xu *et al.* [18], regions around VSP genes tend to be gene-poor. To re-evaluate that finding, the length of all intergenic regions in the *G. lamblia* genome was calculated; results yielded a median and mean length of 79 and 392.5 nt, respectively. In contrast, the length of VSP intergenic regions (i.e., the length in nt from a VSP gene to its two proximal genes/pseudogenes) were longer, with a median and mean of 1280.0 and 2131.1 nt, respectively. The length of CRMP intergenic regions (median and mean of 363.0 and 812.7 nt, respectively) tend to be larger than the average *G. lamblia* intergenic regions, but shorter than that of VSPs.

Although the genome is not perfectly annotated and intergenic distances may change as the annotation is refined and new genes are discovered or discarded, it can be stated that most of the *G. lamblia* intergenic regions are short, but those of the Cys-rich, and especially those of the VSPs, tend to be longer. Moreover, many VSP genes have contiguous hypothetic genes that do not have transcription or translation according to the data deposited in the GiardiaDB (www.GiardiaDB.org) and to our own analyses (see below). Therefore, the intergenic regions could be longer than appreciated, since those hypothetical genes might not be actual genes.

It was earlier observed, using a sliding window GC content analysis, that the VSP containing chromosomal regions had a higher GC content than the rest of the genome [51]. A re-analysis of 5’-upstream regions of the Cys-rich genes showed that the average GC content in the 2000 bp 5’-upstream region is 53.9% for VSPs and 50.6% for CRMPs; whereas if the 200 bp 5’-upstream region is analyzed, the average GC level for VSP genes is 50.6% and 43.6% for CRMPs. This should be compared to the average GC content of all protein-coding genes, which is 45.2%. Thus, the “promoter” region of these genes seemed larger and with a GC content higher for Cys-rich genes than of any other *Giardia* gene.

Although it was known for a long time that some VSP genes were duplicated, as reported for the VSP1267 (GL50803_00112208) recognized by mAb 5C1 [52], they have been questioned as assembly artifacts in the earlier GL2.0 genome. Recently, Xu *et al*. found that among the 133 VSPs identified by them in the new genome assembly, 38 were in duplicated pairs [18]. Here, 40 VSPs were confirmed to be in duplicated pairs (in nt sequence as well as in aa sequence, thus representing 20 pairs). Thirteen of them have tail-to-tail orientation (→ ←), and 7 have head-to-head orientation (← →). Such pairs were found in all chromosomes but non-identical VSP genes were found in different chromosomes (**S1 table**). Among them, 11 pairs were well annotated, with one member of the pair having a “d” added after the locus tag prefix GL50803_00 (e.g., GL50803_00112208 and GL50803_00d112208; VSP1267) However, the remaining 9 pairs were not explicitly indicated as duplicated, since their locus tags were different (e.g., GL50803_00117472 and GL50803_00117473 or GL50803_00137714 and GL50803_0011470). Therefore, to improve their annotation, these nine pairs should be renamed with a single locus tag and a “d” should be added to one member of the pair. Furthermore, two VSPs identified as GL50803_00101765 and GL50803_00d101765 were wrongly annotated as duplicates, since they have slightly different sequences. Thus, they should be renamed accordingly. All this information about VSP duplicated pairs is summarized in **S1 table** and **S4 table**.

As seen for the VSP genes, there are also duplicated CRMP pairs, but not as many as the VSPs (**S1 table** and **S4 table**). Only 6 CRMPs were identified in duplicated pairs, both in nt sequence and in aa sequence. Of the 3 pairs, 1 has head-to-tail orientation (← ←), and 2 have head-to-head orientation (← →). Furthermore, it was also noted that two CRMPs identified as GL50803_0026981 and GL50803_00d26981 are annotated as duplicates, but they have different sequences and they should be renamed accordingly. In fact, GL50803_00d26981 is identical to the protein annotated as GL50803_00112673; therefore, they are both the duplicated pair.

SCRP genes are also distributed in noncontiguous locations on chromosomes (**Fig 2**), but there are no duplicated pairs. Moreover, there is no difference in the length of intergenic regions of SCRP compared to all *G. lamblia* intergenic regions (median and mean of 158.0 and 300.6 nt versus 79 and 392.5 nt, respectively).

Remarkably, apart from the VSP and CRMP genes, there are not many identical duplicated genes in the *G. lamblia* genome. Only 2.4% of the 4,966 protein coding genes are duplicated (most of which are Cys-rich proteins, hypothetical proteins located near duplicated VSPs, and some members of the multifamily of NEK kinases and Ankyrin repeat-containing proteins).

Therefore, the analysis of the local genomic context turns crucial to understand the expansion of the VSP gene family (**Fig 6**). It was observed that some VSP genes are partially duplicated and that they have similar sequences in their intergenic region. Interestingly, many VSP pairs contain an annotated pseudogene between the members of the pair. To search for genes/pseudogenes that might have been skipped in the previous annotation, the intergenic regions of the duplicated pairs were analyzed using BLASTn. Results showed that when pairs are oriented tail-to-tail, it was frequent to find pseudogenes in between, and when pairs are oriented head-to-head, it was frequent to find retrotransposon remnant sequences between them (**S4 table**).

**Fig 6.**
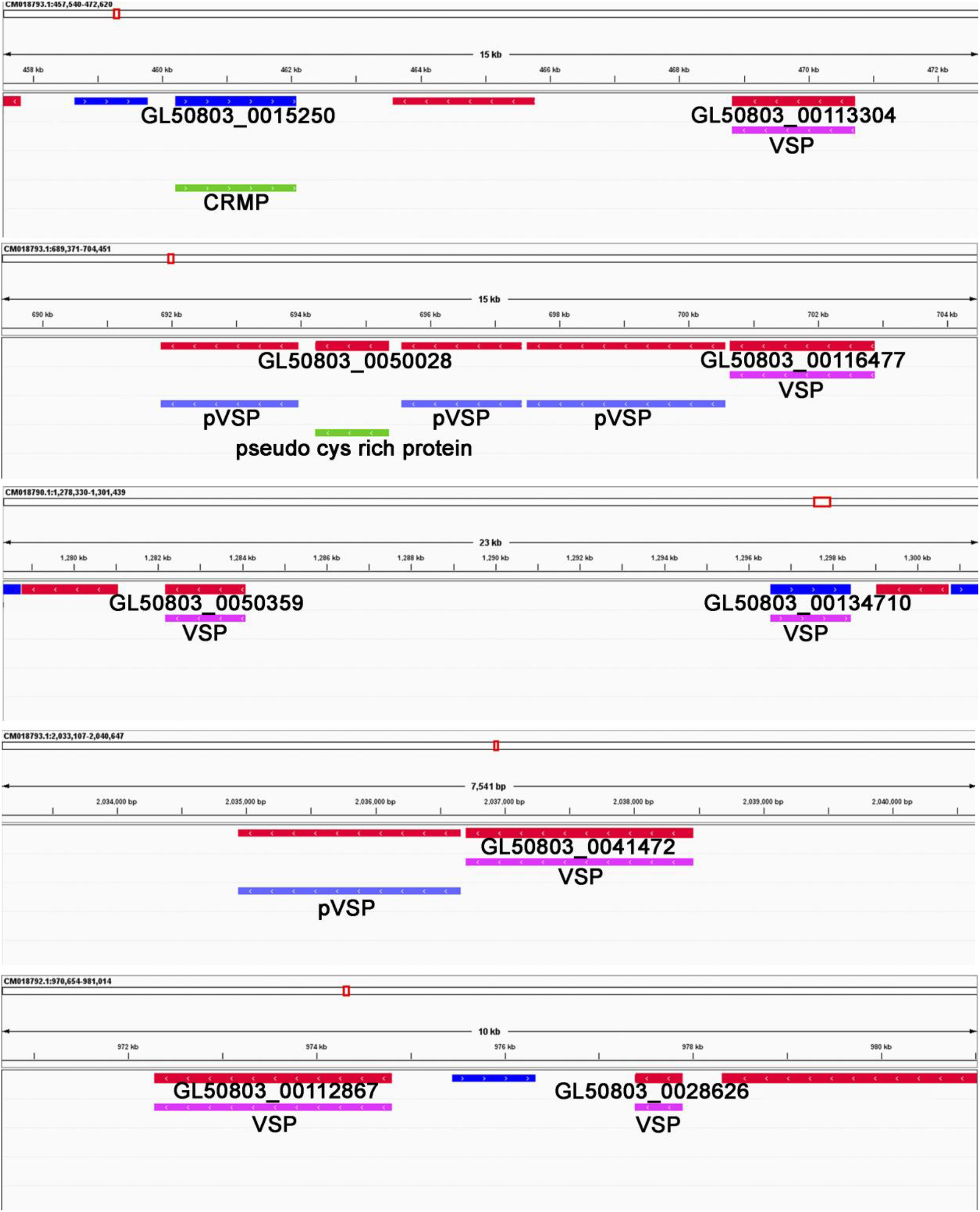
Genome context of VSP and CRMP genes and pseudogenes. IGV snapshots of the genome context of selected genes. Note the presence of pseudogenes and duplicated (identical and divergent VSP sequences).

For example, VSP GL50803_00112113 and VSP GL50803_0050375 are positioned head to head, separated by 5,172 nt, and their intergenic region is highly homologous to gene-free subtelomeric regions. The region between the duplicated VSP genes GL50803_00119706 and GL50803_00119707, which are located tail to tail, has 3,301 nt, with the presence of an annotated non-transcribed pseudogene that BLASTed with two retrotransposon relics of the non-LTR LINE type [53]. Moreover, almost identical intergenic sequences are located between duplicated VSP genes and have similarity with other VSP genes, such as the intergenic region between GL50803_00d112208 and GL50803_00112208 (head to head, 2,648 nt in length) that is highly conserved with the region located between the duplicated VSPs GL50803_00d115797 and GL50803_00115797 (head to head, 2,649 nt apart). On the other hand, there are genomic regions where VSP, CRMP and pseudo Cys-rich genes are located. For example, VSP GL50803_00113024 and VSP GL50803_0050225 have an identity of 98% in 85% of their sequence, but they are located head to head and separated by 21,223 nt, sequence that have two duplicated pVSPs and two duplicated CRMPs (**S4 table**).

Remarkably, when the 7000-nt long 5’-upstream region of VSP1267 (GL50803_00112208 and GL50803_d00112208) was BLASTed against the entire *Giardia* genome, similar sequences were found in most of the 5-upstream regions of head-to-head duplicated VSP/VSP and VSP/CRMP pairs. CENSOR analysis of these regions identified sequences derived from the LTR retrotransposons BEL, Gypsy and Mariner/Tc1 [54]. Since relics of transposable elements sequences were also found in the flanking regions of tail-to-tail duplicated VSP genes, these observations strongly suggest that VSP genes appeared earlier in evolution. Then, they expanded by initial duplications driven by transposition which, after suffering subsequent mutations, deletions and insertions (especially of tandem repeats), generated not only different VSPs, as previously reported [16], but also multiple CRMPs and SCRPs. Although maintaining the basic modules responsible for the protective capability of these molecules at the trophozoite surface [9], the sequence variability of components of the dense surface coat of *Giardia* also provided the parasite with the ability to evade the host immune responses [5].

### 1.6. 5’-UTR and Initiator element (Inr)

The alignment of the 200 nt upstream of all VSP genes shows the absence of any known sequence or consensus motif common to all of them, as previously observed [16,55]. However, there seems to be groups of VSP genes that share highly similar sequences as can be seen in the tree and alignment shown in **S4 fig**. Notably, all VSP found in pairs also share the same 5’ upstream sequence and even some VSPs not forming pairs share their upstream sequences as well.

For the appropriate expression of an eukaryotic gene, most organisms use two ubiquitous sequence motifs, the TATA box and the Initiator element (Inr), both required for binding of the TFIID, a subunit of the RNA polymerase II preinitiation complex [56–58]. The TATA box is the binding site of the TATA-binding protein (TBP) and other transcription factors. A representative TATA box (TATAWAW) is not present in the 5’ upstream sequences of the *Giardia* genes, including the VSP genes (**S4A fig**), coincident with the absence of a typical TBP in the genome of the parasite (only a TBP-like protein GL50803_001721 was found). A tree showing these groupings are shown in **S4B fig**.

On the other hand, the Inr is the simplest promoter that can endorse transcription initiation without a TATA box. The Inr is a 17-nt sequence found at the transcription start site of eukaryotic genes [57]. It was described that some VSP genes have the DNA consensus sequence PyA<u>ATG</u>TT, where Py is C or T and <u>ATG</u> is the start codon, at the beginning of the coding region. Transient transfection studies showed that this consensus sequence is required for efficient expression of firefly luciferase from a VSP promoter region [16]. Therefore, this consensus region surrounding the start codon was named initiator element (Inr). In that study, using the previous version of *G. lamblia* genome, it was found that only few of the VSP genes contained this Inr sequence. Consequently, if Inr was found, VSPs were annotated as “VSP with INR”; otherwise, they were annotated just as “VSP”. Of the 136 VSP identified in this work, only 23 are annotated as VSPs with INR. However, in a first scan, 132 of the 136 VSP genes were found to contain this sequence motif. A closer inspection of the remaining sequences revealed that a pair of duplicated VSPs (GL50803_00137722 and GL50803_00137723) and VSP GL50803_0050390 (formerly named pVSP) contained that sequence in the next in frame ATG codon, indicating that these three sequences might have a wrong annotated start codon and the second Met is actually the first. It is worth mentioning that these shorter proteins also have a predicted signal peptide. Making this correction, all but one (135 out of 136) are now VSPs with Inr (**Fig 2**). Among them, 83 genes (61.5%) contained the sequence CAatgTT, while 52 VSP genes (38.5%) contained the sequence TA<u>ATG</u>TT. The VSP without the Inr element in any ATG codon is GL50803_0014043 and has the sequence (T)AA<u>ATG</u>TT. Interestingly, in this VSP gene, the nucleotide in position +3 is a T, and it is possible that the extra A in +1 or +2 position could result from a single nucleotide insertion, disrupting the consensus Inr sequence. With this finding, the hypothesis that the Inr sequence is required for efficient expression of a subset of VSPs genes is refused, and all VSPs should be annotated as VSP with Inr, or simply VSP. Interestingly, the Inr element seems to be almost exclusive to VSP genes, since it was not found in other coding genes, with only very few exceptions (**S5 table**). The Inr was not found in any of the remaining Cys-rich protein-coding genes, except in one CRMP (GL50803_00101589), whose sequence is TA<u>ATG</u>TT. This particular CRMP is the one possessing the YRGKA CT and a TMD identical to that of VSPs. This strongly suggests that this gene was a VSP gene that has undergone the mutation C→Y. No characteristic motifs were found in the 5’-upstream regions of CRMPs and SCRPs, except for a slight preference for AT-rich nucleotides, as also seen in all *Giardia* protein-coding genes.

Additionally, due to the shortness of the Inr sequence as well as its localization, it may also function as a Kozak-like sequence [58] almost exclusive to *Giardia* VSPs. Therefore, it is not clear whether the PyA<u>ATG</u>TT sequence works as an initiator element for DNA transcription or as a Kozak-like sequence in VSP mRNAs that may facilitate the highly efficient translation of members of this gene family (**S1 table)**.

It was previously reported that the 50 nt upstream region of housekeeping genes suffices to drive the expression of firefly luciferase [59], proteins of interest [60–63] as well as VSPs [6] by constitutively expressing those genes in plasmids also encoding either neomycin or puromycin resistance genes with the 5’- and 3’-regions of housekeeping genes [63]. To evaluate the influence of 5’ upstream region on the expression of a VSP gene, plasmids with different 100 nt-upstream regions were designed. By expressing VSP417 (GL50803_00113797) with its wild type “promoter” that includes the Inr (WT) in trophozoites expressing VSP1267 (GL50803_00112808), a very low percentage of cells expressing VSP417 was observed (<5%), even the fact that those cells were under the pressure of puromycin. However, by expressing the same VSP gene with the 5’-upstream region of either ornithine carbamoyl transferase (OCT) or α-tubulin (TUB), which have no Inr and are highly transcribed and translated (see below), more than 50% of the cell population expressed VSP417, as determined by IFA using mAb 7C2 (**Fig 7A**). These results suggest two different possibilities: (a) the transcription factors that drive the expression of housekeeping genes may drive transcription of any *Giardia* gene or (b) the VSP’s 5’-upstream region including the Inr is not “active” in trophozoites already expressing another VSP, which is likely due to epigenetic control of mutually exclusive expression of VSP genes [7].

**Fig. 7.**
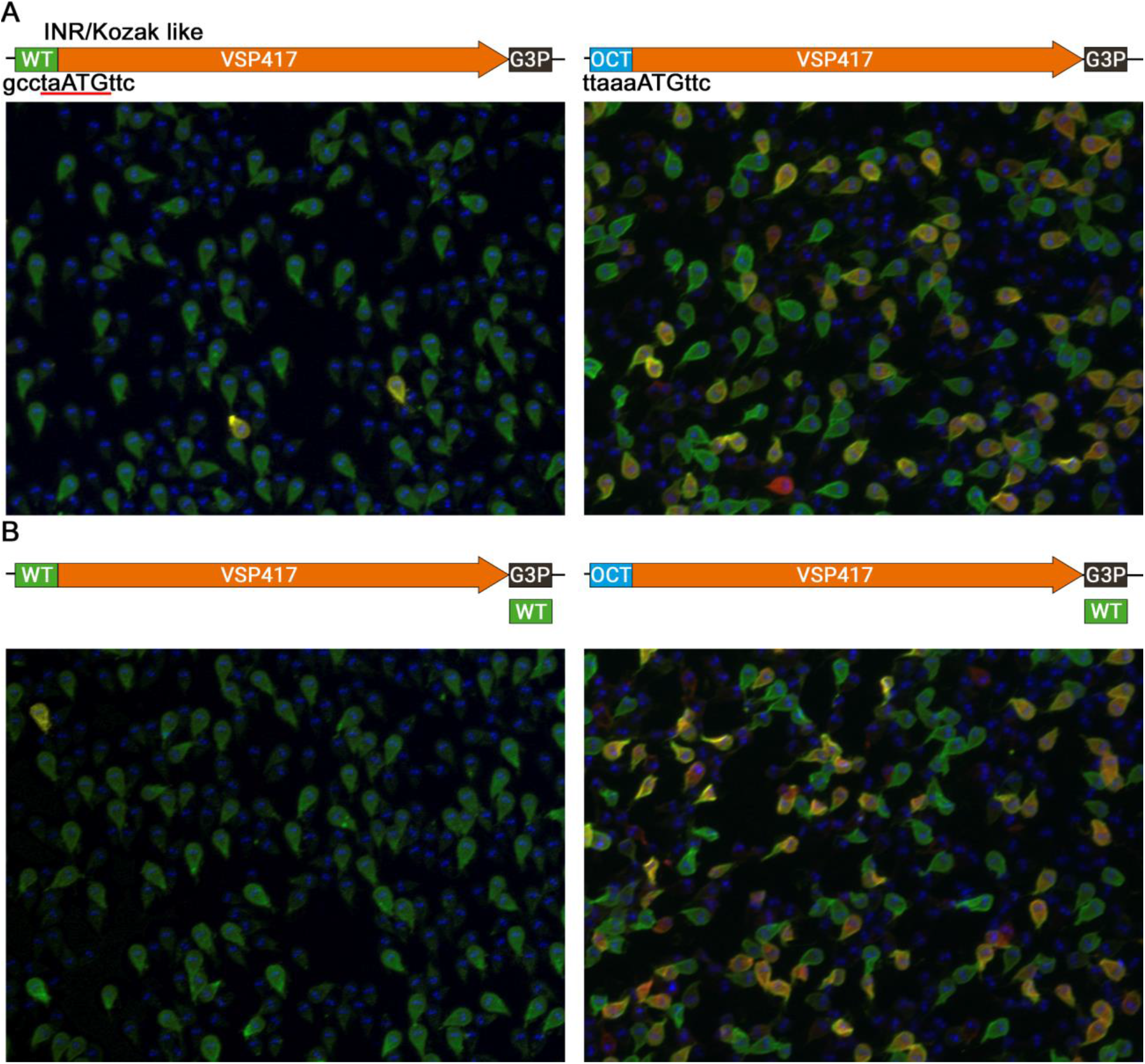
Constitutive expression of VSPs under the control of different 5’ and 3’ regions. The pCIDIEpac vector was modified to replace include the ORF of VSP417 and different “promoter” region and the 3’-downstream region of the VSP or housekeeping genes. The 100 nt-long 5’-upstream region of the either VSP417 or OCT (**A, top panels**) and the 100-nt long 3’-downtream regions of either VSP417 or G3P (**B, bottom panels**) were used. Then, these plasmids encoding the VSP417 were transfected on VSP1267 expressing trophozoites. Green cells correspond to VSP417, red cells to VSP1267 and yellow cells are those expressing both VSP. Note that the presence of the INR does not promote the expression of VSP417 in VSP1267 expressing trophozoites. However, the 5’-upstream region of housekeeping genes (no INR) promote a high percentage of cells to express both VSP. The 3-downstream regions seemed irrelevant.

### 1.7. 3’-UTR and Polyadenylation sites – Alternative Polyadenylation (APA)

In earlier reports it was stated that the presumed poly(A) signal sites (PAS) of *Giardia* genes was AGRAAA, where R is a purine, resembling that of higher eukaryotes AAUAAA [51,55]. In a recent report, using long read sequencing, it was verified on a genome-wide scale, that *G. lamblia* uses an AGURAA PAS [64]. In that study, AGUAAA and AGUGAA were found in 45% and 15% of protein encoding genes, respectively. In contrast, AAUAAA was rare, occurring in only 5% of genes. The most frequent PAS, which is AGUAAA, differs from the metazoan AAUAAA motif by only a single nucleotide. Moreover, these hexamers were found to be depleted in coding regions while, occasionally overlapping with stop codons, like in the murine parasite *G. muris* [65]. The 3’-UTR lengths generated by that approach had a median of 59 nucleotides, which is consistent with the general idea that 3’ UTRs of *G. lamblia* are unusually short [51,64]. *Giardia* PAS were typically found less than 20 nt upstream of the cleavage site, and the most common distance was 13–15 nt.

Having this in mind, the presence of PAS in the 3ˈ end of all Cys-rich genes was determined. All 136 VSPs have one of the two PAS, and the hexamer is found less than 28 nt downstream of the stop codon. The results showed that all 136 VSPs have an extended PAS with the sequence ACUUAGRU**AGURAA**YRY (R= purines and Y=pyrimidines) (**S5A fig**), placed in a median of 6 nt from the stop codon (**S5B fig**). This extended PAS is not found in most of the CRMPs and the 4 cases where something similar is found the C-terminal pentapeptide is similar to CRGKA. Thus, the search an extended PAS in the 3’UTR of all *Giardia* genes was performed and it was found that the motif is intimately associated with VSPs and pVSPs. Manual curation of the detected motifs revealed that 323 of 327 hits are associated with VSPs and pVSPs with the remaining 4 cases being found in CRMPs (**S6 table**).

The cleavage site of some VSPs was found between 16 and 38 nt downstream of the coding region [51,64]. For example, VSP GL50803_00113797 was described as having a 27-nucleotide 3’ UTR, and our analysis detected the hexamer AGUAAA located 15 nt upstream the putative cleavage site and 12 nt downstream the stop codon. Likewise, VSP GL50803_00112208 was described as having a 24-nt 3’ UTR, and the PAS AGUAAA was found 16 nt upstream the cleavage site and 8 nt downstream the stop codon.

For previously annotated HCMPs, the length of the 3’ UTR has been determined to be much more variable, ranging from only a few nt up to thousands (between 7 and 2000 nt downstream of the coding region) [51,64]. Consistent with this finding, only 67 of 94 CRMPs have AGURAA in the 2000 nt downstream of the stop codon. The same was true for SCRPs, only 11 of 16 have AGURAA in the 2000 nt downstream of the stop codon. The remaining CRMPs and SCRPs might have the PAS located farther away or might have another signal. In this regard, 13 PAS were previously described for all *Giardia* genes [51]. Another explanation could be that a strict PAS may not be required for proper 3’-end formation, as it was also suggested [51].

Again, plasmids were designed to express VSP417 using different 100-nt3’-regions (VSP417, OCT or TUB) alternating with the 5’ promoter regions described above. In any case, an expression greater than 5% was never achieved, as determined by IFA using mAb 7C2 to detect the episomally encoded VSP417 (**Fig 7B**). These results indicate that, regardless the downstream region, no differences were observed in the level of expression of an exogenous VSP.

For that reason, the presence of alternative polyadenylation (APA) was analyzed and evidence that some VSP genes show APA was found. For example, VSP GL50803_0040591, which is, as it will be described later, one of the preferably expressed VSPs in culture, has two AGUAAA PAS in its 2000-nt downstream region. The PAS are located 12 and 1274 nt downstream of the stop codon. Interestingly, when a single cell RNA-seq experiment was reanalyzed (see later), in some cells this VSP was found to use the first PAS, giving place to a transcript with a very short 3’-UTR (**Fig 8A**), whereas in other cells the transcript of this VSP was found to have a significantly longer 3’-UTR and the second PAS was found nearby (**Fig 8B**). This is, to our knowledge, the first description that APA might occur in the VSP gene family. This was also seen with other VSPs, such as GL50803_00113797. APA was not as evident in other Cys-rich proteins, although its occurrence also in CRMP- and SCRP-encoding genes cannot be ruled out.

**Fig 8.**
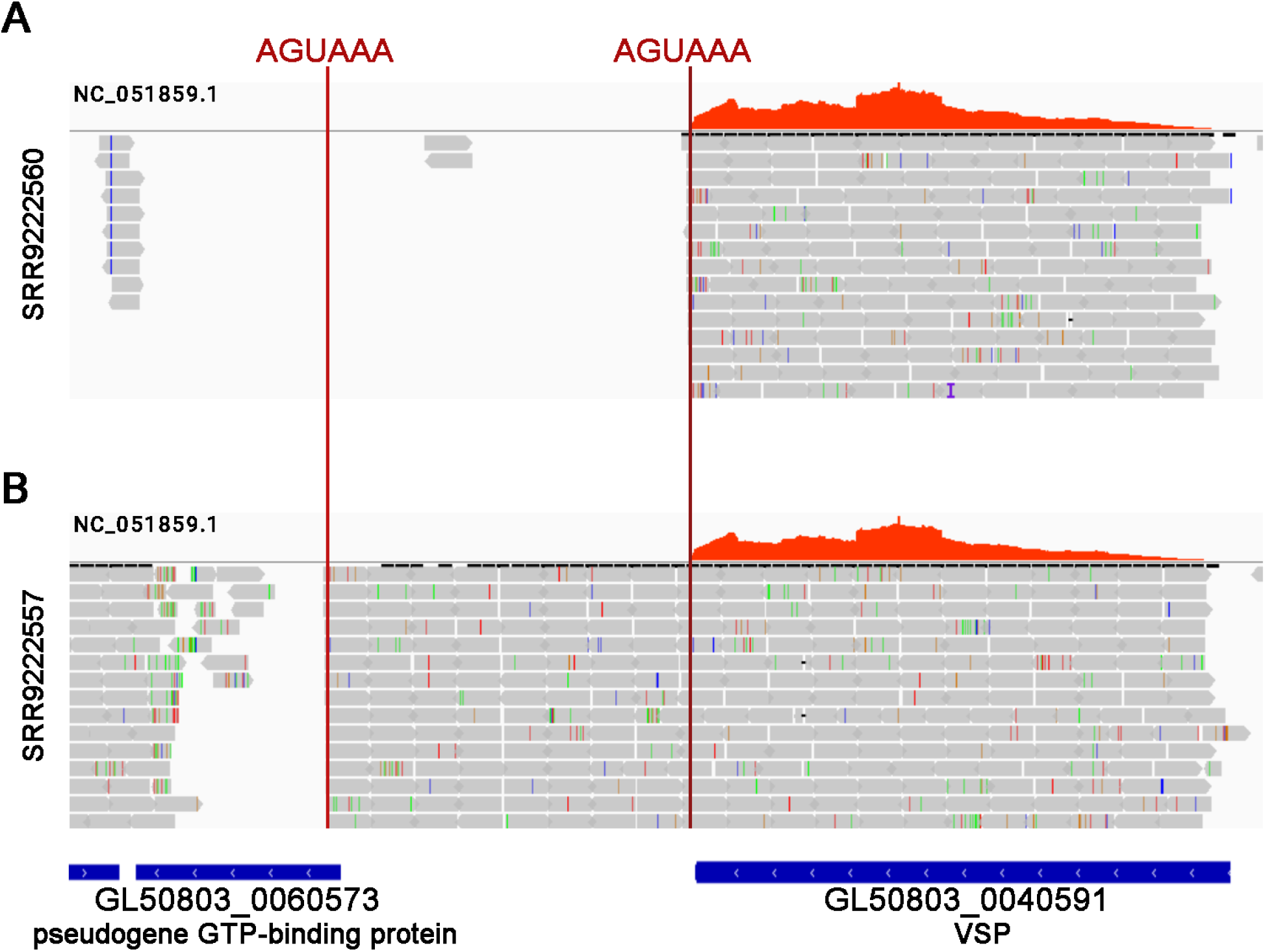
Evidence of alternative polyadenylation in a VSP transcript. IGV snapshots showing coverage and reads of a single cell experiment that aligned to VSP GL50803_0040591. In cell **A**, this VSP has a very short 3’ UTR, whereas in cell **B** it has a longer one, consistent with the two PAS (vertical red lines) found at the 2000-nt downstream of the stop codon.

## 2. Cys-rich Pseudogenes

All pseudogenes (type I and type II) were divided into pseudo VSP (pVSP), pseudo CRMP (pCRMP) or pseudo SCRP (pSCRP) based on their similarity to each group, using BLASTx against a database constructed with the VSP, CRMP and SCRP sequences defined in this work. The database sequence with the highest score and the lowest E value was taken into account to define whether a pseudogene should rather be named pVSP, pCRMP or pSCRP.

A vast number of pVSPs, representing ∼89% of total Cys-rich pseudogenes, were detected. Interestingly, even though the remaining ∼11% of the sequences BLASTed first with a Cys-rich protein other than a VSP, they also BLASTed with VSPs with lower query coverage or score. This finding suggests that the differences between VSPs and CRMPs or SCRPs are not as significant as speculated previously. In contrast to the abundance of pVSPs in the genome, pCRMP and pSCRP are scarcer, representing only ∼9% and ∼2% of total Cys-rich pseudogenes, respectively.

Some examples of pVSPs are shown in **Table 5**. For example, pVSP GL50803_0050472 has multiple internal stop codons, and a closer inspection revealed that the putative coding sequence is split across all three frames, which can be explained by insertions/deletions that occurred during evolution. Most of pVSPs without an ORF and without a proper start codon are incomplete and lack the 5’ portion due to their location in unassembled scaffolds, near a chromosome end or in arrays of multiple pVSPs, as shown for pVSP GL50803_0050404. On the other hand, pVSP GL50803_0050265 has a complete ORF starting with a Met and ending in CRGKA, but no SP could be predicted in any Met in frame. pVSPs truncated at the C-terminal end, without a proper TMD or CT, were also found, and pVSP GL50803_0050028 is an example of this group. Moreover, a thorough analysis of the 5’-regions of each of these pseudogenes revealed that in a small group of pVSP with ORF but without SP, it is possible to find a SP in a frame other than the ORF. The presence of these numerous partial and remnant VSP genes raises questions about their function. Additional information about Cys-rich pseudo genes can be found in **S7 table**. Intergenic regions with similarity to Cys-rich genes were also found, suggesting that there could be more pseudogenes of these families than those annotated (**S7 table)**.

**Table 5.**
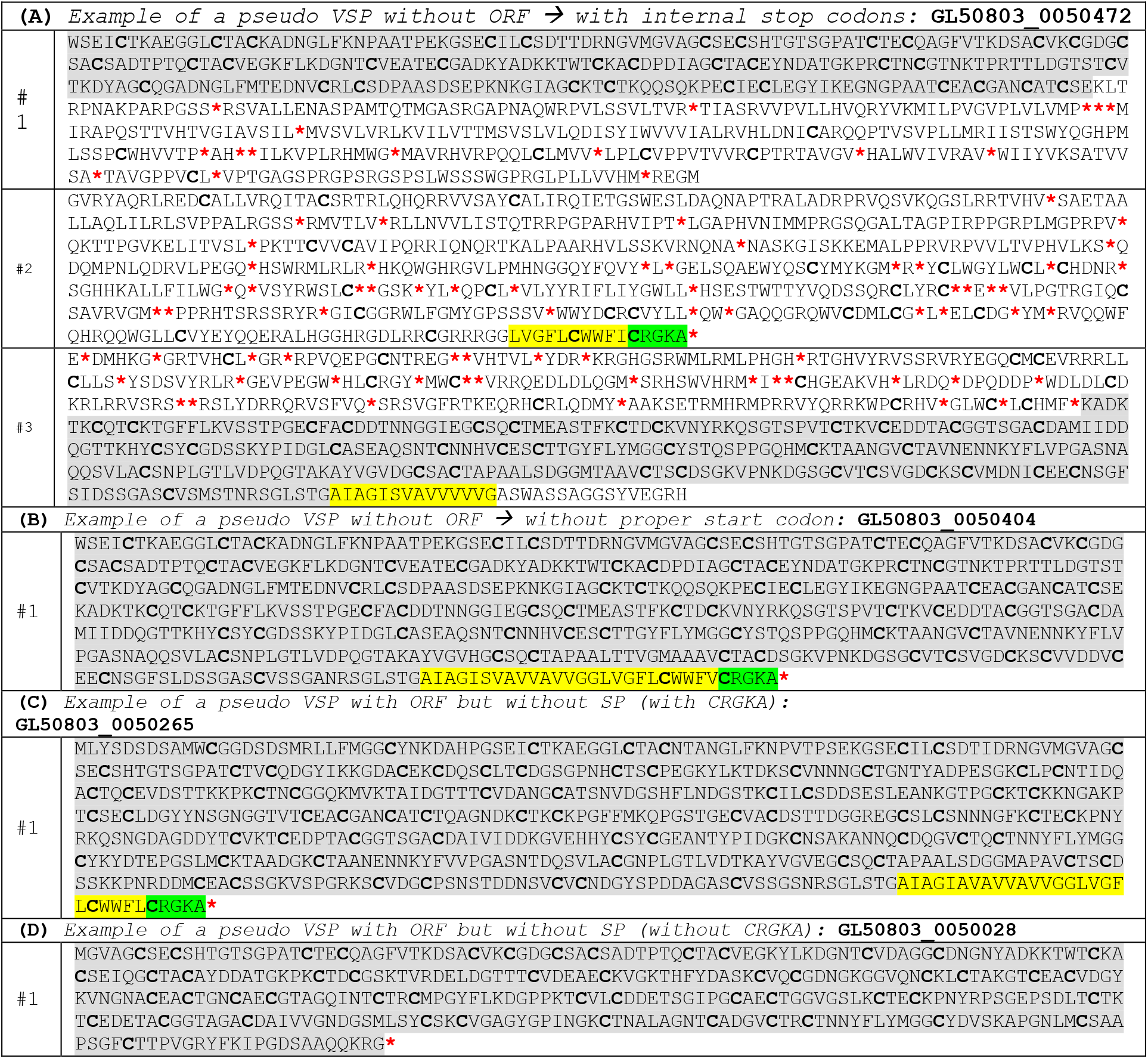
Examples of pseudo VSPs. The reconstruction of the ORF by searching in the three frames is highlighted in grey. If found, the TMD and CT are highlighted in yellow and green, respectively. #1, #2 and #3 represent frame 1, 2 and 3, respectively. **(A)** pVSP GL50803_0050472 is a pseudogene annotated without ORF and with internal stop codons. However, a closer inspection revealed that the putative coding sequence is split across all three frames, which can be explained by insertions/deletions that occurred during evolution. In the region highlighted in grey no stop codon is found, but it is split between #1 and #3. In #3, it is found half of a putative TMD, which continues in #2. A CT is also found in #2. **(B)** pVSP GL50803_0050404 is a pseudogene without ORF that has no internal stop codons, but no proper start codon can be found. This is seen in many pVSPs that are incomplete and lack the 5’ portion due to their location in unassembled scaffolds, near a chromosome end or in arrays of multiple pVSPs. **(C)** pVSP GL50803_0050265 is a pseudogene that has a complete ORF starting with a Met and ending in CRGKA, but no SP could be predicted in any Met in frame. **(D)** pVSP GL50803_0050028 is a pseudogene with truncated C-terminal end and without a proper TMD or CT.

## 3. Cys-rich proteins across isolates, assemblages and species

VSPs, CRMPs, SCRPs and pseudogenes are not only present in the *G. lamblia* isolate WB (assemblage A1), to which the GL2.1 genome belongs to, but also common to the other sequenced assemblages and the related species *G. muris*. All VSP sequences in *G. lamblia* isolates have a similar TMD and a CRGKA polypeptide in their C-terminal CT, but in *G. muris* the CT has a 4-amino acid CT (CRGK). They also have the same preference for amino acids at the CXXC motifs. Interestingly, there are not identical Cys-rich proteins among all isolates that have been sequenced, including other members of the assemblage A such as DH, which suggests that each group presents its own repertoire of these protein families (only CRMP GL50803_0011690 on WB and DHA2_11690 on DH were found in both isolates).

VSPs of *G. muris* and *G. lamblia* DH, GS, GS-B, P15 and WB isolates were compared across multiple parameters, such as length, number of CXXC motifs and relationship between length and number of CXXC motifs. Except for isolate WB, sequences were obtained from the *Giardia*DB, but only *G. muris* has a genome which is near completion, only rDNA regions are assembled [65]. Analyses were carried out taking into account only the sequences that presented an SP, a conserved TMD, CRGKA (or CRGK) and multiple CXXC motifs. In total, of 329 sequences, 136 correspond to WB, 93 to P15, 1 to GS, 44 to GS-B, 32 to DH and 23 to *G. muris*. The *G. muris* VSP organization is different than the *G. lamblia* WB organization with few complete VSPs (only 16 unique) and many pVSPs (239) in long arrays in the telomeric regions lacking SPs. Phylogenetic analyses indicated that these pVSP arrays might serve as a reservoir of gene fragments for gene conversion to generate variability in the VSPs [65].

CRMPs and SCRPs of *G. muris* and *G. lamblia* DH, GS, GS-B and P15 were reclassified according the decision tree proposed in this work. For this analysis, only well-annotated sequences with an initial Methionine were taken into account. For 392 sequences that present the parameters of a CRMP, 94 correspond to WB, 88 to P15, 37 to GS, 84 to GS-B, 58 to DH, and 31 to *G. muris*. In the case of SCRP, 169 sequences were found and 16 correspond to WB, 34 to P15, 23 to GS, 53 to GS-B, 34 to DH and 9 to *G. muris* (**S8 table**). It is important to note that the poor assembly of some chromosomes of the other *G. lamblia* isolates than WB, as well as the deficient annotation of the sequences, makes it difficult to classify these Cys-rich family proteins correctly.

*G. muris*’ VSPs have homogenous length, whereas *G. lamblia* WB VSPs are the longest compared to all other isolates. GS presents the highest length/CXXC motifs ratio whereas *G. muris* present the lowest ratio. *G. muris* presents the highest mean of CXXC motif, whereas GS presents the lowest mean (**S6 fig**).

## 4. Transcriptomic and proteomic analyses among all isolates

Most of the available RNA-seq experiments in databases were performed with non-clonal parasite lines or with mixed populations of trophozoites. Besides, most of them used the previous version of the *Giardia* genome as reference. Therefore, it was not possible to obtain a clear expression profile for VSPs and related genes. To minimize the heterogeneity of expressed VSPs in culture, a clonal trophozoite population from the original WB isolate “clone” C6 was generated using serial dilutions. Sequencing libraries were prepared according to a strand-specific protocol using rRNA-depleted total RNA from 2 different clones, one expressing the VSP417 (formerly named TSA417 [66]) (GL50803_00113797) and the other expressing a VSP duplicated pair (VSP1267; GL50803_00112208 and GL50803_00d112208 [52]). The expression of these VSPs on the cell surface was determined by immunofluorescence assays using a specific mAb, as reported [6] (**Fig 9**). These were called clone 1 and clone 2, respectively. From clone 1, triplicate replicas were generated (rep1, rep2 and rep3) and from clone 2, a single replica (rep1) was used. The sequencing protocol and the pipeline used for analysis are described in materials and methods, and the new version of the *Giardia* WB genome was used (RefSeq accession: GCF_000002435.2). On average, 80.3% of all sequenced reads could be mapped to a single location in the genome assembly.

**Fig. 9.**
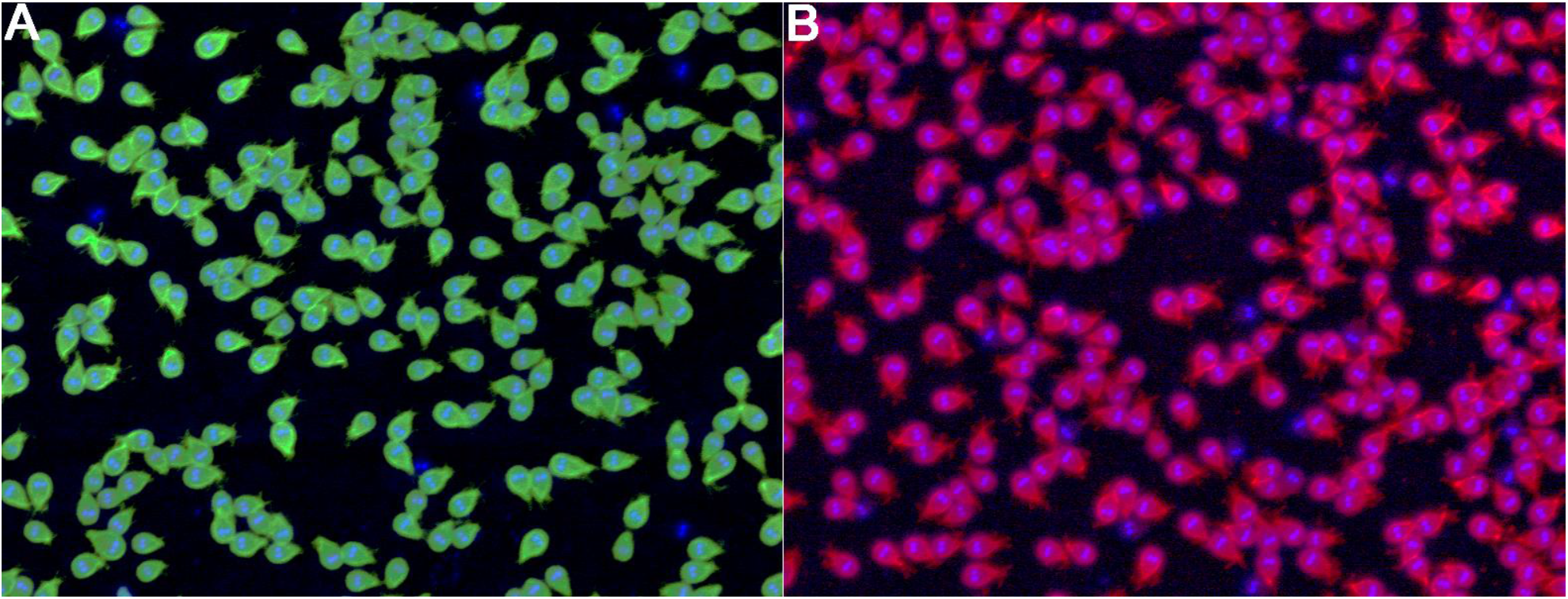
Immunofluorescence images of two clonal trophozoite population derived from the original WB isolate C6, stained with a monoclonal antibody (mAb) that specifically recognizes a single VSP. **(A)** Clone 1 stained with mAb 7C2, specific for VSP GL50803_00113797 (VSP417), and **(B)** Clone 2 stained with mAb 8F12, specific for VSP GL50803_00112208/GL50803_00d112208 (VSP1267). Cell nuclei were stained with DAPI. Magnifications 200X. Note that more than 90% of the trophozoites in each population express the corresponding VSP on their surface.

The highest transcribed (or accumulated) transcript in clone 1 was the one corresponding to VSP417 (GL50803_00113797) when using unique mapping reads (**Fig 10A, left panel**). On the other hand, in clone 2, VSP1267 (GL50803_00112208 and GL50803_00d112208) was not detected as major transcripts, because they are a duplicated pair and have very few unique counts (**Fig 10A, left panel**). Since many VSP genes are duplicated, two strategies for counting reads mapped to these duplicated genes were used. In one strategy, unique mapping reads (UM) were considered. In the second strategy, multi-mapping reads were used, setting to 2 the maximum number of loci that a read is allowed to map to (2M). In the latter scenario, multi-mapping reads were fractionally counted (each alignment carrying 1/x count, where x is the total number of alignments reported for the read, in this case x=2). Using this strategy, it was possible to incorporate approximately 3.35% extra reads mapping to two locations in the genome.

**Fig 10.**
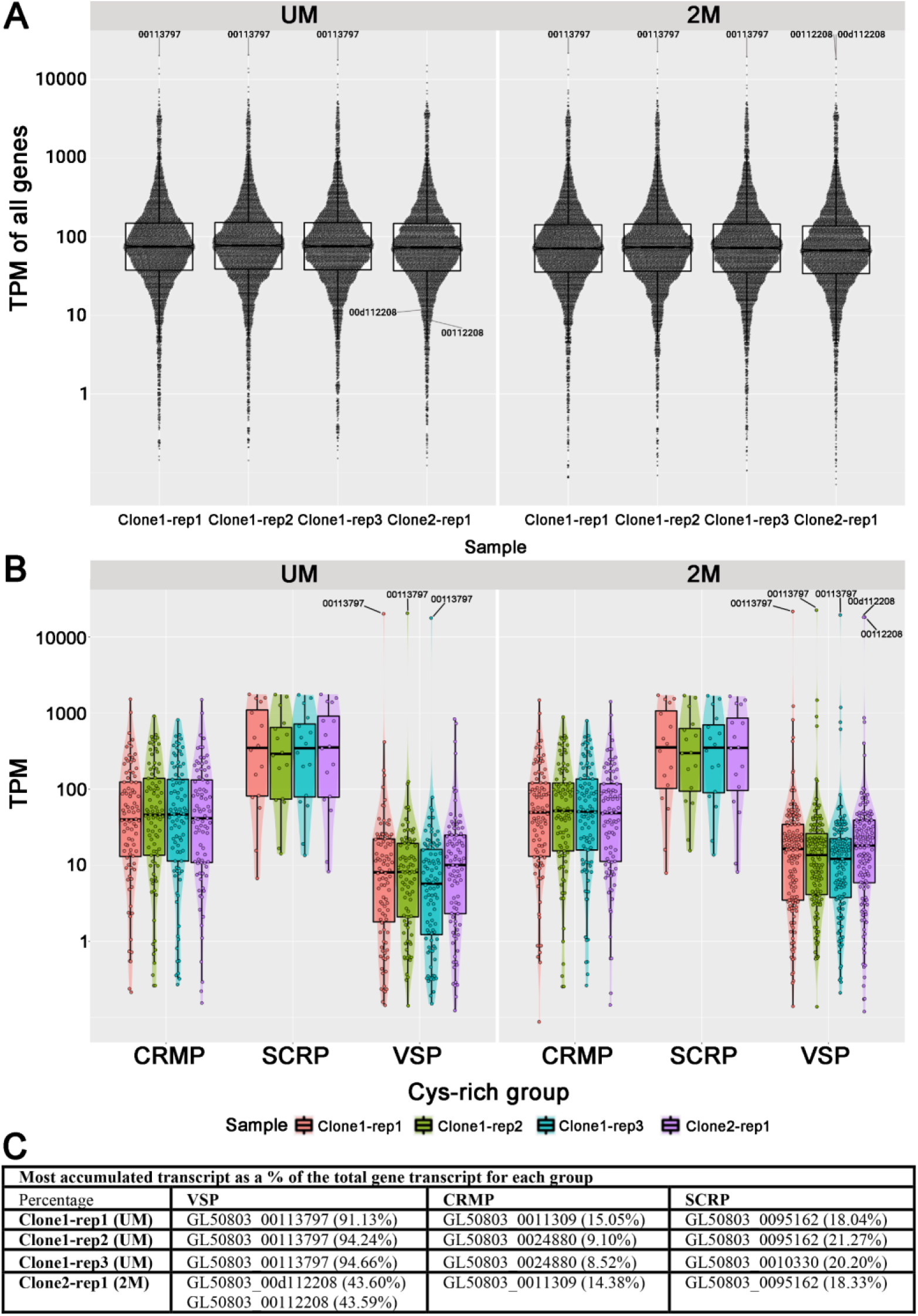
Analysis of our own RNA-seq experiment. **(A)** Gene expression/transcript accumulation in TPM of all genes and pseudogenes. **(B)** Gene expression/transcript accumulation in TPM of genes corresponding to each Cyr-rich family: CRMP, SCRP and VSP. **(C)** Most accumulated transcript as a percentage of the total gene transcript for each group.

When the 2M strategy was used, the pair GL50803_00112208/GL50803_00d112208 (VSP1267) corresponded to the most accumulated transcripts in clone 2 (**Fig 10A, right panel**). At this point, it cannot be discriminated whether duplicated VSPs are expressed together or only one member of the pair is transcribed; however, whichever the case, the VSP that is expressed in the plasma membrane correlates with the transcript that accumulates the most in each clone, even far above housekeeping genes. Except for the specific VSP of each clone, the transcripts that accumulate the most are the same in both clones (e.g. Ribosomal protein S15A, Ornithine Carbamoyl Transferase, Dynein light chain LC8, Ribosomal protein P1B, β-Giardin and Cytosolic Heat Shock Protein 70, just to mention a few). Transcript per million (TPM) values for each transcript in each sample can be found in **S9 table**.

The analysis was then focused on VSP, CRMP and SCRP transcripts. VSPs are transcribed and accumulated at different levels. However, it is clear how the VSP that is translated into protein is the one that accumulates the most and its abundance differs radically from the remaining transcripts of this gene family (**Fig 10B**). For each sample, the transcript level of an individual VSP gene was calculated as the percentage of the total VSP transcript level in that sample. It was found that in clone 1, more than 90% of the total VSP gene transcript level is monopolized by VSP417. In the case of clone 2, using the 2M strategy for counting, the VSP1267 monopolized 87.2% (∼43.6% each) of the total VSP gene transcripts (**Fig 10C**). This indicates that clones expressing a single surface antigen efficiently transcribe several VSP genes, but only accumulate in abundance transcripts encoding the VSP that is translated and expressed on the trophozoite surface.

A similar analysis was done with individual CRMP and SCRP transcripts. CRMPs and SCRPs are also transcribed and accumulated at different levels. However, no single transcript differs radically from the rest in term of its TPM (**Fig 10B**). Unlike VSPs, where there is always one gene that accounts for nearly 90% of all VSP transcripts in clonal populations, in CRMPs and SCRPs none exceeds ∼15% and ∼21% of the total, respectively (**Fig 10C**).

Moreover, biological triplicates of clone1 were incubated for 4 h with low concentrations of mAb 7C2 to induce VSP switching and RNA sequencing was performed (Clone1T-rep1, Clone1T-rep2 and Clone1T-rep3). The aim was to identify differentially expressed genes (DEGs) that might be related to the process of antigenic variation triggered by antibodies (<u>Tenaglia, Luján, Ríos *et al*., accompanying manuscript for Reviewers’ consideration only</u>). The sequencing protocol and the pipeline used for analysis was the same as that described for clone1 without treatment. EdgeR was used to obtain the DEGs.

Surprisingly, only 7 DEGs (FDR<0.05) were found between trophozoites before and after treatment (**S10 table**). The DGEs encode 6 formerly known High Cysteine Membrane Proteins now called CRMP (GL50803_00137727, GL50803_00112135, GL50803_0025816, GL50803_00113531, GL50803_00115066, and GL50803_00114930) and 1 hypothetical protein (GL50803_0087826), which were all upregulated in anti-VSP antibody-treated clones. The level of upregulation was extremely small, with the highest logFC found being 1 (**S10 table**). These small logFCs were previously reported, although in experiments not related to antigenic variation, suggesting either a very tight level of regulation or post-transcriptional regulation [12].

Even more surprisingly was the fact that although coming from a single clone, the triplicates have a vast intrinsic variation. However, as it will be shown below, the single cell transcriptomic analysis also shows a massive variation in expression/accumulation mRNA levels among all *Giardia* genes, clearly evidencing the stochastic nature of gene expression in this organism. Stochastic gene expression occurs when transcriptional regulators are present at very low concentrations, promoting that binding and release of regulators from their binding sites becomes probabilistic. The assumption that stochastic gene expression has a significant effect on the biology of organisms comes from the observation that genetically identical (clonal) organisms, maintained in identical environments, diverge phenotypically. For instance, cell division in bacteria growing in an optimum medium rapidly becomes asynchronous, presumably due to individual stochastic variation in regulatory processes [67].

As mentioned above, most *Giardia* isolates are mixed populations of trophozoites expressing different VSPs and the data obtained from those experiments make it difficult to discern whether all VSPs are transcribed in each trophozoites or whether VSP transcription within the population is different. Besides the present study, there is only one report in which the authors performed RNA-seq of a clonal population [12]. These authors were interested in assessing the effects of iron on the total *G. lamblia* transcriptome; therefore, they performed RNA-seq on trophozoites grown in media with different levels of free iron. The raw data from 4 SRA randomly chosen samples from those experiments were downloaded and analyzed using the described pipeline. These samples were called Clon3-rep1 to Clon3-rep4. In **S11 table**, the corresponding SRA accession numbers are indicated. Unlike our RNA-seq, these libraries were prepared including a polyA selection step. The aim now was to evaluate the transcriptional profile of VSPs, CRMP and SCRP in this clone and to compare those results to ours. Similarly, it was found that this clone highly accumulates the transcript corresponding to a single VSP, in this case GL50803_00113450, which monopolizes between 74.48% and 78.79% of the total VSP gene transcripts (**Fig 11A**). Again, this characteristic of a single transcript accumulated above others is not observed for any member of the CRMP or SCRP families (**Fig 11A**).

**Fig 11.**
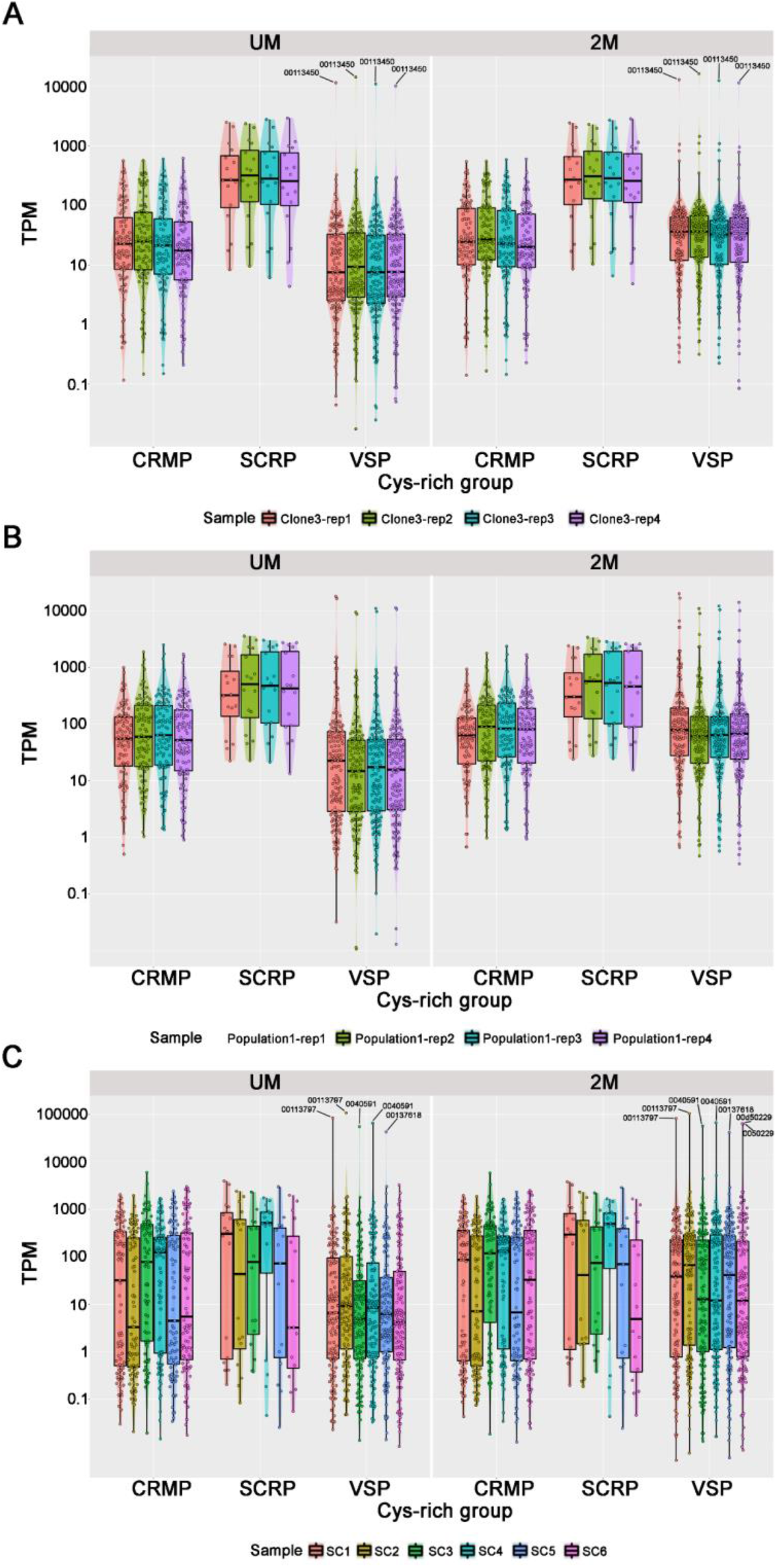
Analysis of RNA-seq experiments from other authors. Gene expression/transcript accumulation of the Cys-rich genes in **(A)** the clone (clone3) used by Peirasmaki *et al*; **(B)** the non-clonal population (population1) used by Peirasmaki *et al*; and **(C)** the single cell (SC) experiment performed by Onsbring *et al*. Note that the characteristic of a single transcript accumulated radically above others is only observed in the VSP family (labeled with the corresponding gene ID).

In another set of RNA-Seq experiments, Peirasmaki *et al*. [12] used an *in vitro* model of interaction of trophozoites with intestinal epithelial cells to identify genes of *G. lamblia* that might correlate with the establishment of infection and disease induction. Unlike the experiment on the effect of iron, in their work they used a non-clonal population of trophozoites. The raw data of 4 randomly chosen samples from SRA were collected and analyzed using our own pipeline. These samples were called Population1-rep1 to Population1-rep4. The corresponding SRA accession numbers are indicated in **S11 table**. In this population, no single VSP dominated most of the transcripts (**Fig 11B**). In fact, two VSP transcripts corresponding to GL50803_0041472 and GL50803_0040591 have only ∼35% of the total VSP transcripts each, indicating that this population is composed of two different clones. This comparison shows the importance of using clonal population of trophozoites to analyze VSP gene expression and raises questions about previous conclusions taken by population-based RNA-seq.

Furthermore, the raw data of a single cell RNA-seq experiment available in the literature was used [68]. In that study, the authors tested different modifications of Smart-seq2, a single-cell RNA sequencing protocol originally developed for mammalian cells, to establish a robust and cost-efficient workflow for protists. They generated 55 transcriptomes of single *G. lamblia* cells. Unfortunately, the authors used the GL2.0 genome for analysis. Libraries were prepared according to an unstranded protocol. The raw data from SRA SRR9222552 to SRR9222606 was analyzed using our own pipeline and the new reference genome. Among 55 cells analyzed, 30 accumulated VSP GL50803_00113797 as the major transcript (the same VSP found in our clone 1 and called VSP417), 11 cells accumulated VSP GL50803_0040591 and the rest accumulated other minority VSPs or pairs of duplicates. Six representative samples were selected for a closer inspection and these samples were called SC1 to SC6 (**S11 table**).

SC1 and SC2 accumulated VSP GL50803_00113797 and this VSP monopolized 87.49% and 86.18% of the total VSP gene transcripts, whereas SC3 and SC4 accumulated VSP GL50803_0040591 (also known as VSPA8) and this transcript monopolized 84.72% and 82.23% of the total VSP gene transcripts, respectively. SC5 accumulated VSP GL50803_00137618 with 74.70% of total VSP transcripts in that cell (**Fig 11C**). These samples were chosen to show that despite coming from a single isolate (ATCC 50803, WB “clone” C6), different cells accumulate a single VSP transcript that is not always the same. On the other hand, SC6 does not accumulate any VSP with more than 17% of the total VSP transcripts, using the UM strategy. Therefore, for this cell transcriptome, the 2M strategy was used. This was done to discern if a pair of duplicated VSPs was expressed (as occurred with clone 2 of our own RNA-seq). It was found that, indeed, SC6 accumulated the VSP pair GL50803_0050229/GL50803_00d50229, and contributed ∼40% each to the total VSP transcripts in that cell (**Fig 11C**). As before, it is impossible to determine whether both genes are being expressed at the same time or only one of them is expressed.

Interestingly, no condition in which samples accumulate transcripts from all 136 VSPs at the same time was found in our RNA-seq, in the one performed by Peirasmaki *et al*. [12], or in the 55 single cells transcriptome analyses [68]; there are always some VSPs with no TPM. It seems that in the same cell or clone, not all VSP genes are either transcribed or accumulated at detectable levels.

However, the analysis of more than 90 transcriptomes deposited in databases showed that practically all VSP genes (130/136) have evidence of transcription. To reach this conclusion, 91 transcriptomes were re-analyzed including the ones mentioned above and others that used populations of trophozoites. All the transcripts whose expression (measured in TPM) was included in the upper quartile of each of the 91 samples were selected. Then, of all the transcripts obtained, a subset of those corresponding to VSP genes was formed, and 121 VSP transcripts using UM strategy and 9 extra VSP transcripts using 2M strategy were found. This gave 130 out of 136 VSP genes with evidence of transcription in the upper quartile in all the samples analyzed.

Remarkably, it looks like few VSPs are preferably selected for expression over others in culture conditions because they appear more frequently as the major transcript that accounts for most of all VSP mRNAs (e.g. GL50803_00113797, GL50803_00113450, GL50803_0040591, and GL50803_00112208/GL50803_00d112208). The expression of these particular VSP did not correlate with either a specific chromosomal location or the proximity to a highly expressed housekeeping gene. Even more interesting is the fact that preferable VSPs are those of predicted molecular weight close to the median (74 kDa), neither too short nor too long. For example, VSP GL50803_00113797 has 714 aa in length and its predicted molecular weight is 82 kDa. However, almost all VSPs can be transcribed at some point under a certain condition.

In this regard, there is a transcriptomic experiment designed to look for a differential gene expression between a *Giardia* foci in the small intestine and *in vitr*o trophozoites grown in axenic culture [69]. The authors found several differentially expressed VSPs, some of them more highly expressed *in vivo*, and some more highly expressed *in vitro*. One could hypothesize that there are preferable VSPs that are expressed in trophozoites *in vitro* and a different subset of VSPs are the preferable when trophozoites colonize the small intestine. There is also evidence that the VSPs with highest expression in trophozoites are not the same as the VSPs expressed in cysts, and that there are expressional changes of several VSPs during encystation [70]. Regarding CRMP, most of them are up regulated *in vivo*, suggesting that CRMPs are important during host-parasite interactions. This up-regulation of formerly HCMPs was also seen *in vitro* when trophozoites where incubated with Caco-2 cells [12]. The expression of the CRMPs and tenascin-like proteins also changed during encystation [69]. As is the case with most CRMPs, the HCNV is overexpressed in trophozoites *in vivo* with respect to trophozoites grown *in vitro* [71]. However, no significant changes were found in the expression of HCNV in trophozoite-epithelial cell interaction experiment [12].

A similar analysis was carried out with CRMP and SCRP genes, and evidence for transcription of all 94 CRMPs (91 using UM strategy plus 3 using 2M strategy) and 16 SCRP genes (16 using UM strategy) was found.

Since both Peirasmaki *et al*. [12] and our own RNA-seq were performed using a strand-specific protocol, the relationship between sense (S) and antisense (AS) transcripts in each Cys-rich group was then analyzed. A subset of 79 housekeeping genes was selected (**S12 table**) and it was found that they all have few antisense counts. In other words, in these genes the amount of antisense transcripts is minimal, and the S/AS ratio is always positive (100% have more S than AS transcripts) (**Fig 12A**). The analysis of VSP genes showed that only between 12-20% has more S than AS transcripts, depending on the clone being analyzed (**Fig 12B**). This leaves the remaining VSPs with no transcripts at all or more AS than S transcripts. In the case of CRMP and SCRP, these percentages rise to 36-41% and 75-87%, respectively (**Fig. 12C and D**). Exactly the same results were found in the analysis of the RNA-seq of Peirasmaki *et al*. [12] and when the 2M strategy was used. AS transcripts appear to be very abundant in VSPs and to a lesser extent in CRMPs, and even less in SCRP. It remains unclear whether the AS transcripts have regulatory functions in controlling gene expression and/or they are the consequence of a loosely regulated transcriptional process, as suggested [72].

**Fig 12.**
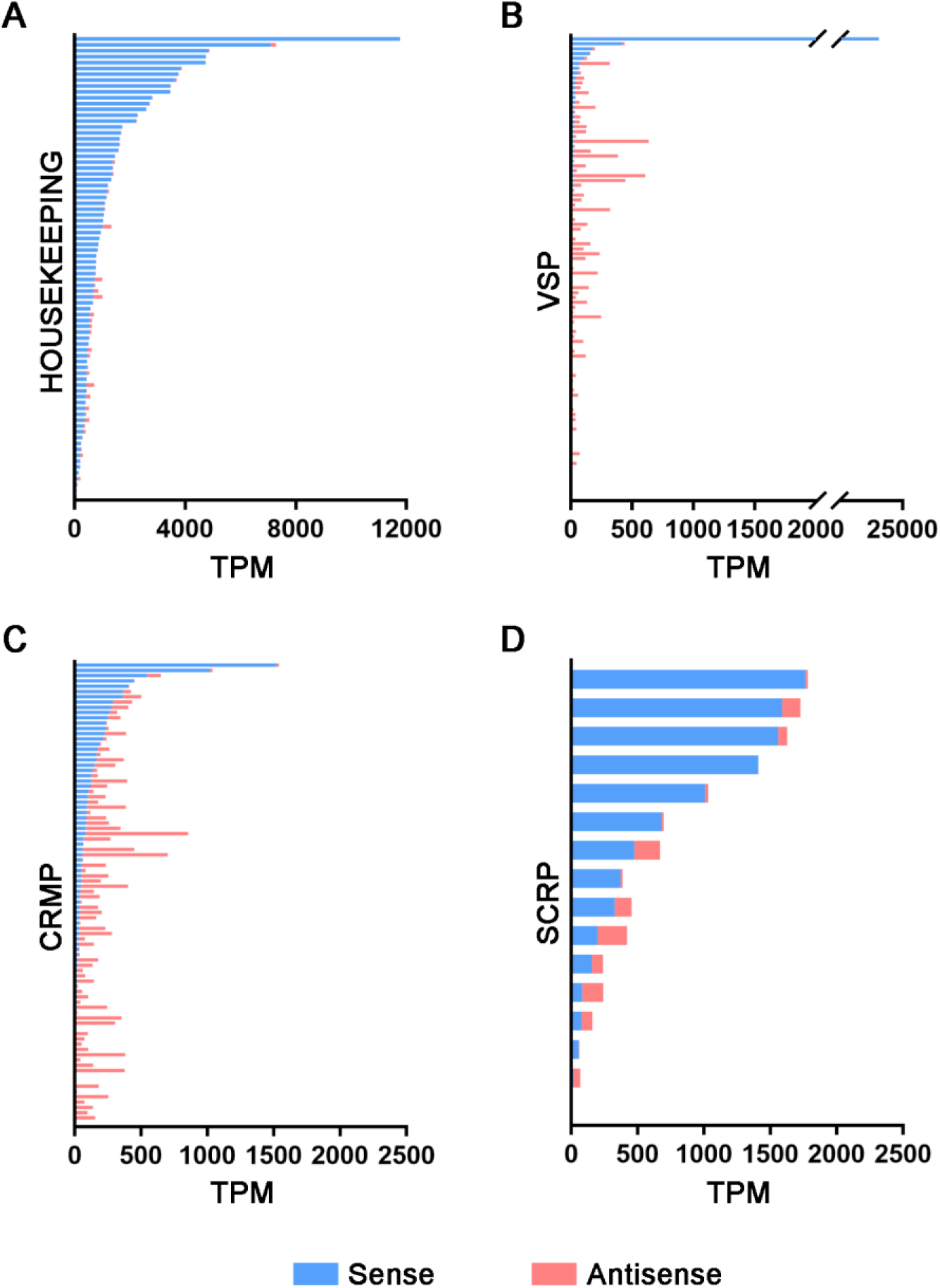
Stacked bar graph comparing the amount of accumulated sense and antisense transcripts in Clone1-rep1 for the different gene pools selected. Each bar represents one gene. **(A)** Selected housekeeping, **(B)** VSP, **(C)** CRMP and **(D)** SCRP genes. Similar results were found in the remaining clones analyzed in this work.

Regarding proteomic data, earlier studies performed on the previous version of the *Giardia* genome and with parasite populations showed translational evidence for all groups of Cys-rich encoding genes [14,73–76]. Notably, in none of the previously performed proteomic analysis of *Giardia* were there fewer than 1,500 identified proteins (out of 4,966 predicted proteins), suggesting the lack of sensitivity of these approaches. Nevertheless, the raw data from mass spectrometry experiments were downloaded from the PRIDE repository and re-analyzed with the MaxQuant software using the specific parameters described in each of these reports, but using the proteins fasta file of the GL2.1 genome. The following PRIDE projects were used: PXD017597, PXD004398, PXD000452, PXD002398, PXD007183 and PXD022565. Additionally, two recent proteomic studies that used the update version of the *Giardia* genome were reanalyzed [77,78].

The results showed evidence of translation for only a subset of VSP proteins (29/136), with unique peptides, which is in agreement with the results of the transcriptional analysis. The analysis of the non-unique peptides showed that the number of VSPs with possible evidence of translation rises to 99 out of 136. For the other Cys-rich proteins, 29 CRMP have unique peptides and 31 additional CRMP have non-unique peptides, giving 60 CRMP with possible evidence of translation. Moreover, 13 SCRP have unique peptides, and 1 extra SCRP has non-unique peptides, totaling 14 SCRP with possible evidence of translation. The proteins with evidence of translation can be found in **S13 table**.

Interestingly, some peptides detected in those analyses were shared between VSPs and CRMPs, while others were shared between CRMPs and SCRPs, strongly suggesting that SCRP may derive from CRMPs and these from VSPs.

## Discussion

The diversity of Cys-rich proteins of *Giardia* was analyzed regarding their sequences, domains and motifs, as well as their potential function in the biology of this parasite. *Giardia* colonizes the lumen of the small intestine, where digestion of nutrients takes place. How *Giardia* can survive in this harsh environment has been suggested to depend of the resistance of the VSPs to proteolytic digestion, properties that are provided by the CXXC motifs and their capability to bind metals [9,42]. Additionally, VSPs are involved in the process of antigenic variation, making the parasite capable to avoid the constant immune pressure generated by their hosts by switching the expression of antigenically different VSPs [79]. Other Cys-rich proteins are also encoded in the *Giardia*’s genome, those formerly called HCMPs, HCPs, EGF-like proteins and Tenascin-like proteins, among others.

Here, after performing an exhaustive analysis of these groups of proteins, we defined a new classification that better describes their characteristics: the VSPs, the CRMPs, and the SCRPs. This much simpler classification is supported not only by the analysis of their sequences, but also by their patterns of expression both at the transcriptional and post-transcriptional levels.

Until the *Giardia* genome is perfectly assembled, the total repertoire of Cys-rich protein genes and pseudogenes can only be estimated. However, since the unassembled contigs only contain VSP pseudogenes, our estimation of the number of members of each of these families appears highly accurate. We predict the number of VSP genes to be 136 with only two uncertainties since GL50803_00113512 and GL50803_00101589 are difficult to classify and could either be considered pVSPs due to the fact that they do not have quite all the features of VSPs or they could also be viewed as newly evolved HCMPs with especially VSP like-regulation. In sum, our analyses clarify the differences between published works about the number of VSPs, CRMPs and SCRPs encoded by the parasite. These results provide general information that paves the way to better study the molecular mechanisms of *Giardia* adaptation, differentiation and virulence.

Despite their sequence variability, VSPs have strikingly constant features that are not shared by any other Cys-rich family. The presence of a CRGKA cytoplasmic tail at the C-terminal end of the protein, an extended PAS and the Inr element /Kozak-like sequence at the 5’-end of the gene seems to be the defining feature of the VSPs, in addition to the presence of an SP, multiple CXXC motifs in the ectodomain and a highly conserved TMD. The Inr/Kozak-like sequence and the extended PAS seem to be exclusive to VSP genes. Interestingly, we found that the extended PAS is associated with most pVSP genes. The high variability in the features of CRMPs (no Inr and no CRGKA CT) and their similarity to the VSPs in their ectodomains raise the question of whether these proteins accomplish functions analogous to that of VSPs.

Undoubtedly, the critical tools to determine the subcellular localization, expression, structure and behavior of these proteins are specific mAbs that allow the cloning of phenotypic variants. Previous results using tagged VSPs or HCMPs only provided unconvincing results. For example, Li *et al*. [17] reported only 73 genes as the putative repertoire of VSPs. They used experimental approaches to determine what is required for a VSP to localize to the cell surface by employing immunofluorescence assays of permeabilized and non-permeabilized trophozoites expressing various mutants of putative VSPs. They reported that when the expression of a VSP tagged at the C-terminus with a 3xMyc tag was examined, the anti-Myc antibody was able to readily detect the protein on the membrane surface of non-permeable cells. These results are questionable, since it is broadly known that (a) the C-termini of VSPs are cytosolic and (b) permeabilization is necessary to disrupt the cell membranes sufficiently to allow the passage of antibodies to the cell interior. In a well-designed experiment, detection of a VSP tagged at the C-terminus with anti-tag antibody could only be achieved in permeabilized cells. Additionally, the authors localized on the plasma membrane “VSPs” that lack the SP. These particular genes are now determined to be pVSPs both by Xu *et al*. [18] and in this work. Since all the subsequent conclusions are drawn based on these misconceptions, no clear conclusions can be drawn from those experiments.

On the other hand, formerly known HCMPs were classified into nine groups with apparently similar internal motifs, according to MEME and MAST analysis [11]. These authors reported that 7 of the 9 similarity groups contained regular repeat patterns. In addition to the repeated motifs, three groups (called VSP-like, EGF-like, and TMK-like) contained recognizable motif signatures. The proteins placed in the VSP-like group had similarity against the VSP domain of the Pfam database (PF03302), the EGF-like group had high similarities to the EGF functional domain (PF00008), and the TMK-like group scored highly to a HMM constructed using alignments from the extracellular domains from *E. histolytica* transmembrane kinases. Despite the attempt to classify these proteins, the authors also mentioned that some members of one group had instances of motifs from other groups, indicating that HCMPs are extremely difficult to classify because they differ in various parameters, and they can be grouped differently according to the parameter that is chosen for classification. In other words, the separation between current HCMP groups is diffuse. We have found that most CRMPs (74 of 94) have a marked majority of CXXC motifs over CXC or CXXXC motifs, as seen for the VSP group. However, there are 4 CRMPs, 1 annotated as Tenascin-like protein (GL50803_0094510) and 3 as hypothetical proteins (GL50803_00113565, GL50803_008733 and GL50803_0015450) that have a strikingly high content of CXXXC motifs over the other two. Similarly, there are 16 CRMPs with majority of CXC motifs, with only one being annotated as CXC-rich protein (GL50803_0014225). This can also be a way to classify CRMPs as CXC-rich, CXXC-rich, or CXXXC-rich, and these subclasses might represent specific groups of proteins with different functions.

Based on our observations, we thus propose that VSPs appeared earlier during evolution and confirm that the VSP repertoire arose from duplications and subsequent divergence. Only duplicated pairs are present within the VSP and CRMP families, always surrounded by retrotransposon relics. Conversely, SCRP have no duplicates but have sequences and peptides that can also be found in the VSPs. Therefore, mutations and deletions/insertions appear to be a common characteristic of these genes that have derived in VSP-like proteins (CRMP) that lost their Inr/Kozak-like region and changed their TMD and CT. Subsequent changes made the CRMP to lose their TMD becoming SCRPs.

We have previously identified 6 *G. muris* VSPs [80], and reported that they possess a typical *G. lamblia* VSP structure and motifs. These included 10 to 13% of Cys (mostly as multiple CXXC motifs), GGCY motifs, a variable amino-terminal region, and a highly conserved C terminus with the conserved amino acids CRGKA. In light of new genomic evidence obtained with whole genome sequencing, the C-terminal CT of *G. muris* VSPs was found to be CRGK [79], differing slightly from that of *G. lamblia*. Therefore, we have to assume that we misidentified the *Giardia* species that was isolated from a laboratory rat and that the VSPs of *G. muris* have a shorter CT that lacks Ala as the last amino acid.

Although CRMPs seem to behave similarly to the housekeeping genes of *Giardia* in transcriptomic and proteomic analyses of proliferating trophozoites, CRMPs are up-regulated during host-parasite interactions *in vivo* [71], during host-epithelial cell interactions *in vitro* [12], during encystation [70] and after confronting trophozoites to antibodies against their cognate VSP (this work). It seems that some relative CRMP transcript levels increase in response to stress factors. In the case of AV, it can be speculated that, due to their similarity to VSPs, CRMPs may serve as a protective cover of the parasite from the hostile environment of the small intestine when one VSP is being replaced with a different one during antigenic switching.

Finally, it is important to note that since the quality of a transcriptomic analysis is only as good as the reference data, it cannot be ruled out that, as the annotation of the *G. lamblia* genome improves; better conclusions can be drawn regarding Cys-rich families transcription profiles. Nevertheless, several key features of these groups of Cys-rich genes were identified.

First, when *Giardia* populations are analyzed, most VSPs appear to be transcribed at very low levels and antisense transcripts of VSPs can be found. In clonal populations, however, only one VSP transcript accumulates a very high levels, even 10 times higher than the most accumulated housekeeping genes OCT and ADI, among others. These results clearly demonstrate the mutually exclusive expression of the VSPs. Based on the transcriptomic analyses, it is not clear whether it this transcript accumulation is due to increased transcription or decreased decay of VSP mRNAs. Additional studies are required to provide novel insights into the molecular basis of this process.

Second, most CRMPs and SCRPs behave like any other housekeeping gene, having regular but variable levels of accumulation and proteomic evidence of expression. These results make a clear difference from the VSPs. Based on those observations, we support the fact that the SCRPs can be considered secretory proteins that serve as virulence factors [14], but we also propose that they can stabilize the structure of the VSPs by interacting either through their cysteines or by metal coordination.

Lastly, we provide a simple decision strategy to classify Cys-rich proteins in the genomes of other *Giardia* assemblages and species as well as a detailed pipeline for rapid and simple re-evaluation of improved genomes when they become available. This will be a simple tool to compare transcriptomes obtained under different experimental strategies and, to increase our understanding of the biology of this and other important human pathogens.

## Materials and Methods

### Sources of genomic, transcriptomic and proteomic sequences

Genomic, transcriptomic and proteomic data were obtained from RefSeq (GCF_000002435.2), the sequence read archive SRA and PRIDE proteomics repository, respectively. Genomic and transcriptomic data were visualized using the IGV_2.8.13 software [81].

### Computational characterization of cysteine rich proteins

The nucleotide sequences of the 4,966 CDS and of the 320 pseudogenes deposited in *G. lamblia* RefSeq database were collected. For annotated CDSs, the sequences that present more than 4% of cysteines in their protein sequence and at least two or more CXC, CXXC, and/or CXXXC motifs were included. Furthermore, a protein BLAST search was performed on the sequences retrieved in the previous step in order to keep Cys-rich proteins with similarity (E value <0.0001) to previously annotated VSP, HCMP, HCP and TLP. For annotated pseudogenes, a BLASTx search was performed using the nucleotide sequence as query. The presence of a predicted SP was determined using Phobius [16], SignalP-5.0 [20] and TOPCONS [21]. Phobius [16], TMHMM [19] and TOPCONS [18] were used to infer transmembrane regions. The subcellular localization of all Cys-rich proteins was predicted by DeepLoc 1.0 software [24].

### Sequence analysis

For sequence analysis, R and R studio were used with multiple packages like Biostrings, seqinr and stringr. Different parameters were searched using R for both nucleotide and amino acid sequences. Some of these parameters were the length of the protein, number of CXXC, CXC and CXXXC motifs, presence of PAS and of Inr. Statistical analyses were also performed in R. The amino acid sequences of the proteins rich in cysteines were analyzed using DiANNA Analysis 1.1 web server [40] to look for cysteines that coordinate metals such as Fe^2+^, Zn^2+^ and Cd^2+^. Conserved motifs in VSP were searched with MEME Suite 5.3.3 [36]. The classic mode was selected in the motif discovery mode. Zero or one occurrence per sequence was selected and the software was set to search for 10 motifs of up to 70 characters. No other parameter was modified. The XSTREME software of the MEME suite [36] was used to scan the 136 VSP 3’UTR regions (stop codon + 50 bp) of VSP genes for the presence of conserved motifs using standard settings. The top-matching MEME output was used to extract a sequence logo of the motif and the position. The Fimo software of the MEME suite was used to scan the 3’UTR region (stop codon + 50 bp) of all genes (including pseudogenes) in the *Giardia* genome using the extended PAS. 381 putative hits were reported at a p-value of 1E-5. The hits were manually curated, with 327 cases remaining after the hits showing poor motif conservation in otherwise high conserved positions, shifted placement, or wrong orientation were excluded. Using the manual analyses the cutoff representing a genuine hit had a p-value of 1.41 e^-07^.

### VSP and CRMP duplicated pairs and intergenic regions

To study the genomic context of the duplicated VSP and CRMP gene pairs, the specific intergenic region of each gene pair was analyzed and BLASTn was done using the BioProject 1439: *Giardia intestinalis* strain: WB C6.

### *In silico* VSP TMD oligomerization models

Oligomerization capability of the TMD was analyzed with TMHOP [82] and PREDIMMER [29] and visualized with VMD [83]. Alignments were performed with Clustal Omega [84] and visualized with WebLogo [85].

### Parasite culture

*Giardia lamblia* assemblage A1 isolate WB (ATCC^®^ 50803) was cultured in borosilicate glass tubes containing TYI-S-33 medium supplemented with 0.5 mg/ml of bovine bile (Sigma-Aldrich, Cat. #B3883) and 10% adult bovine serum (Natocor), as described [6]. Clones expressing different surface antigens were obtained by limiting dilution in 96-well culture plates placed in anaerobiosis chambers (Anaerogen® Compact, Thermo Scientific® Oxoid®, Cat. # AN0010C) at 37 °C during 5 days; positive clones were then selected using specific anti-VSP mAb by immunofluorescence assays (IFA). Reactive clones were then expanded in culture medium overnight and tested for homogeneity before use. The percentage of cells expressing particular VSPs was calculated by counting 500 cells in triplicate experiments or by flow cytometry using an Accuri® C6 flow cytometer (BD).

### Cloning and transfection

The plasmid pCIDIEpac was used to express variants of the 5’- upstream and 3’-downstream regions of VSP417. The 100-nt -upstream of the OCT gene or of the α-TUB gene and the 100-nt 3’-downstream of the G3P gene were used to drive expression of VSP417 (in some cases, the 100-nt upstream and downstream regions of VSP417 were also used), and the 100-nt 5’-upstream of the FBA gene and the 100-nt 3’-domstream of the β-TUB gene were used to express the puromycin resistance gene. Sequences were confirmed by sequencing using dye terminator cycle sequencing (Beckman Coulter). Trophozoites were transfected by electroporation and selected with puromycin (InvivoGen, Cat. # Ant-pr-5), as previously described [6].

### Transcriptomic analysis

To minimize the heterogeneity of expressed VSPs in culture, clonal trophozoite populations were created from the original WB C6 “clone” performing serial dilutions. Total RNA was purified from clonal population by Trizol extraction according to the manufacturer’s protocol. The experiment had 7 RNA samples collected from 2 clones, 6 samples from clone 1 expressing VSP GL50803_00113797 (3 for control and 3 for antibody treatment) and 1 sample from clone 2 expressing the VSP pair GL50803_00112208 and GL50803_00d112208. Ribosomal RNA was depleted in all samples and strand-specific Illumina sequencing libraries were constructed. RNA sequencing was performed as a paid service by Genehub.com. The libraries were sequenced as 2×150 bp paired-end reads on an Illumina MiSeq instrument, and 3.1-4.3 million reads were subsequently generated for each library. Raw sequence reads from Illumina were submitted to the NCBI Sequence Read Archive, with SRA accession numbers SRR17933280 to SRR17933282 and SRR17933276 for control samples, and SRR17933277 to SRR17933279 for antibody-treated samples. The sequences can also be accessed from the BioProject PRJNA804420. Genome index was generated using STARv2.5.4b with default parameters except for --genomeSAindexNbases. According to the STAR manual, for small genomes, the parameter --genomeSAindexNbases must be scaled down, with a typical value of min(14, log_2_(GenomeLength)/2 - 1). Given that *G. lamblia* genome is 12.6 Mb in length, this parameter was set to 11. Quality control of raw reads was performed using FastQC v0.11.5. If needed, trimming of low quality base calls (below 30) and adapter sequences was performed with TrimGalore v0.6.4 and Cutadapt v1.15, in order to improve mapping efficiency. Sequences shorter than 70 bp were removed. Reads that passed quality criteria were aligned to the latest reference genome using STAR v2.5.4b with default parameters except for: (1) --alignIntronMax (maximum intron size), which was set to 1 because, with few exceptions, *G. lamblia* genes lack introns, and (2) --outFilterMultimapNmax (max number of multiple alignments allowed for a read), which was set to 2 to account for duplicated genes. On average, ∼80% of all sequenced reads could be mapped to a single location in the genome assembly. After mapping, reads were assigned to genomic features using FeatureCounts v2.0.0. Multi-overlapping and reads not overlapping with any features in the annotation file were not counted. Since many VSP genes are duplicated, two strategies for counting multimapping reads were used. In strategy UM, multimapping reads were not counted, only unique mapping reads were considered, whereas in strategy 2M, multimapping reads were fractionally counted (each alignment carrying 1/2 count, since the parameter --outFilterMultimapNmax was previously set to 2 in the alignment step). Using 2M strategy, it was possible to incorporate approximately 3.35% extra reads mapping to two locations in the genome. Sense and antisense counts for each gene in each library were then normalized to transcript per million (TPM). Commands used are indicated in <u>S1 file</u>. The same procedure used for analyzing our own samples was used for analyzing several transcriptomic datasets deposited in SRA. Differential expression analysis was done using edgeR v3.28 package. Counts-per-million were calculated for each gene to standardize for differences in library-size. Genes that do not have a worthwhile number of counts in any sample were filtered out of downstream analyses. Therefore, filtering was carried out to retain genes with a baseline expression level of at least 10 cpm in 3 or more samples. For each data set, TMM normalization was then applied to account for the compositional biases. The aim of the analysis was to detect genes differentially expressed between antibody-treated and control samples, adjusting for any differences between samples. This is a paired design in which each sample receives both the control and the treatment. Therefore, it is possible to compare the treatment to the control for each sample separately; thus, baseline differences between samples are subtracted out. All plots, images and statistical analyses were performed in R (v4.0.3) and GraphPad Prism (v9.00), and were later compiled in Adobe Illustrator.

### Proteomic analysis

Data were processed using MaxQuant version 2.0.1.0 with the parameters indicated in the corresponding PRIDE projects. In general, however, the following parameters were used: Enzyme, Trypsin/P; Database, NCBI *Giardia lamblia* RefSeq 2.1 proteome (concatenated forward and reverse plus common contaminants); Fixed modification, Carbamidomethyl (C); Variable modifications, Oxidation (M), Acetyl (N-term), Pyro-Glu (N-term Q), Deamidation (N/Q); Mass values, Monoisotopic; Peptide Mass Tolerance, 10 ppm; Fragment Mass Tolerance, 0.02 Da; Max Missed Cleavages, 2. Data were filtered using at 1% protein and peptide FDR and requiring at least two unique peptides per protein.

## Acknowledgments

Authors thank Dr. Albano H. Tenaglia for preparing the clones for transcriptomic analysis. This work was supported by grants FONCYT (PICT-13469, PICT-2703, PICT-E 0234, and PICT-2116), CONICET (D4408), and UCC (80020150200144CC) of Argentina to H.D.L.

## Author contributions

MRW and CRM performed most of the sequences analyses. MRW contributed the transcriptomic analysis. LAL contributed the proteomic analyses, the cloning and the immunofluorescence assays. MRW, CRM and LAL verified each other results. AS generated monoclonal antibodies. VMB contributed with *in silico* structural analysis and reproduction of the transcriptomic analyses. EAF and HDL supervised the bioinformatic analyses. SGS and JJH contributed with PAS analyses. HDL conceived the project and designed the experiments. All authors analyzed the data. MRW, CRM, EAF, SGS and HDL wrote the paper. All authors read and commented on the manuscript.

