## Supplemental Figures and Tables for "Characterization of Cysteine-rich protein families of *Giardia lamblia* and their role during antigenic variation"

**Supporting information** (tables and figures in order of appearance in the text of the manuscript)

**S1 table. Cys-rich genes.** Detailed information of all the gene and protein IDs, chromosome/scaffold localization, annotated name, nucleotide and protein sequence and length, presence of INR, cysteine content, number of CXC, CXXC and CXXXC motifs, predicted TMD, CT and SP for each VSP, CRMP and SCRP. (See separate Excel table)

**S2 table. Repeats in all VSPs according to Tandem Repeat Finder.** In the first tab, the VSP with repeats and de number of the repeats in each one is shown. In the second tab, the consensus sequences in nucleotides of each VSP with repeat is described, including the consensus sequences of the repeat belonging to each particular VSP. (See separate Excel document)

**S1 fig. Repeats in VSPs.** The amino acid sequence of each VSP with their corresponding tandem repeats as found using RADAR is shown. Note the high similarity of each repeat belonging to the same VSP. Some VSP share the sequences of the repeats. (See separate PDF document)

**S3 table. Metal binding properties of Cys-rich genes.** Prediction of free cysteines and cysteines that can bind iron, zinc and cadmium in each Cyr-rich group. (See separate Excel table)

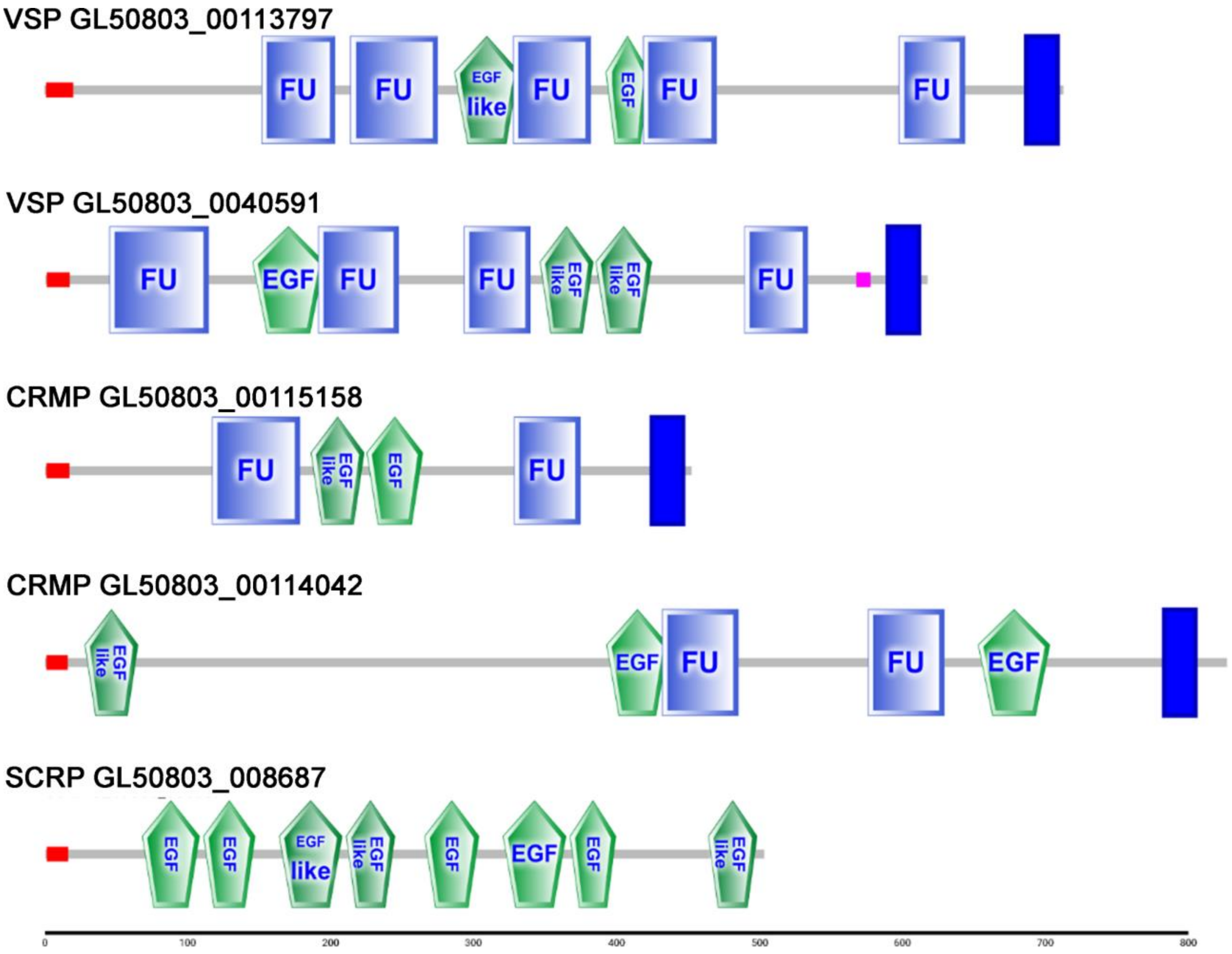


**S2 fig. Examples of domains predicted by SMART in selected Cys-rich proteins.** Red rectangle: signal peptide. Blue rectangle: transmembrane domain. Magenta rectangle: low complexity region. FU: Furin-like repeats (accession SM000261). EGF: Epidermal growth factor-like domain (accession SM000181). EGF-like: EGF domain, unclassified subfamily (accession SM000001). The figure indicates the gene ID and the group to which each sequence belongs (VSP, CRMP or SCRP). The line at the bottom indicates the amino acid range.

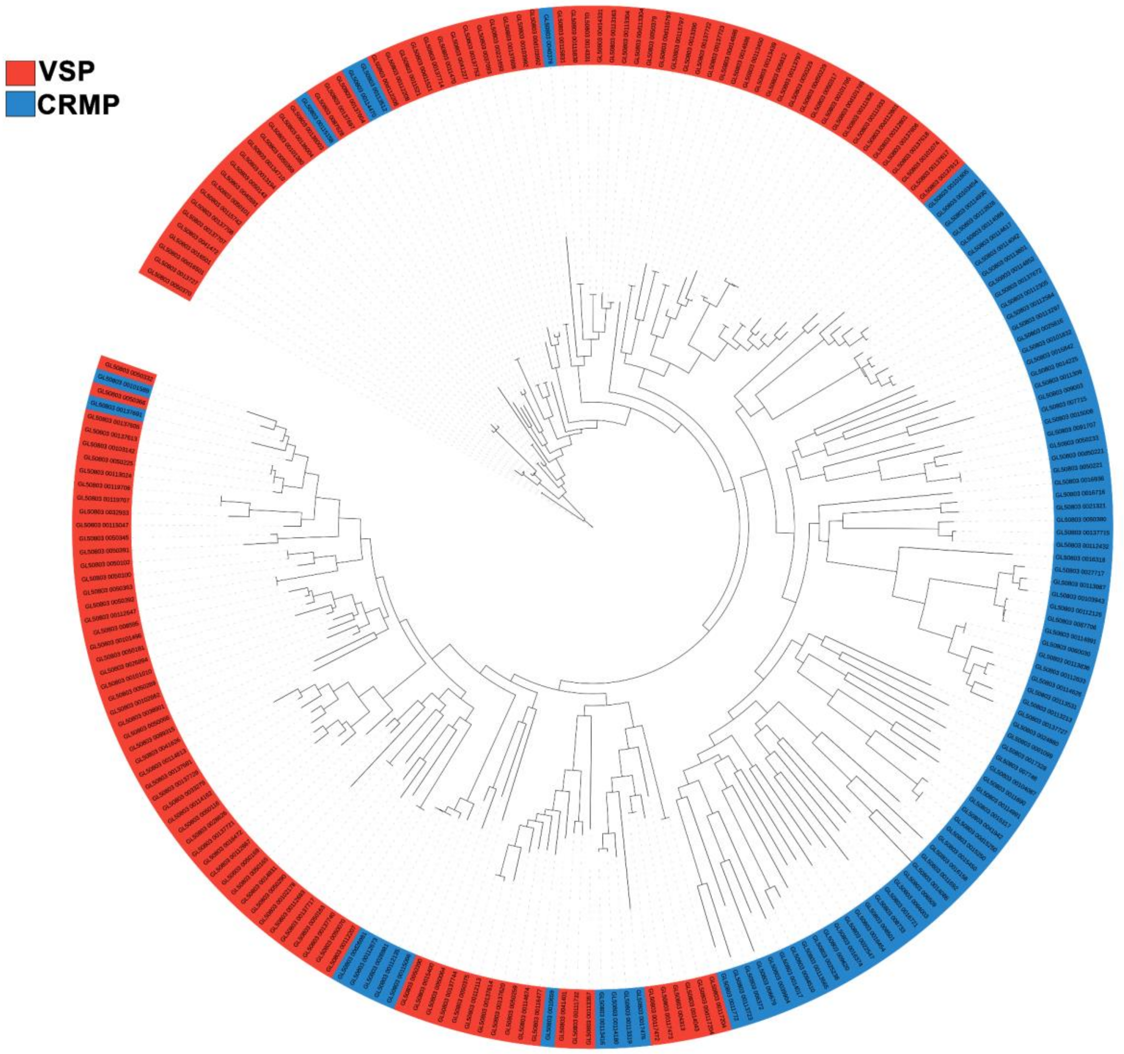


**S3 fig. Tree of VSP and HCMP.** A phylogenetic analysis was performed using multiple sequence alignment by MAFFT and RAxML and the output tree assembled using iTOL. A clear separation of the VSP and CRMP groups can be observed, except for those proteins that share common characteristics.

**S4 table. VSP and CRMP duplicated pairs.** Information about gene ID of duplicated pairs, orientation, chromosomal location, intergenic region length and genes/pseudogenes found in between.

| Chr | Duplicated Genes | Genes | Orientation | Gene(s) in between | Intergenic region | Length |
|---|---|---|---|---|---|---|
| 1 | GL50803_00d112208 and GL50803_00112208 | VSP | → ← | None | VSP like region (*2) (*5) | 2,648 |
| 1 | GL50803_00d115797 and GL50803_00115797 | VSP | → ← | None | VSP like region (*2) (*5) | 2,649 |
| 1 | GL50803_00112673 and GL50803_00d26981 (*) | CRMP | ← ← | Many genes | VSP like region<br>Hypothetical proteins<br><br>Dynein binding protein<br>Phenylalanyl-tRNA synthase beta chain<br>Chaperonin subunit epsilon CCT epsilon (Ccte)<br>Phenylalanyl-tRNA synthase beta chain<br>TCP-1 chaperonin subunit epsilon<br>Papain family peptidase<br>Translation initiation factor eIF3 subunit<br>ADP-sugar diphosphatase<br>Mechanosensory abnormality MEC-17-like protein<br>Translation elongation factor 1-beta<br>Ribosome biogenesis regulatory protein (RRS1)<br>Myb-like DNA-binding domain-containing protein<br>Ribosomal protein S18 | 4,0376 |
| 2 | GL50803_00d11521 and GL50803_0011521 | VSP | → ← | None | VSP like region (*3) | 2,747 |
| 2 | GL50803_00d117204 and GL50803_00117204 | VSP | → ← | None | VSP like region (*5) | 2,720 |
| 2 | GL50803_00117472 and GL50803_00117473 | VSP | ← → | GL50803_101534 NEK Kinase | None | |
| 2 | GL50803_0050359 and GL50803_00134710 | | ← → | None | None | 12,440 |
| 3 | GL50803_00115830 and GL50803_00115831 | VSP | → ← | None | VSP like region (*3) | 2,813 |
| 3 | GL50803_00119706 and GL50803_00119707 | VSP | ← → | Pseudogene | Two retrotransposon relics (*4) | 3,301 |
| 3 | GL50803_00136003 and GL50803_00136004 | VSP | → ← | None | VSP like region (*5) | 2,686 |
| 4 | GL50803_0050221 and GL50803_00d50221 | CRMP | ← → | 2 duplicated pVSP, other pseudogene and a kinase NEK (GL50803_00113030) | Kinases NEK<br>Two retrotransposon relics<br>VSP like region | 1,5462 |
| 4 | GL50803_00d103992 and GL50803_00103992 | VSP | → ← | None | VSP like region (*5) | 2,758 |

| | | | | | | |
|---|---|---|---|---|---|---|
| 4 | GL50803_00111933 and GL50803_00111936 | VSP | ⟵ ⟶ | Pseudogene | Two retrotransposon relics (*4) | 3,846 |
| 4 | GL50803_00d112801 and GL50803_00112801 | VSP | ⟵ ⟶ | Pseudogene | Two retrotransposon relics | 6,264 |
| 4 | GL50803_0016501 and GL50803_00d16501 | VSP | ⟶ ⟵ | None | VSP like region (*3) | 2,526 |
| 4 | GL50803_0050229 and GL50803_00d50229 | VSP | ⟵ ⟶ | Pseudogene | Two retrotransposon relics | 3,242 |
| 5 | GL50803_00d113304 and GL50803_00113304 | VSP | ⟶ ⟵ | Many genes and pseudogenes | CRMP and pseudo Cys-rich | 72,283 |
| 5 | GL50803_00137714 and GL50803_0011470 | VSP | ⟶ ⟵ | None | VSP like region (*5) | 2,669 |
| 5 | GL50803_00137708 and GL50803_00137707 | VSP | ⟶ ⟵ | None | VSP like region (*3) | 2,777 |
| 5 | GL50803_00137722 and GL50803_00137723 | VSP | ⟵ ⟶ | None | VSP like region (*5) | 2,513 |
| 5 | GL50803_00d14331 and GL50803_0014331 | VSP | ⟶ ⟵ | None | VSP like region (*5) | 2,637 |
| 5 | GL50803_0014586 and GL50803_00d14586 | VSP | ⟶ ⟵ | None | Very similar region intergenic region between non duplicated VSP genes GL50803_0050392 and GL50803_00112647 | 2,689 |
| 5 | GL50803_00d15250 and GL50803_0015250 | CRMP | ⟵ ⟶ | Many genes and pseudogenes, including other HCMP (GL50803_00113836) | VSP like region<br>N-acetyltransferase-like protein<br>Hypotheticals proteins<br>Coiled-coil protein (GL50803_0095653)<br>Ankyrin repeat protein 1 and protein 2<br>Tubulin tyrosine ligase (GL50803_0095661)<br>Nuclear ATP/GTP-binding protein (GL50803_0017025)<br>Multidrug MFS transporter (GL50803_008444)<br>Kinase, NEK (GL50803_008445)<br>Las1-like protein | 55,063 |

(*) Note that GL50803_00112673 and GL50803_00d26981 are the duplicated pair and not GL50803_0026981 and GL50803_00d26981 as the Gene ID nomenclature indicates.

(*2) Note that the intergenic regions of these two clusters of duplicated VSPs are identical.

(*3, *4, *5, *6) Note that the intergenic regions of these clusters of duplicated genes are very similar.

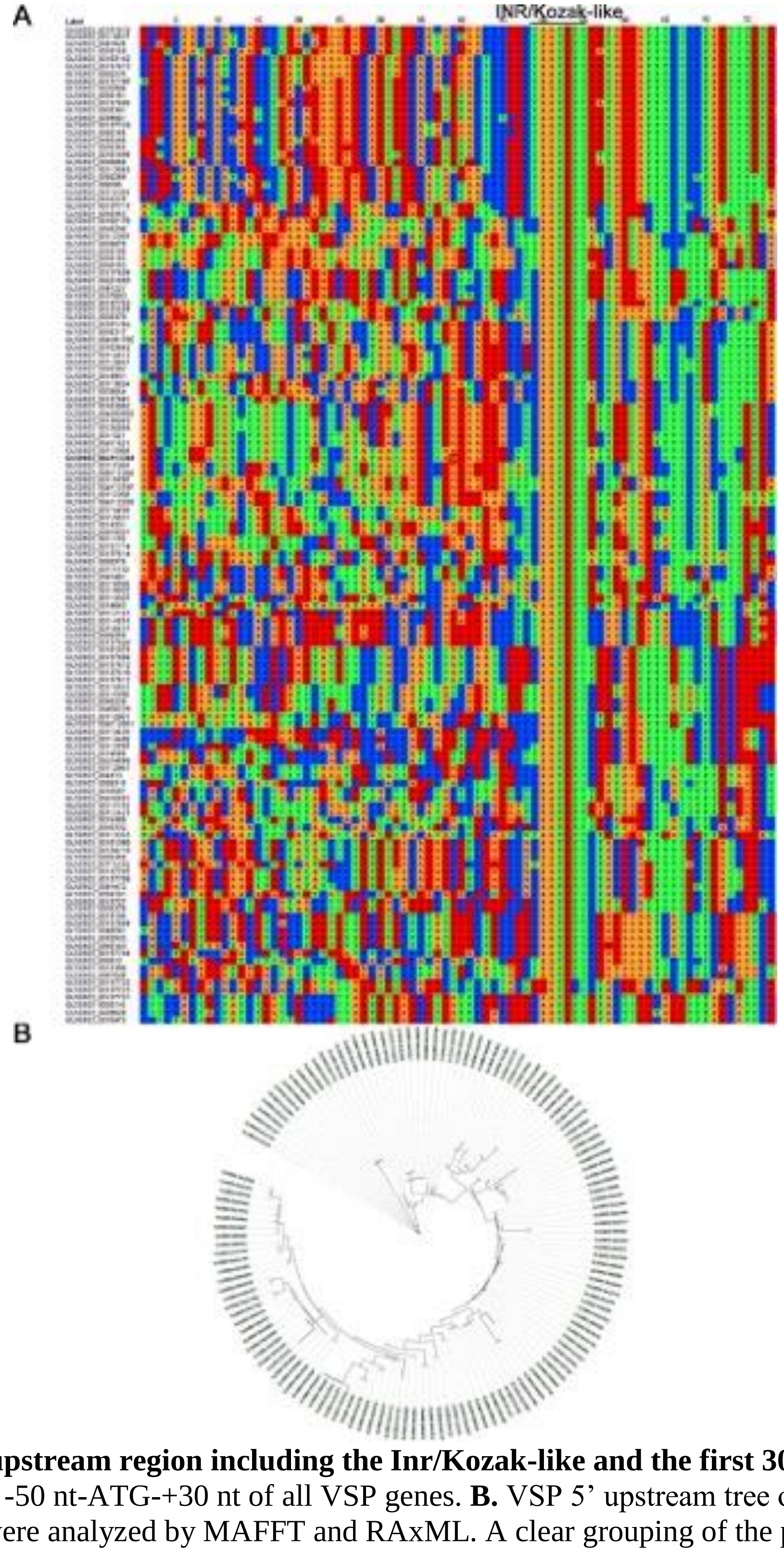


**S4 fig. VSP 5'-upstream region including the Inr/Kozak-like and the first 30 nt. A.** MAFFT alignment of the -50 nt-ATG-+30 nt of all VSP genes. **B.** VSP 5' upstream tree of -50 nt-ATG-+30 nt of VSP were analyzed by MAFFT and RAxML. A clear grouping of the portions of the genes can be observed.

**S5 table. *Giardia* genes (other than VSPs) with INR.** Gene and protein ID, annotated name and chromosomal localization of the few genes, apart from VSP genes, in which INR can be detected.

| gene id | protein id | annotated name | INR | Chromosome |
|---|---|---|---|---|
| GL50803_0016722 | XP_001704332.2 | Xaa-Pro dipeptidase | TAATGTT | NC_051856.1 |
| GL50803_0017506 | XP_001709082.1 | hypothetical protein | CAATGTT | NC_051856.1 |
| GL50803_00101589 | XP_001704555.1 | HCMP | TAATGTT | NC_051857.1 |
| GL50803_005062 | XP_001704724.1 | Transcription factor b2 | TAATGTT | NC_051858.1 |
| GL50803_0015035 | XP_001707360.2 | Kinase, NEK | TAATGTT | NC_051859.1 |
| GL50803_0012059 | XP_001707290.2 | hypothetical protein | CAATGTT | NC_051859.1 |
| GL50803_0017577 | XP_001708338.1 | hypothetical protein | CAATGTT | NC_051859.1 |
| GL50803_002362 | XP_037902118.1 | hypothetical protein | TAATGTT | NC_051859.1 |
| GL50803_ 0091504 | XP_037902138.1 | hypothetical protein | CAATGTT | NC_051860.1 |
| GL50803_008650 | XP_001704194.2 | hypothetical protein | TAATGTT | NC_051860.1 |
| GL50803_0017104 | XP_001706769.1 | hypothetical protein | CAATGTT | NC_051860.1 |
| GL50803_008722 | XP_001704771.1 | Myb 1-like protein | TAATGTT | NC_051860.1 |
| GL50803_0017447 | XP_037902382.1 | hypothetical protein | TAATGTT | NC_051860.1 |

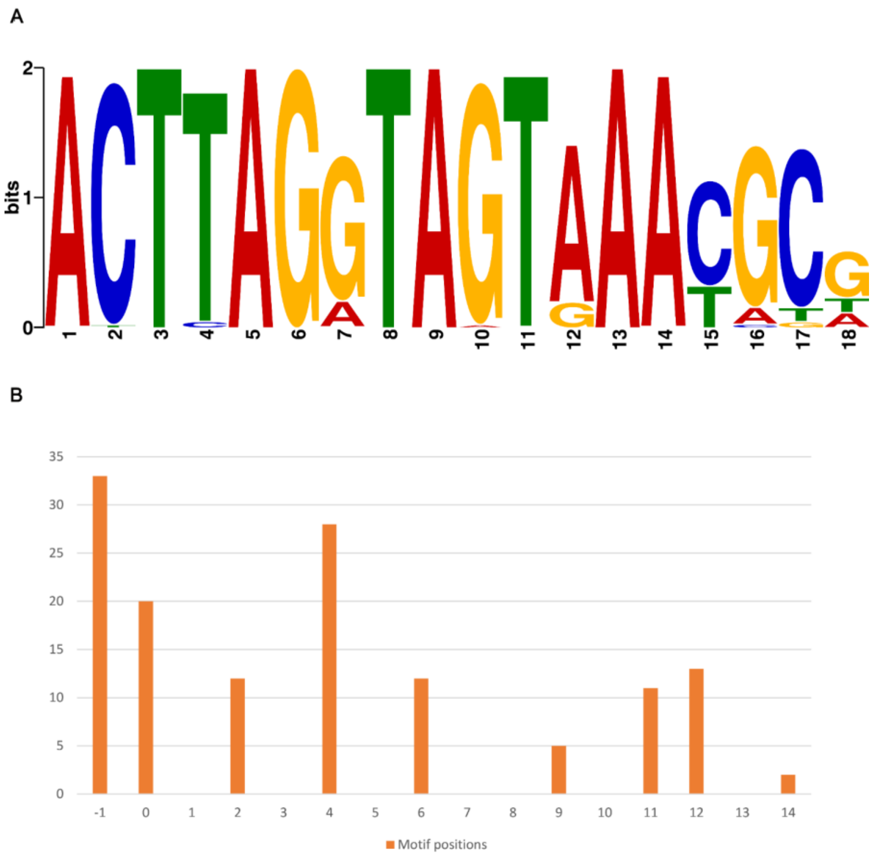


**S5 fig. An extended polyadenylation adjacent sequence (PAS) defines the VSP gene family.** **(A)** The PAS is found downstream of all VSP genes (n=136) *in G. intestinalis* WB. (**B)** The PAS is found at discrete locations at -1 to +14 nucleotides from the end of the VSP genes.}

**S6 table. Genome-wide search for the extended PAS.** The extended PAS motif GACTTAGGTAGTRAAYGCKK was searched in the 3'UTR region (stop codon + 50 bp) of all genes (including pseudogenes) in the *Giardia* WB genome using the Fimo software from the MEME suite. The genes with 3'UTR matches are reported with start, end, strand, significance (p-value and q-value), and annotation. Manually curated hits are indicated by colors significant (green) and false positive (red). (See separate Excel table)

**S7 table. Cys-rich pseudogenes.** Detailed information about Cys-rich pseudogenes type I and type II defined in this work, including gene ID, annotated name, chromosome/scaffold localization, nucleotide sequence and length, among other features. (See separate Excel table)

**S8 table. Cys-rich proteins across isolates, assemblages and species.** Detailed information about Cys-rich genes of G. *muris*, P15, GS (GS and GS-B) and DH; including gene ID, annotated name, possible Cys-rich group, nucleotide and amino-acid sequence and length, among other features. (See separate Excel table)

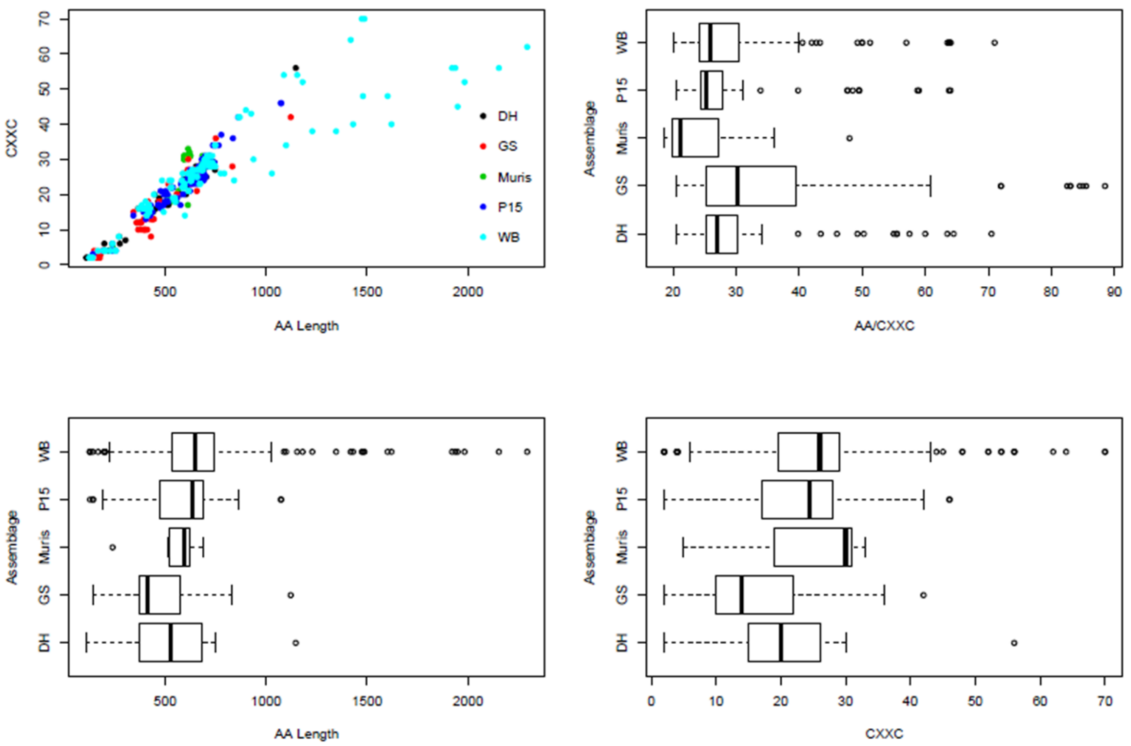


**S6 fig. Cys-rich proteins across isolates, assemblages and species.** Amino acid (aa) length vs. number of CXXC, aa/CXXC ratio, length and number of CXXC motifs between assemblages.

**S9 table. Transcript per million (TPM) of the RNA-seq experiment described in this work.** Sense and antisense TPM for each transcript in each sample, using both UM and 2M strategies for counting. (See separate Excel table)

**S10 table. Differential gene expression (DGE) analysis.** The gene-wise tests shown are for Clone1T (treated with mAb) vs. Clone1 (control), adjusting for baseline differences between batches. EdgeR with quasi-likelihood F-test was used. Green rows indicate differentially expressed genes with FDR < 0.05. (See separate Excel table)

**S11 table. SRA accession number of transcriptomic raw data analyzed.** The corresponding SRA accession number of each sample analyzed is shown, together with the reference.

| Sample | SRA accession | Reference |
|---|---|---|
| Clone1-rep1 | SRR17933282 | This work |
| Clone1-rep2 | SRR17933281 | This work |
| Clone1-rep3 | SRR17933280 | This work |
| Clone1T-rep1 | SRR17933279 | This work |
| Clone1T-rep2 | SRR17933278 | This work |
| Clone1T-rep3 | SRR17933277 | This work |
| Clone2-rep1 | SRR17933276 | This work |
| Clone3-rep1 | SRR10063826 | doi: 10.3389/fgene.2020.00913 |
| Clone3-rep2 | SRR10063830 | doi: 10.3389/fgene.2020.00913 |
| Clone3-rep3 | SRR10063831 | doi: 10.3389/fgene.2020.00913 |
| Clone3-rep4 | SRR10063832 | doi: 10.3389/fgene.2020.00913 |
| Population1-rep1 | SRR10919358 | doi: 10.3389/fgene.2020.00913 |
| Population1-rep2 | SRR10919350 | doi: 10.3389/fgene.2020.00913 |
| Population1-rep3 | SRR10919356 | doi: 10.3389/fgene.2020.00913 |
| Population1-rep4 | SRR10919352 | doi: 10.3389/fgene.2020.00913 |
| SC1 | SRR9222556 | doi.org/10.1186/s12864-020-06858-7 |
| SC2 | SRR9222558 | doi.org/10.1186/s12864-020-06858-7 |
| SC3 | SRR9222577 | doi.org/10.1186/s12864-020-06858-7 |
| SC4 | SRR9222587 | doi.org/10.1186/s12864-020-06858-7 |
| SC5 | SRR9222595 | doi.org/10.1186/s12864-020-06858-7 |
| SC6 | SRR9222598 | doi.org/10.1186/s12864-020-06858-7 |

**S12 table. Selected housekeeping genes.** Gene ID and annotated name of selected housekeeping genes.

| gene_id | annotated name |
|---|---|
| GL50803_00101291 | Beta-tubulin 1 |
| GL50803_00102515 | RNA-directed RNA polymerase |
| GL50803_0010311 | Ornithine carbamoyl transferase |
| GL50803_00103676 | Alpha-tubulin 1 |
| GL50803_0010570 | Peptidyl-prolyl cis-trans isomerase |
| GL50803_0010623 | Phosphoenolpyruvate carboxy Kinase |
| GL50803_0010885 | Amylo-alpha-1,6-glucosidase |
| GL50803_0011043 | Fructose-bisphosphate aldolase |
| GL50803_0011118 | Enolase |
| GL50803_00112079 | Alpha-tubulin |
| GL50803_00112103 | Arginine deiminase |
| GL50803_00112304 | Elongation factor 1-alpha |
| GL50803_00112312 | Elongation factor 1-alpha |
| GL50803_0011301 | Nucleoside diphosphate kinase |
| GL50803_00114609 | Pyruvate-flavodoxin oxidoreductase |
| GL50803_00114787 | Alpha-7.3 giardin |
| GL50803_0011654 | Alpha-1 giardin |
| GL50803_0012102 | Elongation factor 1-gamma |
| GL50803_0012150 | Alanine aminotransferase |
| GL50803_0013350 | Alcohol dehydrogenase 3 |
| GL50803_0013608 | Acetyl-CoA synthetase |
| GL50803_0014058 | CAMP-specific 3',5'-cyclic phosphodiesterase 4B |
| GL50803_0014285 | Malate dehydrogenase |
| GL50803_0014993 | Diphosphate-fructose-6-phosphate 1-phosphotransferase |
| GL50803_0015106 | Importin beta-3 subunit |
| GL50803_0015574 | Alanyl dipeptidyl peptidase |
| GL50803_0015832 | Aminoacyl-histidine dipeptidase |
| GL50803_0016125 | FAD-dependent glycerol-3-phosphate dehydrogenase |
| GL50803_0016343 | Median body protein |
| GL50803_0016453 | Carbamate kinase |
| GL50803_0016549 | Uridine kinase |
| GL50803_0016667 | Acyl-CoA synthetase |
| GL50803_0017063 | Pyruvate-flavodoxin oxidoreductase |
| GL50803_0017121 | Bip |
| GL50803_0017143 | Pyruvate kinase |
| GL50803_0017150 | NADPH oxidoreductase |
| GL50803_0017153 | Alpha-11 giardin |
| GL50803_0017163 | Peptidyl-prolyl cis-trans isomerase B |
| GL50803_0017230 | Gamma giardin |
| GL50803_0017254 | phosphoglucomutase |

| | |
|---|---|
| GL50803_0017327 | Xaa-Pro dipeptidase |
| GL50803_0017570 | Elongation factor 2 |
| GL50803_0017587 | CTP synthase |
| GL50803_002101 | Cytidine deaminase |
| GL50803_0021423 | Beta adaptin |
| GL50803_0021942 | NADP-specific glutamate dehydrogenase |
| GL50803_002452 | Ornithine cyclodeaminase |
| GL50803_0028234 | Adenylate kinase |
| GL50803_003206 | Pyruvate kinase |
| GL50803_0032658 | Plasma membrane calcium-transporting ATPase 2 |
| GL50803_003287 | Acetyl-CoA acetyltransferase |
| GL50803_003331 | Malate dehydrogenase |
| GL50803_0033769 | NADH oxidase |
| GL50803_0035180 | GTOR |
| GL50803_003593 | Alcohol dehydrogenase 3 |
| GL50803_003861 | Alcohol dehydrogenase 3 |
| GL50803_0040244 | Emp24/gp25L/p24 family/GOLD protein |
| GL50803_0040817 | Actin |
| GL50803_004812 | Beta-giardin |
| GL50803_004964 | Mevalonate kinase |
| GL50803_005333 | Calmodulin |
| GL50803_006148 | Alanyl dipeptidyl peptidase |
| GL50803_006226 | Glycogen phosphorylase |
| GL50803_006687 | Glyceraldehyde 3-phosphate dehydrogenase |
| GL50803_007195 | Glutamate synthase |
| GL50803_007260 | Aldose reductase |
| GL50803_007556 | Acid phosphatase |
| GL50803_008217 | Uridine kinase |
| GL50803_008407 | Aminoacyl-histidine dipeptidase |
| GL50803_0086511 | Acyl-CoA synthetase |
| GL50803_008822 | 2,3-bisphosphoglycerate-independent phosphoglycerate mutase |
| GL50803_008826 | Glucokinase |
| GL50803_009008 | Aldose reductase |
| GL50803_0090872 | Phosphoglycerate kinase |
| GL50803_009115 | Glucose-6-phosphate isomerase |
| GL50803_0093358 | Bifunctional acetaldehyde-CoA/alcohol dehydrogenase |
| GL50803_009704 | Transketolase |
| GL50803_009779 | Uridine phosphorylase 1 |
| GL50803_009909 | Pyruvate, phosphate dikinase |

**S13 table. Cys-rich proteins with evidence of translation.** VSP, CRMP and SCRP proteins with peptides (either unique or not unique) detected in all of the following PRIDE projects: PXD017597, PXD004398, PXD000452, PXD002398, PXD007183 and PXD022565. (See separate Excel table)

### Supporting method

**S1 file. Transcriptomic analysis workflow.** Systematic pipeline from raw reads to TPM of each transcript in each library. (See separate PDF document)
