## Supplemental Method file 1 for "Characterization of Cysteine-rich protein families of *Giardia lamblia* and their role during antigenic variation"

### Transcriptomic analysis workflow

All of the following steps were run on a Windows Subsystem Linux machine. You could run it on Linux as well.

##### Download genome assembly

- Download genome assembly from RefSeq. BioProject: PRJNA1439. (https://www.ncbi.nlm.nih.gov/assembly/GCF_000002435.2).

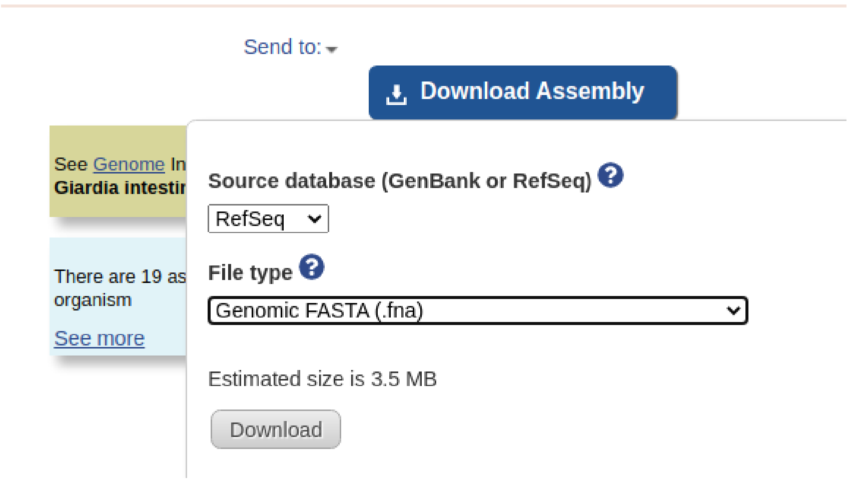


- This will download the following file: genome_assemblies_genome_fasta.tar.
- Extract the files in the compress folder. At the end of this step, you should conserve the file GCF_000002435.2_UU_WB_2.1_genomic.fna.
- Then, download the annotation file from RefSeq in .gtf format.

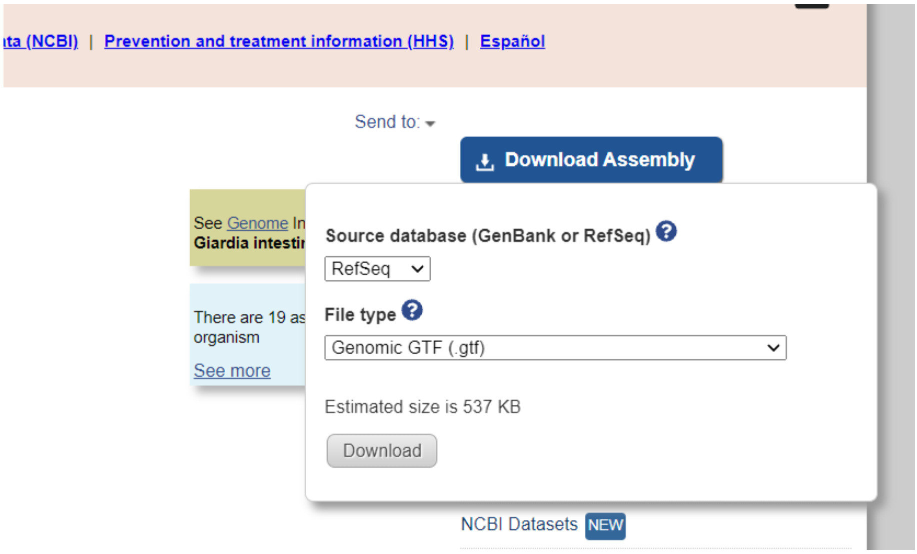

- This will download the file genome_assemblies_genome_gtf.tar.
- Extract the files in the compress folder. At the end of this step, you should conserve the file GCF_000002435.2_UU_WB_2.1_genomic.gtf.

- Store both files (.fna and .gtf) in your favorite directory, we will called it *Giardia*.

  ./Giardia/GCF_000002435.2_UU_WB_2.1_genomic.fna

  ./Giardia/GCF_000002435.2_UU_WB_2.1_genomic.gtf

**Build the genome index**

- Go to ./Giardia directory and type the following commands in your terminal. Make sure you have STAR in the PATH.

```
mkdir ./genomeDir

STAR --runThreadN 8 --runMode genomeGenerate --genomeDir ./genomeDir --genomeSAindexNbases 11 --genomeFastaFiles GCF_000002435.2_UU_WB_2.1_genomic.fna --sjdbGTFfile GCF_000002435.2_UU_WB_2.1_genomic.gtf
```

```
macarw@DESKTOP-Q9UPIUR:/mnt/c/Users/MacaRW/Desktop/Giardia$ ls
GCF_000002435.2_UU_WB_2.1_genomic.fna  GCF_000002435.2_UU_WB_2.1_genomic.gtf
macarw@DESKTOP-Q9UPIUR:/mnt/c/Users/MacaRW/Desktop/Giardia$ mkdir ./genomeDir
macarw@DESKTOP-Q9UPIUR:/mnt/c/Users/MacaRW/Desktop/Giardia$ ls
GCF_000002435.2_UU_WB_2.1_genomic.fna  GCF_000002435.2_UU_WB_2.1_genomic.gtf  genomeDir
macarw@DESKTOP-Q9UPIUR:/mnt/c/Users/MacaRW/Desktop/Giardia$ STAR --runThreadN 8 --runMode genomeGenerate --genomeDir ./g
enomeDir --genomeSAindexNbases 11 --genomeFastaFiles GCF_000002435.2_UU_WB_2.1_genomic.fna --sjdbGTFfile GCF_000002435.2
_UU_WB_2.1_genomic.gtf
Oct 25 20:50:38 ..... started STAR run
Oct 25 20:50:38 ... starting to generate Genome files
Oct 25 20:50:38 ... starting to sort Suffix Array. This may take a long time...
Oct 25 20:50:38 ... sorting Suffix Array chunks and saving them to disk...
Oct 25 20:50:45 ... loading chunks from disk, packing SA...
Oct 25 20:50:46 ... finished generating suffix array
Oct 25 20:50:46 ... generating Suffix Array index
Oct 25 20:50:47 ... completed Suffix Array index
Oct 25 20:50:47 ..... processing annotations GTF
Oct 25 20:50:48 ..... inserting junctions into the genome indices
Oct 25 20:50:48 ... writing Genome to disk ...
Oct 25 20:50:48 ... writing Suffix Array to disk ...
Oct 25 20:50:49 ... writing SAindex to disk
Oct 25 20:50:49 ..... finished successfully
macarw@DESKTOP-Q9UPIUR:/mnt/c/Users/MacaRW/Desktop/Giardia$
```

- Note that genome index is generated using STAR with default parameters except for --genomeSAindexNbases. According to the STAR manual, for small genomes, this must be scaled down, with a typical value of min(14, $\log_2$(GenomeLength)/2 - 1). Given that *G. lamblia* genome is 12.6 Mb in length, this parameter is set to 11.
- You can adjust --runThreadN parameter according to the number of available cores on your server node.

- This should end up with the index files in Giardia/genomeDir directory (You can move the Log.out file to /genomeDir).

```
macarw@DESKTOP-Q9UPIUR:/mnt/c/Users/MacaRW/Desktop/Giardia$ ls
GCF_000002435.2_UU_WB_2.1_genomic.fna  GCF_000002435.2_UU_WB_2.1_genomic.gtf  Log.out  genomeDir
macarw@DESKTOP-Q9UPIUR:/mnt/c/Users/MacaRW/Desktop/Giardia$ mv Log.out ./genomeDir/
macarw@DESKTOP-Q9UPIUR:/mnt/c/Users/MacaRW/Desktop/Giardia$ ls
GCF_000002435.2_UU_WB_2.1_genomic.fna  GCF_000002435.2_UU_WB_2.1_genomic.gtf  genomeDir
macarw@DESKTOP-Q9UPIUR:/mnt/c/Users/MacaRW/Desktop/Giardia$ cd genomeDir/
macarw@DESKTOP-Q9UPIUR:/mnt/c/Users/MacaRW/Desktop/Giardia/genomeDir$ ls
Genome   SAindex         chrNameLength.txt  exonInfo.tab          sjdbInfo.txt              transcriptInfo.tab
Log.out  chrLength.txt   chrStart.txt       geneInfo.tab          sjdbList.fromGTF.out.tab
SA       chrName.txt     exonGeTrInfo.tab   genomeParameters.txt  sjdbList.out.tab
macarw@DESKTOP-Q9UPIUR:/mnt/c/Users/MacaRW/Desktop/Giardia/genomeDir$ |
```

For the remaining steps, we will use two paired-end fastq files (R1.fq.gz and R2.fq.gz located in /Giardia directory) as an example of how to trim reads, align reads to the genome, count number of reads per gene and convert raw reads to transcript per million (TPM). You can download those files from here.

**Quality check**

- A thorough quality check of the fastq data is performed with FastQC. Run the following command in your terminal on the right directory:

```
for file in *fq.gz; do fastqc $file; done
```

```
macarw@DESKTOP-Q9UPIUR:/mnt/c/Users/MacaRW/Desktop/Giardia$ ls
GCF_000002435.2_UU_WB_2.1_genomic.fna  GCF_000002435.2_UU_WB_2.1_genomic.gtf  R1.fq.gz  R2.fq.gz  genomeDir
macarw@DESKTOP-Q9UPIUR:/mnt/c/Users/MacaRW/Desktop/Giardia$ for file in *fq.gz; do fastqc $file; done
Started analysis of R1.fq.gz
Approx 5% complete for R1.fq.gz
Approx 10% complete for R1.fq.gz
Approx 15% complete for R1.fq.gz
Approx 20% complete for R1.fq.gz
Approx 25% complete for R1.fq.gz
Approx 30% complete for R1.fq.gz
Approx 35% complete for R1.fq.gz
Approx 40% complete for R1.fq.gz
Approx 45% complete for R1.fq.gz
Approx 50% complete for R1.fq.gz
Approx 55% complete for R1.fq.gz
Approx 60% complete for R1.fq.gz
Approx 65% complete for R1.fq.gz
Approx 70% complete for R1.fq.gz
Approx 75% complete for R1.fq.gz
Approx 80% complete for R1.fq.gz
Approx 85% complete for R1.fq.gz
Approx 90% complete for R1.fq.gz
Approx 95% complete for R1.fq.gz
Analysis complete for R1.fq.gz
Started analysis of R2.fq.gz
Approx 5% complete for R2.fq.gz
Approx 10% complete for R2.fq.gz
Approx 15% complete for R2.fq.gz
Approx 20% complete for R2.fq.gz
Approx 25% complete for R2.fq.gz
Approx 30% complete for R2.fq.gz
Approx 35% complete for R2.fq.gz
Approx 40% complete for R2.fq.gz
Approx 45% complete for R2.fq.gz
Approx 50% complete for R2.fq.gz
Approx 55% complete for R2.fq.gz
Approx 60% complete for R2.fq.gz
Approx 65% complete for R2.fq.gz
Approx 70% complete for R2.fq.gz
Approx 75% complete for R2.fq.gz
Approx 80% complete for R2.fq.gz
Approx 85% complete for R2.fq.gz
Approx 90% complete for R2.fq.gz
Approx 95% complete for R2.fq.gz
Analysis complete for R2.fq.gz
macarw@DESKTOP-Q9UPIUR:/mnt/c/Users/MacaRW/Desktop/Giardia$
```

- You should end up with two extra files for each fastq file (e.g. for R1.fq.gz you obtain R1_fastqc.html and R1_fastqc.zip). The .html can be open in a browser of your preference, in order to visualize the quality of raw reads.

```
macarw@DESKTOP-Q9UPIUR:/mnt/c/Users/MacaRW/Desktop/Giardia$ ls
GCF_000002435.2_UU_WB_2.1_genomic.fna  R1.fq.gz         R1_fastqc.zip  R2_fastqc.html  genomeDir
GCF_000002435.2_UU_WB_2.1_genomic.gtf  R1_fastqc.html  R2.fq.gz       R2_fastqc.zip
macarw@DESKTOP-Q9UPIUR:/mnt/c/Users/MacaRW/Desktop/Giardia$
```

#### Trimming of reads

- If needed, some quality improvement processing is done with TrimGalore.
- Low-quality base calls (below 30) are trimmed off from the 3' end of the reads. This efficiently removed poor quality portions of the reads that may lead to mis-mapping.
- Adapter sequences are removed for each library, in order to improve mapping efficiency.
- Sequences shorter than 70 bp are removed.
- Bear in mind that trimming parameters may vary according to the original quality of fastq files, and in some cases, this step can be skipped if no improvement is seen in the alignment step.

```
trim_galore -q 30 --length 70 --paired --fastqc R1.fq.gz R2.fq.gz
```

```
macarw@DESKTOP-Q9UPIUR:/mnt/c/Users/MacaRW/Desktop/Giardia$ trim_galore -q 30 --length 70 --paired --fastqc R1.fq.gz R2.
fq.gz
Multicore support not enabled. Proceeding with single-core trimming.
Path to Cutadapt set as: 'cutadapt' (default)
Cutadapt seems to be working fine (tested command 'cutadapt --version')
Cutadapt version: 2.8
single-core operation.
No quality encoding type selected. Assuming that the data provided uses Sanger encoded Phred scores (default)


AUTO-DETECTING ADAPTER TYPE
===========================
Attempting to auto-detect adapter type from the first 1 million sequences of the first file (>> R1.fq.gz <<)

Found perfect matches for the following adapter sequences:
Adapter type    Count   Sequence        Sequences analysed      Percentage
Illumina        264408  AGATCGGAAGAGC   1000000 26.44
Nextera 0       CTGTCTCTTATA    1000000 0.00
smallRNA        0       TGGAATTCTCGG    1000000 0.00
Using Illumina adapter for trimming (count: 264408). Second best hit was Nextera (count: 0)

Writing report to 'R1.fq.gz_trimming_report.txt'

SUMMARISING RUN PARAMETERS
```

- Use --paired parameter when dealing with paired-end data. Otherwise, remove --paired parameter.
- You should end up with four extra files for each fastq file (e.g. for R1.fq.gz you obtain R1_val_1.fq.gz, R1.fq.gz_trimming_report.txt, R1_val_1_fastqc.html and R1_val_1_fastqc.zip). The .html can be open in a browser of your preference, in order to visualize the quality of trimmed reads. R1_val_1.fq.gz and R2_val_2.fq.gz contain trimmed reads that will be used in the alignment step.

```
macarw@DESKTOP-Q9UPIUR:/mnt/c/Users/MacaRW/Desktop/Giardia$ ls
GCF_000002435.2_UU_WB_2.1_genomic.fna  R1_fastqc.zip         R2.fq.gz_trimming_report.txt  R2_val_2_fastqc.zip
GCF_000002435.2_UU_WB_2.1_genomic.gtf  R1_val_1.fq.gz        R2_fastqc.html                genomeDir
R1.fq.gz                               R1_val_1_fastqc.html  R2_fastqc.zip
R1.fq.gz_trimming_report.txt           R1_val_1_fastqc.zip   R2_val_2.fq.gz
R1_fastqc.html                         R2.fq.gz              R2_val_2_fastqc.html
macarw@DESKTOP-Q9UPIUR:/mnt/c/Users/MacaRW/Desktop/Giardia$ |
```

#### Alignment

- Reads that passed quality criteria are aligned to the latest reference genome using STAR with default parameters except for: (1) --alignIntronMax (maximum intron size) that is set to 1 because with few exceptions, *G. lamblia* genes lack introns, (2) --outFilterMultimapNmax (max number of multiple alignments allowed for a read) that is set to 2 to account for duplicated genes, and (3) --outSAMtype that is set to BAM Unsorted in order to output alignments directly in binary BAM format.
- You can set --outFileNamePrefix to your preferred output files name prefix. We will use Sample1_.
- Remember that you can adjust --runThreadN parameter according to the number of available cores on your server node.

```
STAR --runThreadN 8 --runMode alignReads --genomeDir ./genomeDir --outFilterMultimapNmax 2 --alignIntronMax 1 --outSAMtype BAM Unsorted --outFileNamePrefix Sample1_ --readFilesIn <(gunzip -c R1_val_1.fq.gz) <(gunzip -c R2_val_2.fq.gz)
```

- This will generate the following files:

**Sample1_**Aligned.out.bam → This is the file with the alignment

**Sample1_**Log.final.out

**Sample1_**Log.out

**Sample1_**Log.progress.out

**Sample1_**SJ.out.tab

```
macarw@DESKTOP-Q9UPIUR:/mnt/c/Users/MacaRW/Desktop/Giardia$ STAR --runThreadN 8 --runMode alignReads --genomeDir ./genom
eDir --outFilterMultimapNmax 2 --alignIntronMax 1 --outSAMtype BAM Unsorted --outFileNamePrefix Sample1_ --readFilesIn <
(gunzip -c R1_val_1.fq.gz) <(gunzip -c R2_val_2.fq.gz)
Oct 26 00:43:25 ..... started STAR run
Oct 26 00:43:25 ..... loading genome
Oct 26 00:43:25 ..... started mapping
Oct 26 00:44:57 ..... finished successfully
macarw@DESKTOP-Q9UPIUR:/mnt/c/Users/MacaRW/Desktop/Giardia$ ls
GCF_000002435.2_UU_WB_2.1_genomic.fna  R1_val_1.fq.gz                 R2_fastqc.zip             Sample1_Log.out
GCF_000002435.2_UU_WB_2.1_genomic.gtf  R1_val_1_fastqc.html           R2_val_2.fq.gz            Sample1_Log.progress.out
R1.fq.gz                               R1_val_1_fastqc.zip            R2_val_2_fastqc.html      Sample1_SJ.out.tab
R1.fq.gz_trimming_report.txt           R2.fq.gz                       R2_val_2_fastqc.zip       genomeDir
R1_fastqc.html                         R2.fq.gz_trimming_report.txt   Sample1_Aligned.out.bam
R1_fastqc.zip                          R2_fastqc.html                 Sample1_Log.final.out
macarw@DESKTOP-Q9UPIUR:/mnt/c/Users/MacaRW/Desktop/Giardia$
```

#### Sorting and indexing the alignment

- You will need to install samtools.
- For sorting of .bam file write the following command in your terminal:

```
for file in *.out.bam; do samtools sort $file > ${file/.out.bam/.out.sorted.bam}; done
```

- This will generate a new file: Sample1_Aligned.out.sorted.bam

```
macarw@DESKTOP-Q9UPIUR:/mnt/c/Users/MacaRW/Desktop/Giardia$ for file in *.out.bam; do samtools sort $file > ${file/.out.
bam/.out.sorted.bam}; done
[bam_sort_core] merging from 3 files and 1 in-memory blocks...
macarw@DESKTOP-Q9UPIUR:/mnt/c/Users/MacaRW/Desktop/Giardia$ ls
GCF_000002435.2_UU_WB_2.1_genomic.fna  R1_val_1_fastqc.zip            Sample1_Aligned.out.bam
GCF_000002435.2_UU_WB_2.1_genomic.gtf  R2.fq.gz                       Sample1_Aligned.out.sorted.bam
R1.fq.gz                               R2.fq.gz_trimming_report.txt   Sample1_Log.final.out
R1.fq.gz_trimming_report.txt           R2_fastqc.html                 Sample1_Log.out
R1_fastqc.html                         R2_fastqc.zip                  Sample1_Log.progress.out
R1_fastqc.zip                          R2_val_2.fq.gz                 Sample1_SJ.out.tab
R1_val_1.fq.gz                         R2_val_2_fastqc.html           genomeDir
R1_val_1_fastqc.html                   R2_val_2_fastqc.zip
macarw@DESKTOP-Q9UPIUR:/mnt/c/Users/MacaRW/Desktop/Giardia$
```

- For indexing of the .sorted.bam file write the following command:

```
for file in *_Aligned.out.sorted.bam; do samtools index $file; done
```

- This will generate a new file: Sample1_Aligned.out.sorted.bam.bai

```
macarw@DESKTOP-Q9UPIUR:/mnt/c/Users/MacaRW/Desktop/Giardia$ for file in *_Aligned.out.sorted.bam; do samtools index $fil
e; done
macarw@DESKTOP-Q9UPIUR:/mnt/c/Users/MacaRW/Desktop/Giardia$ ls
GCF_000002435.2_UU_WB_2.1_genomic.fna  R1_val_1_fastqc.zip         Sample1_Aligned.out.bam
GCF_000002435.2_UU_WB_2.1_genomic.gtf  R2.fq.gz                    Sample1_Aligned.out.sorted.bam
R1.fq.gz                               R2.fq.gz_trimming_report.txt  Sample1_Aligned.out.sorted.bam.bai
R1.fq.gz_trimming_report.txt           R2_fastqc.html              Sample1_Log.final.out
R1_fastqc.html                         R2_fastqc.zip               Sample1_Log.out
R1_fastqc.zip                          R2_val_2.fq.gz              Sample1_Log.progress.out
R1_val_1.fq.gz                         R2_val_2_fastqc.html        Sample1_SJ.out.tab
R1_val_1_fastqc.html                   R2_val_2_fastqc.zip         genomeDir
macarw@DESKTOP-Q9UPIUR:/mnt/c/Users/MacaRW/Desktop/Giardia$
```

- The *_Aligned.out.sorted.bam files can be viewed and explored with an interactive tool such as Integrative Genome Viewer (IGV).

**Count number of reads per gene**

- After mapping, reads are assigned to genomic features using FeatureCounts. Multi-overlapping and reads not overlapping with any features in the annotation file are not counted. Since many VSP genes are duplicated, two strategies for counting multimapping reads are used. In strategy UM, multimapping reads are not counted, only unique mapping reads are considered, whereas in strategy 2M, multimapping reads are fractionally counted (each alignment carrying 1/2 count, since the parameter --outFilterMultimapNmax was previously set to 2 in the alignment step).
- You will need to install subread.

```
featureCounts -T 8 -t gene -g gene_id -s 2 -p -M --fraction -a
GCF_000002435.2_UU_WB_2.1_genomic.gtf -o Counts_Sample1_sense_2M
Sample1_Aligned.out.sorted.bam
```

Some considerations:

- You can adjust -T parameter according to the number of available cores on your server node.
- Use -p for paired-end alignments. Remove -p parameter for single-end alignments.
- Use -M --fraction when using 2M strategy. Remove -M --fraction when using UM strategy.

- Use -s 0, -s 1 or -s 2 for unstranded, stranded or reversely-stranded data, respectively. In this example, as the library was reversely-stranded, parameter –s 2 indicates sense counts.
- You can use your preferred output files name in -o. We will use Counts_Sample1_sense_2M.

```
macarw@DESKTOP-Q9UPIUR:/mnt/c/Users/MacaRW/Desktop/Giardia$ featureCounts -T 8 -t gene -g gene_id -s 2 -p -M --fraction
-a GCF_000002435.2_UU_WB_2.1_genomic.gtf -o Counts_Sample1_sense_2M Sample1_Aligned.out.sorted.bam

        ==========     _____ _    _ ____  _____  ______          _____
        =====         / ____| |  | |  _ \|  __ \|  ____|   /\   |  __ \
          =====      | (___ | |  | | |_) | |__) | |__     /  \  | |  | |
            ====      \___ \| |  | |  _ <|  _  /|  __|   / /\ \ | |  | |
              ====    ____) | |__| | |_) | | \ \| |____ / ____ \| |__| |
        ==========   |_____/ \____/|____/|_|  \_\______/_/    \_\_____/
          v2.0.0

//========================== featureCounts setting ===========================\\
||                                                                            ||
||             Input files : 1 BAM file                                       ||
||                           o Sample1_Aligned.out.sorted.bam                 ||
||                                                                            ||
||             Output file : Counts_Sample1_sense_2M                          ||
||                 Summary : Counts_Sample1_sense_2M.summary                  ||
||              Annotation : GCF_000002435.2_UU_WB_2.1_genomic.gtf (GTF)      ||
||      Dir for temp files : ./                                               ||
||                                                                            ||
||                 Threads : 8                                                ||
||                   Level : meta-feature level                               ||
||              Paired-end : yes                                              ||
||      Multimapping reads : counted (fractional)                             ||
|| Multi-overlapping reads : not counted                                      ||
||   Min overlapping bases : 1                                                ||
||                                                                            ||
||          Chimeric reads : counted                                          ||
||        Both ends mapped : not required                                     ||
||                                                                            ||
\\============================================================================//

//================================= Running ==================================\\
||                                                                            ||
|| Load annotation file GCF_000002435.2_UU_WB_2.1_genomic.gtf ...              ||
||    Features : 5391                                                         ||
||    Meta-features : 5388                                                    ||
||    Chromosomes/contigs : 28                                                ||
||                                                                            ||
|| Process BAM file Sample1_Aligned.out.sorted.bam...                         ||
||    Strand specific : reversely stranded                                    ||
||    Paired-end reads are included.                                          ||
||    Total alignments : 3676440                                              ||
||    Successfully assigned alignments : 3293673 (89.6%)                      ||
||    Running time : 0.10 minutes                                             ||
||                                                                            ||
||                                                                            ||
|| Summary of counting results can be found in file "Counts_Sample1_sense_2M  ||
|| .summary"                                                                  ||
||                                                                            ||
```

- You will end up with two new files: Counts_Sample1_sense_2M (with the raw reads per gene) and Counts_Sample1_sense_2M.summary.

```
macarw@DESKTOP-Q9UPIUR:/mnt/c/Users/MacaRW/Desktop/Giardia$ ls
Counts_Sample1_sense_2M                 R1_val_1_fastqc.html           Sample1_Aligned.out.bam
Counts_Sample1_sense_2M.summary         R1_val_1_fastqc.zip            Sample1_Aligned.out.sorted.bam
GCF_000002435.2_UU_WB_2.1_genomic.fna   R2.fq.gz                       Sample1_Aligned.out.sorted.bam.bai
GCF_000002435.2_UU_WB_2.1_genomic.gtf   R2.fq.gz_trimming_report.txt   Sample1_Log.final.out
R1.fq.gz                                R2_fastqc.html                 Sample1_Log.out
R1.fq.gz_trimming_report.txt            R2_fastqc.zip                  Sample1_Log.progress.out
R1_fastqc.html                          R2_val_2.fq.gz                 Sample1_SJ.out.tab
R1_fastqc.zip                           R2_val_2_fastqc.html           genomeDir
R1_val_1.fq.gz                          R2_val_2_fastqc.zip
macarw@DESKTOP-Q9UPIUR:/mnt/c/Users/MacaRW/Desktop/Giardia$ |
```

```
macarw@DESKTOP-Q9UPIUR:/mnt/c/Users/MacaRW/Desktop/Giardia$ head Counts_Sample1_sense_2M
# Program:featureCounts v2.0.0; Command:"featureCounts" "-T" "8" "-t" "gene" "-g" "gene_id" "-s" "2" "-p" "-M" "--fracti
on" "-a" "GCF_000002435.2_UU_WB_2.1_genomic.gtf" "-o" "Counts_Sample1_sense_2M" "Sample1_Aligned.out.sorted.bam"
Geneid  Chr     Start   End     Strand  Length  Sample1_Aligned.out.sorted.bam
GL50803_0061446 NC_051856.1     812     1450    +       639     0
GL50803_0061449 NC_051856.1     3953    4437    +       485     0
GL50803_0061452 NC_051856.1     5462    7795    +       2334    97
GL50803_0061453 NC_051856.1     9417    10928   +       1512    74
GL50803_0061458 NC_051856.1     14753   15862   +       1110    126
GL50803_0061460 NC_051856.1     17070   17288   -       219     53
GL50803_0050370 NC_051856.1     20623   22266   -       1644    201.50
GL50803_00137614        NC_051856.1     23545   25614   +       2070    65
```

#### Convert raw reads to TPM

- Open an R session.
- You will need to install scater and openxlsx libraries.
- Execute the following commands one by one:

```
R
setwd("PATH_TO/Giardia") #Replace PATH_TO with the route to Giardia dir
counts <- read.table(Counts_Sample1_sense_2M, header = TRUE)
head(counts)
library(scater)
TPM <- calculateTPM(x = as.matrix(counts[ , 7]),
                lengths = counts$Length)
TPM <- cbind.data.frame(counts[, 1:2], TPM)
head(TPM)
library(openxlsx)
write.xlsx(TPM, file = "TPM_Sample1_sense_UM.xlsx")
```

- Consider sense and antisense counts for each gene in each library when using stranded data.

#### Software versions

| **Software** | **Version** |
|---|---|
| FastQC | 0.11.9 |
| TrimGalore | 0.6.4_dev |

| | |
|---|---|
| STAR | 2.5.4b |
| samtools | 1.7 |
| FeatureCounts | 2.0.0 |
| R | 4.0.3 |
